# Brain disorder mechanisms and therapeutic targets identified from developmental, cell-specific gene regulation

**DOI:** 10.64898/2026.11.15.766341

**Authors:** Narjes Rohani, Rothem Kovner, Yuting Liu, Antonio Garcia-Guerra, Theodora Markati, Soo-Whee Kim, Yujin Kim, Qi Xu, Emilie M. Wigdor, Jakob Weiss, Joseph L. Lalli, Shan Dong, Anandita Nadkarni, Xuran Wang, Lambertus Klei, Sirisha Pochareddy, Daniel Speckhard-Pasque, Chimmi Dema, Sandra Canosa, Hyesun Cho, Ashley Deveau-French, Gracie Gordon, Michael Griffiths, Karan Jaisingh, Joy Kachko, Pooja Kathail, Mingli Li, Dan Liang, Alicia Ljungdahl, Shaojie Ma, Ruhani Makkar, Nicholas F. Page, Alexa-Nicole Sliby, Menglei Zhang, Xuefang Zhao, Homa Adle-Biassette, Chun Jimmie Ye, Harrison Brand, Michael E. Talkowski, Philip L. De Jager, Donna M. Werling, Matthew J. Girgenti, Carlo Rinaldi, Bernie Devlin, Kathryn Roeder, Nenad Sestan, Stephan J. Sanders

## Abstract

Genetic variation influences brain physiology and disease across cell types and development. We integrated whole-genome sequencing, bulk RNA sequencing, and single-nucleus multiome profiling from 231 human dorsolateral prefrontal cortex donors spanning six postconceptional weeks to 92 years. Across 24 brain-related traits, neurodevelopmental and psychiatric genetic risk was enriched in neuronal transcriptomic and chromatin profiles, whereas neurodegenerative risk was enriched in microglia. Thousands of expression, chromatin accessibility, and splicing quantitative trait loci revealed developmental and cell-type-specific regulatory effects. Colocalization linked multiple sclerosis risk to *AHI1* expression in prenatal neurons and problematic alcohol use to *GPX1* expression in postnatal glia. Leveraging dynamic cis-regulatory elements, we identified therapeutic candidates for haploinsufficient neurodevelopmental disorders, including a *SOX5* enhancer whose CRISPR activation increased transcript and protein abundance. Together, these findings establish a publicly accessible genomic resource for resolving the developmental and cellular contexts of genetic risk and identifying therapeutic targets in the human brain.

**HIGHLIGHTS:**

- **Biology**: Chromatin and expression analyses reveal cell-type and developmental dynamics of brain disorder genes and variants.
- **Atlas**: Cell-type-resolved QTLs reveal dynamic genetic regulation during brain development.
- **Mechanisms**: Prenatal neuronal eQTLs implicate *AHI1* in multiple sclerosis mechanisms.
- **Therapy**: CRISPR activation of an enhancer increases *SOX5* mRNA and protein expression.
- **Resource**: BrainVar website enables interactive exploration of genetic regulation during brain development.

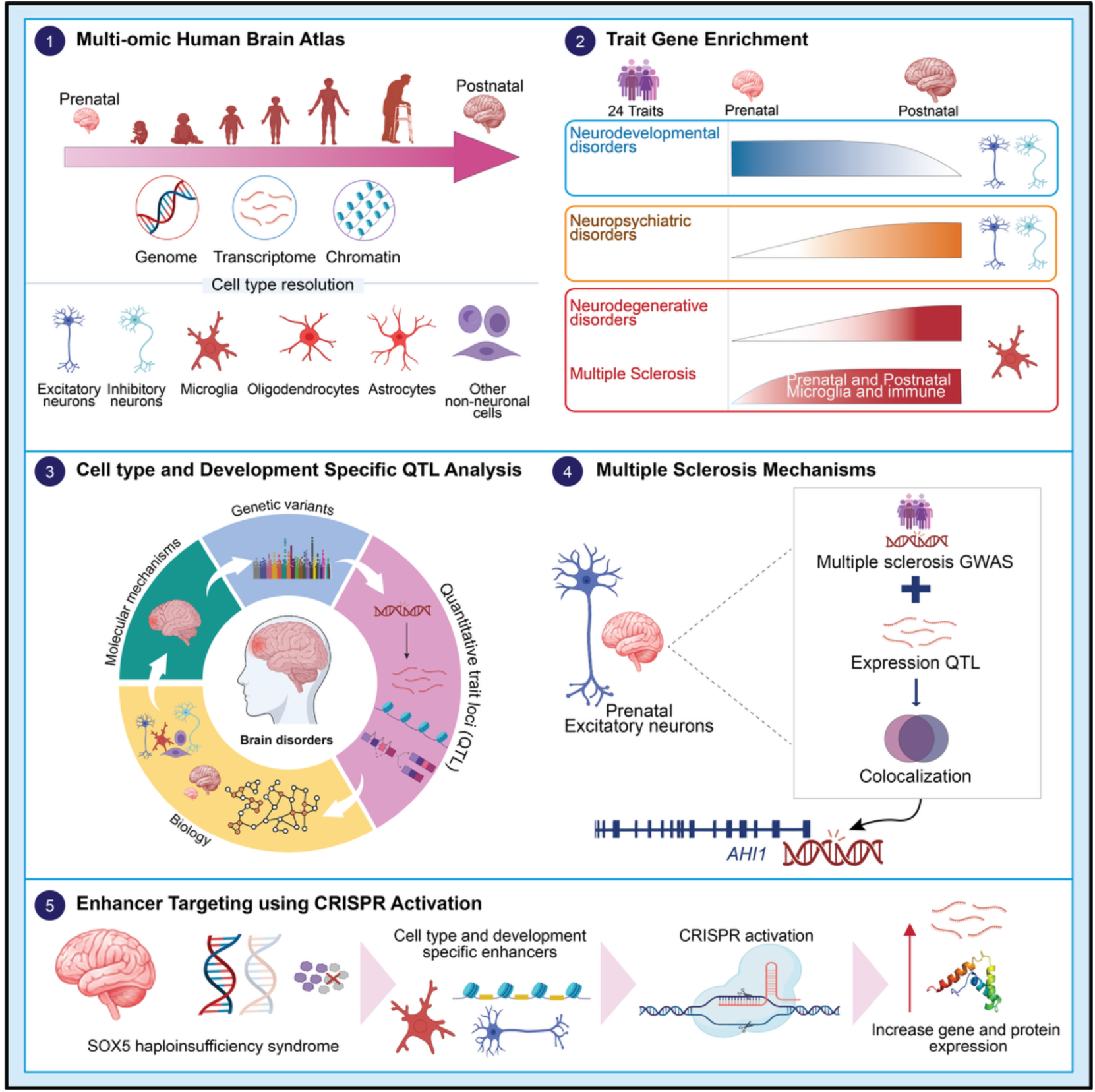

## INTRODUCTION

Genome-wide association studies (GWAS) and sequencing studies have identified thousands of common and rare genetic variants associated with neurodevelopmental, neuropsychiatric, and neurodegenerative disorders.^1–5^ However, most disorder-associated variants reside in noncoding regions of the genome, making it difficult to determine how they influence gene regulation and in which cellular and developmental contexts they act.^6^ Therefore, understanding where and when genetic liability affects molecular programs in the human brain is a central challenge at the cross-roads of neuroscience and human genetics. Recent large-scale single-nuclei and multi-omic studies have begun to define the cellular diversity and regulatory landscapes in the developing human brain,^7–9^ yet the mechanisms connecting brain disorder-associated variation to developmental and cellular gene regulation remain incomplete.

Human cortical development is a highly dynamic process that differs in prenatal and postnatal life. Over time, neurons and glial cells progressively acquire distinct transcriptional and regulatory identities that shape cortical maturation, synaptic function, and circuit organization.^7,10,11^ Genetic variants can perturb these developmental programs at multiple molecular levels, including chromatin accessibility, transcription, and RNA splicing.^7–9^ Increasing evidence suggests that disorder-associated variants act through cell type-predominant regulatory mechanisms and that developmental timing strongly influences their effects.^8,11^ Neurodevelopmental disorders are frequently linked to early corticogenesis and neuronal differentiation;^12^ which occur during prenatal development, whereas psychiatric disorders often involve postnatal synaptic maturation and circuit remodeling.^5,13,14^ In contrast, neurodegenerative disorders commonly implicate glial and inflammatory pathways.^15^

Although recent studies have demonstrated the power of multi-omic approaches for exploring developmental regulatory programs,^8,9,11^ no current atlas integrates whole-genome sequencing (WGS), transcriptomic profiling, and chromatin accessibility with cell type resolution spanning prenatal and postnatal human cortex with diversity in age and sex. Moreover, most existing molecular quantitative trait loci (QTL) studies have been performed in bulk tissue or adult brain samples,^7,16,17^ limiting the ability to resolve how genetic regulation differs across specific cell-types and developmental stages. As a result, developmental-and cell type-predominant expression quantitative trait loci (eQTLs), chromatin accessibility quantitative trait loci (caQTLs), and splicing quantitative trait loci (sQTLs) remain poorly characterized in the human brain.

Here, we present BrainVar, an integrative multi-omic atlas of the human dorsolateral prefrontal cortex across development, spanning early prenatal periods (6 post-conception weeks [PCW]) to late adulthood (92 years). By combining WGS, bulk RNA sequencing, and single-nucleus multiome (snMultiome) profiling, we enable assessment of matched RNA and chromatin accessibility from the same nuclei and analysis of transcriptional and regulatory landscapes over brain cell types and developmental periods. We integrated these data with genetic liability from 24 GWAS/exome datasets for neurodevelopmental, neuropsychiatric, neurodegenerative, and other traits to perform a cross-trait analysis of when and where disorder-associated regulatory programs converge in the human dorsolateral prefrontal cortex (dlPFC). We next mapped cell type-resolved eQTLs, caQTLs, and sQTLs in the human prefrontal cortex throughout development. These analyses revealed extensive developmental and cellular specificity in genetic regulation, with many QTLs showing prenatal- or postnatal-predominant effects. Integration of QTL maps with GWAS signals identified disorder-associated regulatory mechanisms with precise developmental and cellular resolution. Finally, we leveraged these developmental and cell type-resolved atlas to identify cis-regulatory elements capable of modulating haploinsufficient neurodevelopmental disorder (NDD) genes. We used CRISPR activation (CRISPRa) to increase expression of *SOX5,* a haploinsufficient NDD-associated gene, providing proof-of-concept for leveraging developmental regulatory atlases for early-stage translation of therapies. Together, our findings establish a developmental and cell type-resolved framework for interpreting genetic variation in the human brain and we provide a dedicated web resource for these analyses: www.brainvar.org.

## RESULTS

### A comprehensive multi-omic atlas of the human brain over cell types and developmental periods

We generated a multi-omic atlas of the human postmortem frontal cerebral wall (donors from 6-10 PCW) or dorsolateral prefrontal cortex (dlPFC, Brodmann area 46) spanning early prenatal (6 PCW) to late adult (92 years) development; **Figure 1A; Figure S1A**). Joint profiling of WGS, bulk RNA sequencing, and snMultiome (snRNA-seq and snATAC-seq from the same nuclei) yielded cell-level insights into transcription and chromatin accessibility and how these are influenced by genetic variation and developmental age. Further details on the snMultiome data are provided in our companion study [Liu et al., 2026], which examines gene expression trends across human brain development without a specific focus on brain disorders.

**Figure 1.**
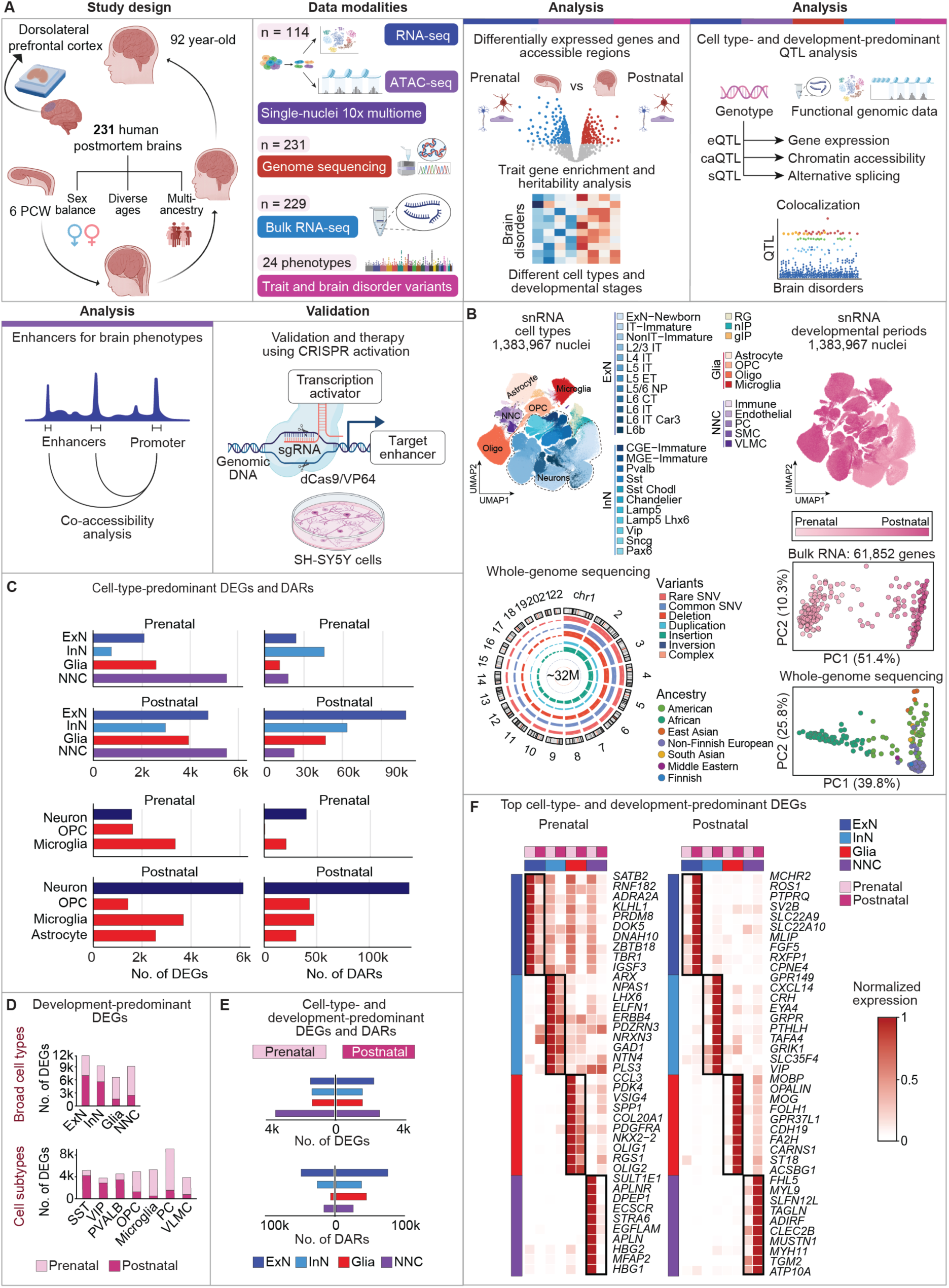
A multi-omic atlas of the human dorsolateral prefrontal cortex across development. **(A)** Study design and cohort composition. Left: Postmortem dlPFC tissue was obtained from 231 donors spanning 6 PCW to 92 years of age (132 prenatal and 99 postnatal). See also **Figure S1A** for the number of donors in each period. Middle: Multi-omic profiling included WGS, bulk RNA sequencing, and snMultiome. The number of donors per modality and the number of traits analyzed are indicated (see also **Table S1 and S4**). Right: Downstream analyses included differential expression analysis (cell markers), differential chromatin accessibility analysis, SNP-based heritability enrichment, QTL mapping (eQTL, sQTL, and caQTL), co-accessibility analysis and enhancer discovery, and CRISPR activation experiment. Colors above each analysis denote the data modality used for that analysis. **(B)** Cellular, developmental, and genomic landscape of the human dlPFC. Top left: RNA UMAP embedding of the high-quality nuclei derived from 114 donors out of 118 attempted, colored by annotated cell types (see **Table S3)**. Top right: RNA UMAP colored by developmental periods (Periods 3–15), Down left: Distribution of genetic variant classes identified by WGS (262 donors attempted; 231 post-QC). Down right: PCA of bulk RNA-seq data (240 donors attempted; 229 post-QC demonstrating developmental periods and principal component analysis of WGS showing ancestry diversity. **(C)** Cell type-predominant transcriptional and regulatory programs within prenatal and postnatal. Bar plots show the number of markers calculated by Wilcoxon rank sum test (DEGs; adjusted *p* < 0.05, average log₂ fold change > 0.5 used for all analysis) and DARs identified across broad cell types at each developmental stage (full list of results in **Data S1**). **(D)** Developmentally dynamic gene expression across cell types. Numbers of prenatal-and postnatal-pre-dominant DEGs (developmental stage markers) identified by direct comparison of prenatal against postna-tal within each broad cell type and cell subtype (full list of results in **Data S2**). See also **Figure S1D** for development-predominant DARs. **(E)** Joint developmental and cell type DEGs and DARs, highlighting genes and regulatory regions upregulated in specific stage–cell-type combinations (full list of results in **Data S3**). **(F)** Heatmap of top development- and cell-type-predominant protein-coding DEGs ranked by adjusted *p* value and log_2_ fold change. The normalized expression shows min-max scaled pseudobulk average expression across prenatal and postnatal stages for each cell type (full list of results in **Data S3**). **Abbreviations:** caQTLs, chromatin accessibility quantitative trait loci; DARs, differentially accessible regions; DEGs, differentially expressed genes; dlPFC, dorsolateral prefrontal cortex; eQTLs, expression quantitative trait loci; PCA, principal component analysis; PCW, post-conception weeks; RG, radial glia; SMC, smooth muscle cells; snMultiome, single-nucleus multiome; sQTLs, splicing quantitative trait loci; SST, somatostatin-expressing interneurons; UMAP, uniform manifold approximation and projection; VIP, vasoactive intestinal peptide-expressing interneurons; VLMC, vascular and leptomeningeal cells

High-quality WGS and at least one functional genomic modality were generated from 231 donors (out of 263 donors attempted), including 132 prenatal and 99 postnatal donors (**Figure 1A; Figure S1A; Table S1**). WGS (231 donors) yielded over 32 million high-quality variants including single-nucleotide variants, insertions/deletions, and structural variants (**Figure 1B**). The cohort had similar representation by sex (59% male, 41% female) and diverse ancestral backgrounds, including Non-Finnish European (47%), African (29%), American (20%), South Asian (2%), East Asian (2%), Finnish (0.4%), and Middle Eastern (0.4%) individuals (**Figure 1B**). Bulk RNA-seq from 229 donors provided data on 61,852 genes, with clear transcriptional differences between prenatal and postnatal stages (**Figure 1B**).

High-quality snMultiome data were achieved for 114 donors. These data yielded high-quality data for 1,383,967 nuclei for RNA and 904,418 nuclei for ATAC; 715,035 nuclei had both modalities. (**Figure 1B; Table S2; Table S3**). Clustering (**Figure 1B; Figure S1C**) resolved all major cellular lineages found in the cortex, including excitatory (ExN) and inhibitory (InN) neurons, neuronal progenitors, glial populations (astrocytes, oligodendrocytes, oligodendrocyte precursor cells [OPCs], and microglia), vascular, and immune cells. Some clusters were composed of primarily of prenatal (e.g., excitatory neuron, ExN-Newborn) or postnatal (e.g., oligodendrocytes) nuclei, while others (e.g., L6b) exhibited gradients across development within the cluster, indicating progressive transcriptional remodelling within a common cellular identity (**Figure 1B**). Consistent with known developmental dynamics,^18^ neuronal proportions declined between the prenatal and postnatal stage as glial populations expanded postnatally (**Figure S1B**). The dataset can be explored on UCSC cell browser (brainvar.cells.ucsc.edu), UCSC genome browser (https://genome-test.gi.ucsc.edu/cgi-bin/hgTrackUi?db=hg38&g=singleCellSignalsPeaks) and a dedicated interactive web resource (www.brainvar.org/).

### Widespread postnatal expansion of cell type-predominant transcriptional and regulatory programs

Given the dramatic late prenatal changes (**Figure 1B**), we defined cell type-prominent differentially expressed genes (DEGs) and differentially accessible regions (DARs) separately within prenatal (6-24 PCW) and postnatal (birth to 92 years of age) stages (adjusted *p* < 0.05; mean log₂FC > 0.5). Across four broad cell types (ExN, InN, Glia, NNC [non-neuronal cells]), we identified 11,431 prenatal DEGs and 163,143 prenatal DARs, and 17,198 postnatal DEGs and 250,441 postnatal DARs (**Figure 1C; Data S1**). Most cell types had more cell type-predominant DEGs and DARs postnatally than prenatally; this was especially marked for both excitatory and inhibitory neurons (**Figure 1C**). Excitatory neurons, in particular, showed a substantial postnatal expansion of regulatory elements (prenatal: 27,529 DARs; postnatal: 108,728 DARs, **Figure 1C**). Throughout this study, we use the term *predominant* rather than *specific*, as a lack of statistical significance or a smaller effect size in another developmental stage or cell type does not necessarily indicate the absence of an underlying regulatory effect.

### Developmental-predominant transcriptional signatures distinguish neuronal and glial lineages

In bulk tissue, a dramatic shift in gene expression is observed between the mid-fetal development and infancy (**Figure 1B**), named the late-fetal transition.^7,18,19^ Expansion of glial cell populations explains much of this shift (**Figure S1B**); however, the increase of DEGs across development shows that transcriptional changes within cell types also contribute. To define developmental-predominant programs within each cell type, we directly compared prenatal and postnatal nuclei within four broad cell types and seven cell subtypes. In general, neurons had more postnatal-predominant DEGs driven by increasing expression from the late-fetal transition across childhood (**Figure S1F**), while non-neurons had more prenatal-predominant DEGs (**Figure 1D**). Excitatory neurons exhibited the largest number of developmentally dynamic DEGs (n = 11,751; prenatal-predominant: 4,730; postnatal-predominant: 7,021), followed by inhibitory neurons (n = 9,376; prenatal: 3,801; postnatal: 5,575) (**Figure 1D; Data S2**). These findings suggest substantial postnatal maturation of neurons, likely reflecting ongoing synaptic and circuit development. Among non-neuronal populations, OPCs (n = 5,177 DEGs; mostly prenatal-predominant) and microglia (n = 5,540 DEGs; mostly prenatal-predominant) displayed strong developmental signatures. In the postnatal brain, ongoing neuronal diversification contrasts with relatively static glial lineages at the transcriptomic level. We investigated these developmental trajectories in greater depth in our companion study [Liu et al., 2026].

Considering chromatin accessibility, all four broad cell types exhibited more developmental DARs postnatally than prenatally (**Figure 1E**). The most developmental DARs were observed in excitatory neurons (prenatal: 76,608 DARs; postnatal: 133,399 DARs) and inhibitory neurons (prenatal: 45,380 DARs; postnatal: 98,816 DARs) (**Figure 1E; Figure S1D; Data S2)**.

### Cell identity and developmental timing jointly specify cortical molecular states

Given that some genes are cell-type-predominant (**Figure 1C**) while others are developmentally-predominant within cell types (**Figure 1D**), we assessed whether there are genes and chromatin accessible regions that are both cell-type and developmentally predominant (**Figure 1E**). Such a joint analysis can identify unique and complex transcriptional and regulatory programs (**Figure 1E; Data S3**). To achieve this, we contrasted each of the four broad cell-types and two developmental stages against the seven other combinations. Prenatal (n = 3,409) and postnatal (n = 2,500) non-neuronal cells, as well as postnatal excitatory neurons (n = 2,258), exhibited the greatest numbers of cell-type- and development-predominant DEGs. At the regulatory level, excitatory neurons displayed the most differentially accessible chromatin regions in both prenatal (n = 52,279 DARs) and postnatal (n = 93,982) stages, underscoring their pronounced regulatory complexity across development. These cell-type- and development-predominant DEGs showed significantly lower LOEUF (Loss-of-function Observed/Expected Upper-bound Fraction) scores than background genes in keeping with intolerance to gene dosage variation (**Figure S1F**; **Table S6**). Pathway enrichment analyses (**Table S5**) of the cell-type-, development-, and cell-type-and-development-predominant DEGs showed neurons transitioning from prenatal states enriched for protein synthesis to postnatal states enriched for synaptic communication.

To identify regulatory factors driving these transitions, we performed motif enrichment (**Table S4**; **Figure S1G**) of DARs. Known motifs of cell type identify (e.g., NeuroG2, OLIG2) and development (e.g., neurons transitioning from prenatal Homeobox and Forkhead motifs, to ETS-, AP-1/bZIP-, KLF-, and MEF2-family motifs) were observed. Glial cells exhibited a distinct developmental pattern, including significant postnatal enrichment of *SOX*-family motifs.

Among the top cell-type- and development-predominant DEGs (**Figure 1F**), several genes with established links to neurodevelopmental disorders exhibited striking temporal and cell type specificity. *GAD1*, associated with autosomal recessive developmental epileptic encephalopathy,^20^ was highly active in prenatal inhibitory neurons, consistent with its central role in GABA synthesis and inhibitory circuit development during early brain maturation. Conversely, *TBR1*, a transcription factor critical for corticogenesis and in which haploinsufficiency is associated with autism spectrum disorder (ASD), and intellectual disability,^2^ showed strong prenatal predominancy in excitatory neurons, aligning with its established function in early cortical neuron specification.

Together, these results demonstrate that cortical development is characterized by progressive, lineage-specific expansion of transcriptional and regulatory programs. Neurons in particular exhibit extensive postnatal diversity in transcription and chromatin accessibility.

### Cross-disorder cell-type and developmental stage convergent biology

Having categorized high-level differential cell type gene expression and chromatin accessibility across dlPFC development, we integrated gene lists associated with human brain disorders (**Figure S2A**, **Table S7**) and traits to gain insights into the underlying neurobiology. Using gene set enrichment analysis (GSEA), we observed marked patterns across cell-types and development (**Figure 2A, Figure S2B, Table S8**). At a high level, genes associated with neurodevelopmental and psychiatric disorders were enriched in neurons (e.g., *MYT1L* associated with ASD and neurodevelopmental delay),^21–23^ while genes associated with multiple sclerosis (MS) and Alzheimer’s disease (ALZ)^1^ were enriched in microglia (e.g., *MERTK* associated with MS);^24,25^ such patterns are consistent with prior findings.^7,19^ Genes associated with both neurodevelopmental and psychiatric disorders (except major depressive disorder [MDD] and post-traumaric stress disorder [PTSD]) were more likely to be enriched in prenatal neurons than postnatal neurons; however, the effect sizes between the two developmental stages were rarely dramatic.^26,27^

**Figure 2.**
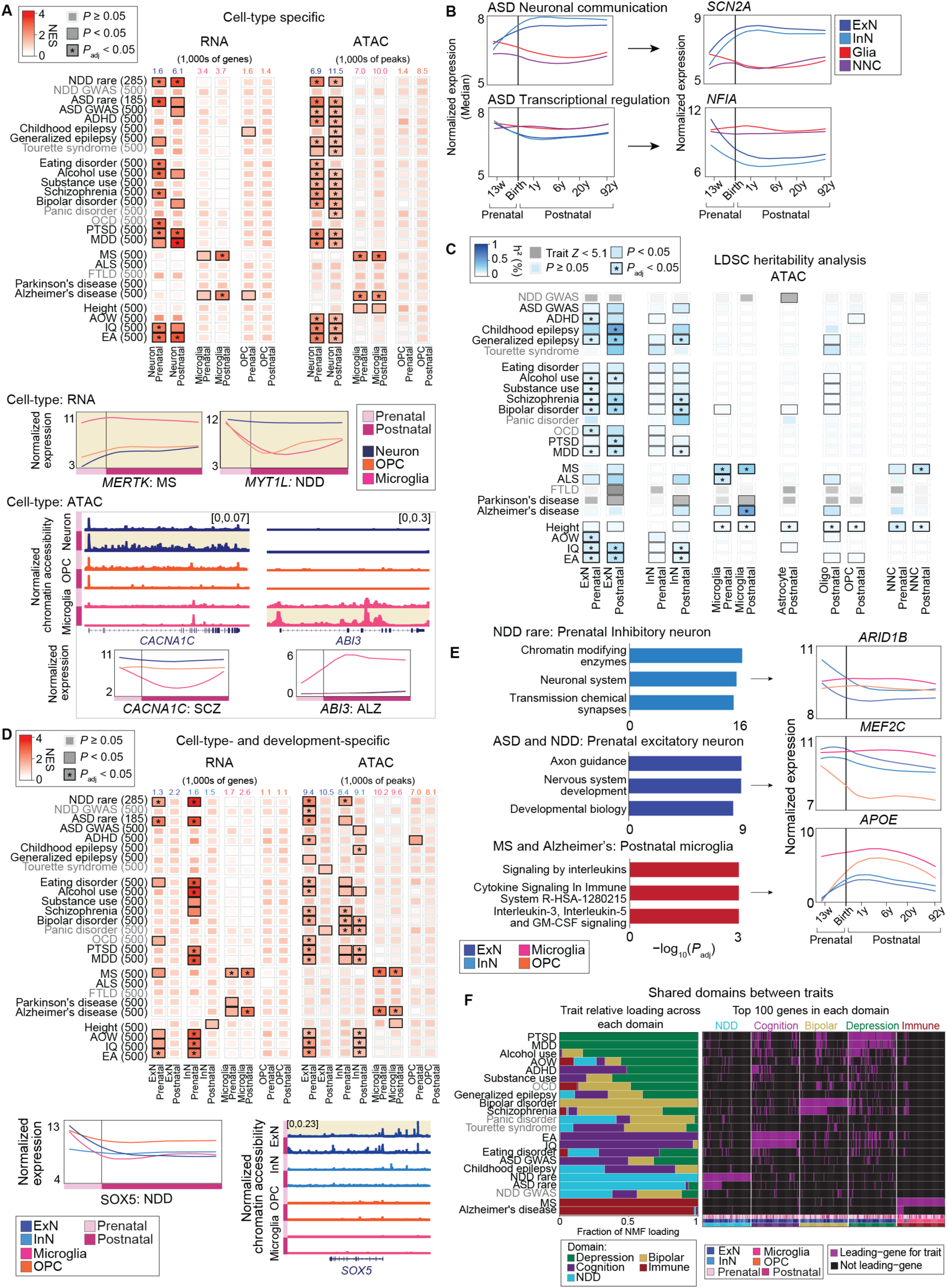
Cell-type and developmental enrichment of genetic liability of brain disorders. **(A)** Up: Gene set enrichment analysis of brain disorder–associated genes across major brain cell types stratified by developmental stage using RNA and ATAC (using gene activity score). Brain disorder variants were mapped to genes using MAGMA (**DATA S5**), and the top 500 genes with MAGMA *p* < 0.05 were included in the enrichment analysis. The number next to each trait indicates the number of genes used for enrichment, and the number above each cell type indicates the number of significant (adjusted *p* < 0.05, log_2_FC>0.5) upregulated genes in that cell population. All genes were ranked by fold change and adjusted *p* value and analyzed using fgsea function in R (**Table S8**), with only positive enrichments shown. Traits shown in grey represent those with low-powered GWAS data (**Figure S2A**). Down: Representative examples of enriched disorder-associated genes illustrating cell-type-predominant smoothed gene expression (log2 [CPM +1]) and chromatin accessibility (ReadsInTSS normalized) patterns across development and cell-type. **(B)** Expression patterns of ASD-associated genes grouped by functional categories, including transcription regulator and neuronal communication genes. Expression values represent the median log₂-normalized CPM+1 averaged over genes within each developmental period and cell-type (see also **Figure S2F**, **Table S9**). **(C)** Partitioned heritability analysis^30^ using ATAC-defined regulatory regions (see **Table S8, S11**) for brain cell types and developmental stages. Asterisks indicate statistical significance (*p* < 0.05; **FDR < 0.05). Traits shown in grey represent those with low-powered GWAS data (**Figure S2A**). The results for traits with *Z* < 5.1 also were shown in grey (**Table S11**). **(D)** Up: GSEA of trait-associated genes using markers jointly defined by both cell type and developmental stage (Figure 1E) derived from RNA and ATAC (based on gene activity score) datasets calculated by fgsea function in R (**Table S8**). Traits shown in grey represent those with low-powered GWAS data (**Figure S2A**). Down: Representative log₂-normalized CPM+1 expression and chromatin accessibility (ReadsInTSS normalized) patterns of disorder-associated genes across cell types and development. **(E)** Right: Reactome pathway enrichment analysis of NDD genes enriched in prenatal inhibitory neurons, brain disorders in prenatal excitatory neurons, and MS and Alzheimer’s disease in postnatal microglia (full list of pathways in **Table S12**). Left: Expression (log₂-normalized CPM+1) of examples of disorder-associated genes enriched in each pathway. **(F)** Similar biological domains across brain disorders. Left: relative contributions of biological domains across disorders, shown as the fraction of NMF loading and grouped into biological domains including depression, bipolar, cognition, neurodevelopment/NDD, and immune. Right: heatmap showing the top 100 genes for each biological domain. Purple indicates genes identified as leading genes for a trait, whereas black indicates non-leading genes. Annotation tracks below the heatmap indicate cell-type, development, and the corresponding biological domain (see also **Table S13**). **Abbreviations:** ASD, autism spectrum disorder; CPM, counts per million; FDR, false discovery rate; GSEA, gene set enrichment analysis; MAGMA, multi-marker analysis of genomic annotation; MS, multiple sclerosis; NDD, neurodevelopmental disorder; NMF, non-negative matrix factorization; TSS, transcription start site.

*En masse*, ASD-associated genes identified by exome sequencing^2^ were enriched in prenatal neurons, consistent with prior findings.^21^ We further explored this relationship by sub-setting ASD-associated genes into two functional roles: transcriptional regulation^23,28,29^ and neuronal communication (**Figure 2B, Table S9**). Transcriptional regulation^23,28,29^ genes (e.g., *NFIA*) showed the highest expression during prenatal stage in neurons, with a decrease in expression to plateau after infancy. In contrast, expression of neuronal communication genes (e.g., *SCN2A*) increased during prenatal development to peak around infancy and plateau at this high level. Similar results were observed for NDD (**Figure S2F**). Despite these opposing patterns of expression between transcriptional regulation and neuronal communication genes, substantive phenotypic differences between patients affected by the two functional groups have not been reported. This may reflect the contribution of common variation in a ‘dual hit’ model or that both functional groups act via a common mechanism. These results raise the prospect of ongoing postnatal vulnerability in ASD and NDD, a potential target for intervention.

Given the distinct trajectories based on gene function in ASD and NDD, we assessed if similar patterns were observed in two well-powered psychiatric GWASs (**Figure S2A**): MDD and schizophrenia (SCZ) (**Figure S2F, Table S10**). MDD-associated genes showed a similar pattern to NDD and ASD (**Figure S2F, Figure 2B**). However, schizophrenia genes were marked by relatively high expression in non-neuronal cells across both transcriptional-regulation and neuronal-communication genes. This result was also observed in 695 rare variant SCZ genes (SCHEMA, false discovery rate [FDR] < 0.05) and exemplified in genes with the strongest evidence such as *SETD1A* and *FURIN* (**Figure S2F**). In keeping with this observation, enrichment for SCZ genes was observed for prenatal developmental-predominant genes in vascular leptomeningeal cells (VLMCs) but not other psychiatric disorders (**Figure S2C**), suggesting a vascular vulnerability specific to SCZ.

Enrichment patterns inferred from chromatin accessibility (linked to genes using gene activity score) were largely concordant with transcriptional signals; snATAC was generally more sensitive but yielded less dramatic effect sizes (**Figure 2A, Figure S2B**). Little variability was seen between prenatal and postnatal enrichment for most disorders, including MS and ALZ that were enriched in microglia throughout development. For some genes, however, snATAC provided extra insights. For example, *CACNA1C*, a gene associated with SCZ, bipolar disorder (BIP), MDD, and NDD, was most highly expressed in neurons throughout development; however, at the level of chromatin, the gene was more accessible in the postnatal period (**Figure 2A**). For other genes, snATAC showed consistent findings between the two modalities, such as *ABI3*, an ALZ-associated gene mostly highly expressed in microglia with both accessibility and expression enriched in postnatal microglia.

Extending the analysis to finer grained cellular subtypes resulted in a largely underpowered analysis at the level of gene expression (**Figure S2B**), except for MS, enriched in microglia, pericytes, endothelial and immune cells and ALZ, enriched in microglia and immune cells. However, the chromatin accessibility analysis was sufficiently sensitive to identify enrichment in multiple postnatal excitatory and inhibitory neurons across most neurodevelopmental and psychiatric phenotypes. This result supports the notion that multiple neuronal cell types might be involved in these disorders and that analysis of regulatory regions could provide greater resolution for such enrichment analyses (**Figure S2B**). For neurodegenerative disorders, the chromatin findings largely replicated those seen with gene expression (**Figure S2B; Table S8**).

### Partitioned heritability further implicates cell-types and developmental stages in brain disorders

Polygenic scores from GWAS provide a largely orthogonal method to assess cell-type and developmental enrichment by assessing variants within chromatin accessibility peaks directly, rather than linking genes to chromatin peaks using gene activity score. Such an analysis of partitioned heritability within ATAC-defined regulatory regions further confirmed the enrichment patterns observed via RNA and ATAC enrichment (**Figure 2C, Table S8, S11**). Genetic liability for most neurodevelopmental and psychiatric disorders was strongly enriched in neurons, especially post-natal neurons, whereas MS and ALZ showed the strongest enrichment in microglia. As with prior analyses, MS demonstrated a mixed developmental profile, with enrichment strongest at postnatal stages, but also present prenatally. Surprisingly, amyotrophic lateral sclerosis (ALS or motor neuron disease [MND]) GWAS data were enriched in prenatal microglia only.

### Cell-type- and development-predominant markers in brain disorders

Following the logic that genes and peaks with the most restricted patterns across cell type and development provide the greatest insight into these dimensions, we restricted the enrichment analyses to the cell-type- and development-predominant DEGs/DARs identified in **Figure 1D** (**Data S3; Figure 2D; Table S8**). In general, similar cell types were identified, with genes associated with neurodevelopmental and psychiatric traits/disorders enriched in neurons, and neurodegenerative disorders (MS, ALZ) enriched in microglia (**Figure 2D**; **Figure S2B; S2C**). Despite observing higher numbers of cell-type- and development-predominant DEGs in postnatal neurons than prenatal neurons, neurodevelopmental/psychiatric enrichment patterns were strongest in prenatal neurons, especially inhibitory neurons, including ASD, NDD, eating disorder, problematic alcohol use (PAU), PTSD, MDD, age of walking (AOW), IQ, and educational attainment (EA). As before, enrichment in ATAC-seq data were largely consistent with RNA, but more sensitive and more likely to capture a postnatal inhibitory neuron contribution (e.g., BIP, PTSD, MDD, IQ). Complex developmental patterns were observed within genes, e.g., several peaks of chromatin accessibility in the NDD-associated gene *SOX5* were most prominent in prenatal excitatory neurons (**Figure 2D**).

Pathway analysis (**Figure 2E; Table S12**) of the NDD genes with prenatal inhibitory neurons upregulation, resulted by trait-gene enrichment analysis (**Figure 2D**), implicated transcription regulation (e.g., chromatin modifying enzymes) and neuronal communication (e.g., transmission across chemical synapses), e.g., *ARID1B*, a transcriptional regulator associated with ASD/NDD, showed elevated expression in prenatal inhibitory neurons (**Figure 2E**). Protein-protein interaction network analysis of enriched NDD genes in prenatal inhibitory neurons further revealed convergence of NDD-associated genes on chromatin remodelling pathways (**Figure S2D**), with a highly interconnected network centered on components of the SWI/SNF (BAF) chromatin remodelling complex (**Figure S2D; Figure S2E; Table S12**), which has critical roles in brain development,^31^ including *ARID1B*. In prenatal excitatory neurons, transcriptional regulation pathways predominated, e.g., *MEF2C*, another ASD/NDD-associated transcriptional regulator (**Figure 2E**). MS and ALZ genes in postnatal microglia were enriched for immune and inflammatory pathways, e.g., *APOE*, in which risk alleles are estimated to contribute to over half of Alzheimer’s disease cases^32^ was most highly expressed in postnatal microglia.

### Cross-trait analysis reveals five biological domains

We observe similar cross-disorder patterns of enrichment using three orthogonal methods: RNA, ATAC, linkage disequilibrium score regression (LDSC) (**Figure 2A**, **C**, **D**). Given the similar patterns, we considered whether cell/development enrichment patterns could identify related biological vulnerability. Focusing on the union of genes per disorder that contributed to significant RNA/ATAC enrichment (**Figure 2A**, **D**) we applied non-negative matrix factorization (NMF), which highlighted five stable domains (**Figure 2F; Figure S2H**). Three of these domains (dubbed NDD, Cognition, Depression for interpretability) shared similar gene ontology (GO) terms relating to gene regulation, neuronal development, and synaptic communication (**Table S13**) and were highly expressed in both prenatal and postnatal neurons despite little overlap of specific genes. All three were enriched for genes with low LOEUF scores (**Figure S2G**), suggesting gene dosage sensitivity, with the NDD domain highly enriched and driven by rare variant gene discovery. The similar named “Bipolar domain” captured a highly heterogenous group of genes expressed across major cell types with high LOEUF scores while the Immune domain captured a distinct set of genes highly expressed in non-neuronal cells with high LOEUF scores (**Figure S2G; Table S13**). Of note, a few genes contributed substantially across four domains, potentially representing cross-domain vulnerability (*FOXP1*, *HIST1H2BO*, *HIST1H3J*, *RBFOX1*, *KANSL1*, *SORCS3*).

### Molecular quantitative trait loci (QTLs) across development and cell type

Integrating WGS data enables the detection of genetic variation impacting gene regulation and expression in the human dlPFC, including quantitative trait loci (QTLs), i.e., genetic variants, that alter gene expression (eQTLs), chromatin accessibility (caQTLs), or splicing (sQTLs). Since datasets with more samples are more sensitive for QTL discovery, we focus on QTLs with developmental (prenatal and postnatal) and cell type (ExN, InN, Glia, and non-neuronal cells [NNC]) variability. Visualization of eQTLs by gene, developmental stage, and cell type are available on the BrainVar website (www.brainvar.org/).

### Dorsolateral prefrontal cortex expression QTLs with cell-type and developmental stage resolution

eQTLs were detected using Bayesian model-based integration of bulk and single-cell data Bayesian model-based integration of bulk and single-cell data (bMIND)^33^ (**Figure 3A; Figure S3A; Table S14**) to define 196,507 unique eQTLs, corresponding to 158,032 unique eQTL variants (Expression quantitative trait locus variants [eVariants]) and 5,950 unique eQTL target genes (eGenes) across developmental stages and cell types (**Figure 3A**). As expected for cis-regulatory effects, eQTL variants were strongly enriched near the transcription start site of the target gene across all cell types and developmental stages (**Figure 3A; Figure S3E**). The majority of eQTLs (62%) were in more than one cell type and developmental stage, though many were specific to cell type, developmental stage, or cell-type and developmental stage (**Figure S3B; Table S15**).

**Figure 3.**
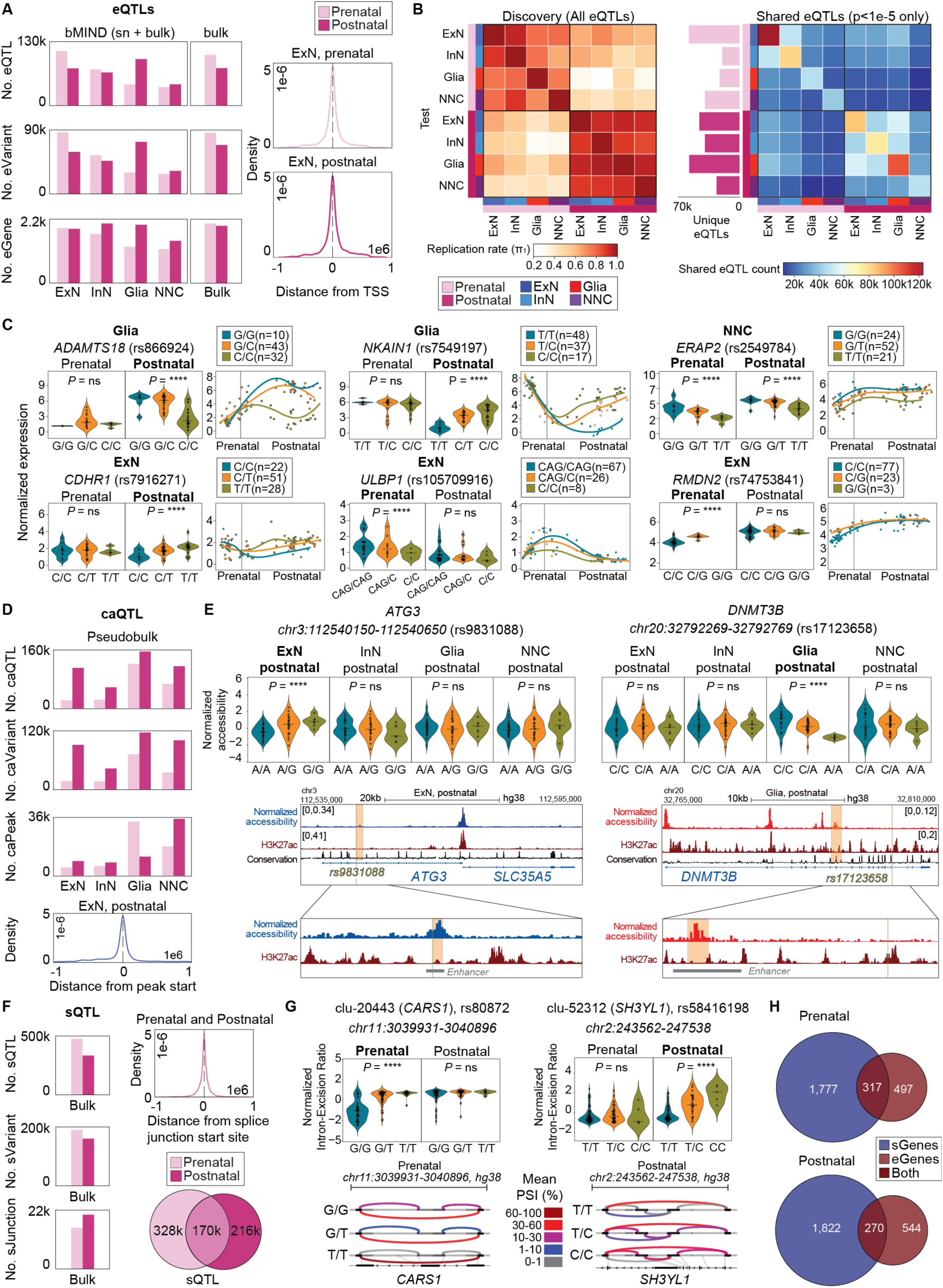
QTL analyses with cell-type and developmental resolution in the human prefrontal cortex. **(A)** Left: Number of significant cis-eQTLs (*p* < 1e-5), eGenes, and eVariants identified across each cell type and developmental stage, and bulk RNA sequencing calculated by TensorQTL pipeline. Right: Distribution of distances between eVariants and TSS in prenatal and postnatal excitatory neurons. The x-axis shows the distance (±1 Mb) relative to the TSS. See also **Figure S3E**. **(B)** Left: π1 reproducibility analysis comparing all bMIND-derived eQTLs across cell types and developmental stages. Stronger color intensity indicates higher reproducibility between datasets. Right: Number of shared and unique significant eQTLs (*p* < 1e-5) identified across developmental stages within each cell type. **(C)** Examples of cell-type and/or development specific eQTLs for disorder-associated genes shown as raw (non-covariate-adjusted) log2CPM+1 expression stratified by genotype. Genotypes are shown as homozygous reference (blue), heterozygous (yellow), and homozygous alternate (green). The *P* shown on plot is nominal significant score calculated by TensorQTL pipeline. Multiple variants showed genotype-dependent expression effects that differed across developmental stages and cell types, including *ADAMTS18* (functions related to metabolism and blood-brain-barrier gene; rs866924), *NKAIN1* (alcohol dependence and cognitive dysfunction gene; rs7549197), *ERAP2* (neuroinflammation and migraine gene; rs2549784), *CDHR1* (inherited retinal dystrophies gene; rs7916271), *ULBP1* (neurodegenerative diseases and depression; rs10579916), and *RMDN2* (ALZ gene; rs74753841). See also **Figure S3D**. **(D)** Top: Number of significant cis-caQTLs, caVariants, and caPeaks identified across each cell type and developmental stage using pseudobulk ATAC-seq profiles. Bottom: Distribution of distances between caQTL variants and start sites of regions in postnatal excitatory neurons. The x-axis shows the distance (±1 Mb) relative to the start sites of regions. **(E)** Representative example of a cell-type- and developmental-stage-predominant caQTL. Violin plots show quantile-normalized chromatin accessibility across donor pseudobulk profiles. UCSC tracks show chromatin accessibility (ReadsInTSS normalized) for the region. H3K27ac is ChIP-seq on dlPFC tissue female adult^36^ (89 years). Conservation score shows phyloP conservation visualized using UCSC genome browser. Enhancers IDs: GH03J112539 and GH20J032791.^37^ **(F)** Left: Number of significant cis-sQTLs, sVariants, and sJunctions identified in prenatal and postnatal stages. Top right: Distribution of distances between sQTL variants and the start sites of splicing junctions for prenatal and postnatal datasets. The x-axis shows the distance (±1 Mb) relative to the start site of the splicing junction. Bottom right: Overlap of significant sQTLs (*p* < 1e-5) between prenatal and postnatal stages showing number of shared and stage-predominant sQTLs. **(G)** Representative examples of sQTLs identified across development. Sashimi plots generated using the LeafCutter/leafviz package show genotype-associated differences as normalized percent spliced-in (PSI) values stratified by genotype. Violin plots show intron excision ratio stratified by genotype. Exons are shown as boxes along the genomic axis; black boxes denote exonic regions participating in the displayed intron cluster, whereas gray boxes indicate additional annotated exonic regions of the gene. The thin horizontal line represents the genomic interval. In the bottom panel, the gene model is displayed, and gray connector lines link splice sites in the intron cluster to their corresponding positions in the gene annotation. Differences in arc color and abundance across genotype groups indicate genotype-associated changes in splice-junction usage and alternative splicing patterns. **(H)** Overlap between sGenes and eGenes showing the number of shared and unique genes regulated at the splicing and expression modalities. All genomic coordinates are based on the hg38 reference genome assembly. See also **Figure S3**. **Abbreviations:** ALZ, Alzheimer’s disease; bMIND, Bayesian model for inferring cell-type-specific gene expression; caQTLs, CPM, counts per million; dlPFC, dorsolateral prefrontal cortex; eQTLs, expression quantitative trait loci; PSI, percent spliced in; sQTLs, splicing quantitative trait loci; TSS, transcription start site.

Cross-context comparison showed that eQTL predominancy was shaped more strongly by developmental stage than by cell type (**Figure 3B**). For genome-wide significant eQTLs (*p* < 1x10^-5^), 74% showed prenatal- or postnatal-predominant patterns while 26% were shared across development stages. The largest numbers of unique genome-wide significant eQTLs were detected in prenatal excitatory neurons and postnatal glia (**Figure 3B**).

Representative loci illustrate the developmental and cellular predominancy of eQTL effects (**Figure 3C**). The same variant could have little or no effect in one developmental period or cell type but a robust effect in another. For example, rs7549197 acted as an eQTL for *NKAIN1* in postnatal glia (*p* = 3.9x10^-5^, effect size = +4.0) but not prenatal glia. Other variants, such as rs2549784 in *ERAP2*, act as an eQTL across cell types and developmental stages, but with wide variation in the effect size (excitatory neuron prenatal/postnatal effect size: -12.1/-11.9, inhibitory neuron: - 1.5/-4.9, Glia: -0.3/-16.0, non-neuronal cells: -0.5/-2.2).

To systematically classify these regulatory patterns, we grouped significant eQTLs into five categories: prenatal-predominant, prenatal-trending, constant, postnatal-predominant, and postnatal-trending (**Figure S3B; Table S15**). We validated BrainVar eQTLs by comparing their effect sizes with previously reported eQTL datasets from bulk tissue^9,16,17,34^ (**Figure S3C**). These datasets with larger sample sizes achieve higher QTL discovery, however, our data reveals unrecognized developmental and cell type resolution.

### Dorsolateral prefrontal cortex chromatin accessibility QTLs with cell type and developmental stage resolution

caQTLs were identified from pseudobulk single-nuclei data alone across prenatal and postnatal samples and four broad cell types. We identified 601,377 significant unique caQTLs (*p* < 1x10^-5^), corresponding to 412,604 unique caQTL variants (caVariants) and 102,797 unique caQTL peaks (caPeaks) (**Figure 3D**). caQTLs were also enriched near the peak start site (**Figure 3D**). Of note, we observed substantial variation in caQTL discovery across cell types and developmental stages. In particular, inhibitory neurons had relatively few caQTLs (**Figure 3D**). To validate caQ-TLs we compared them with previously published caQTL dataset from adult bulk brain tissue and T cells^35^ (**Figure S3F**). caQTLs in postnatal excitatory neurons, inhibitory neurons, and glia showed high correlation of effect sizes, supporting robust caQTL discovery. In our dataset, prenatal caQTLs and postnatal non-neuronal cell caQTLs had limited power and did not correlate well with existing caQTL from adult tissues.

Considering postnatal cell types, we assessed overlap between significant caQTLs and eQTLs (**Table S18**). Among significant caQTL (*p* < 1x10^-5^) variants, 9.7% were also identified as significant eQTL variants in excitatory neurons, 10.3% in Glia, and 13.2% in inhibitory neurons. Of these overlapping ca/eQTLs, in 51.8% the eQTL gene was the gene nearest the corresponding caQTL in excitatory neurons, 43.6% in Glia, and 53.2% in inhibitory neurons. These figures are consistent with previous studies showing that only a fraction of caQTLs yield changes in the transcriptome.^35^

About half of postnatal caQTLs peaks were consistent across major cell types and development, while the other half were associated with cell type/developmental DARs (**Table S17**, **Figure 3D**). Individual caQTL loci highlighted genotype-dependent regulatory element activity in specific cell types, for example, rs9831088 in an intron of *ATG3* (associated with neurogenerative diseases), was associated with chromatin accessibility in postnatal excitatory neurons (effect size = +0.75, *p* = 9.5x10^-10^) and overlapped an accessible enhancer-like element (**Figure 3E**). Similarly, rs17123658 in a *DNMT3B* intron (associated with intellectual disability), acted as a postnatal caQTL in glia (effect size = -0.88, *p* = 4.4x10^-10^); this is near a known enhancer element (ID: GH20J032791,^37^ chr20:32791539-32794727, hg38). These examples link common noncoding variation to cell-type- and development stage-predominant chromatin activity in the human pre-frontal cortex. Assessing motifs in caQTL-associated peaks across development (**Table S16**) identified prenatal enrichment of developmental and cell-fate transcription factors, such as homeobox, *FOX*, *SOX*, and T-box factors, whereas postnatal-predominant caQTLs captured signal-responsive and homeostatic regulators, including *AP-1/bZIP, IRF*, *ATF*, *KLF,* and *NFE2L2-*related motifs. These stage-specific differences were observed across neuronal and glial populations, although the specific regulatory programs varied by cell type.

### Dorsolateral prefrontal cortex splicing QTLs with cell type and developmental stage resolution

To identify sQTLs, we mapped changes in gene transcripts in prenatal and postnatal bulk tissue separately without cell type resolution. We identified 713,252 significant sQTLs, corresponding to 301,787 unique sQTL variants (sVariants) and 34,652 unique sQTL target genes (**Figure 3F**). sQTL variants were enriched near splice junction start sites, consistent with local genetic control of RNA processing (**Figure 3F**). Like eQTLs and caQTLS, sQTLs were dynamic across development with 46% of sQTLs being prenatal-predominant, 30% postnatal-predominant, and 24% shared. We validated our sQTLs by comparing effect sizes with previously published sQTL from genotype-tissue expression (GTEx)^34^ (**Figure S3G**). The strong correlation of effect sizes confirmed the reproducibility of our sQTLs and demonstrated that our analysis provides additional developmental resolution to splicing QTLs previously identified in the literature. Representative sQTLs showed stage-dependent effects on transcript structure. For example, rs80872 was associated with altered splicing of *CARS1* (associated with autosomal recessive developmental delay), with stronger effects in prenatal developmental stage (effect size = +1.3, *p* = 7.0x10^-16^), whereas rs58416198 was associated with postnatal-predominant splicing (effect size = +1.1, *p* = 3.9x10^-7^) of *SH3YL1* (associated with several neuropsychiatric conditions) (**Figure 3G**). Comparing sQTLs to eQTLs we saw that most sGenes were distinct from eGenes (**Figure 3H**), in keeping with complementary regulatory mechanisms.

Together, these results map the influence of common genetic variation on gene regulation involving transcription, chromatin accessibility, and splicing in specific cell types of human prefrontal cortex over development. Because many molecular QTLs are dynamic across developmental stages and cell types, they provide a resource to help interpret the biological implications of common variation linked to human traits and disorder by GWAS.

### Mechanistic insights to brain-related traits and disorders

GWAS have provided hundreds of genetic loci associated with brain-related traits and disorders, however, progressing from a GWAS locus to a specific gene, direction of effect, or, ideally, causal variant, remains a challenge. Identifying such genes, effects, and variants can provide neurobiological insights and present a therapeutic target.^38^ By integrating common variants from GWAS and molecular QTL analyses, we can assess colocalization to implicate a gene and direction of effect in the trait. Colocalization analyses across 24 GWAS, thousands of genes, and many thousands of variants are available through the BrainVar website (www.brainvar.org/). As a validation, we replicated the previously reported^39^ colocalization between with age of walking and *RBL2*; this relationship was especially strong for prenatal excitatory and inhibitory neurons (**Figure S4I–K**). Here, we highlight three disorder-associated loci with cell-type- and development-predominant colocalization of eQTLs, providing mechanistic insights (**Figure 4; Figure S4; Table S19**).

**Figure 4.**
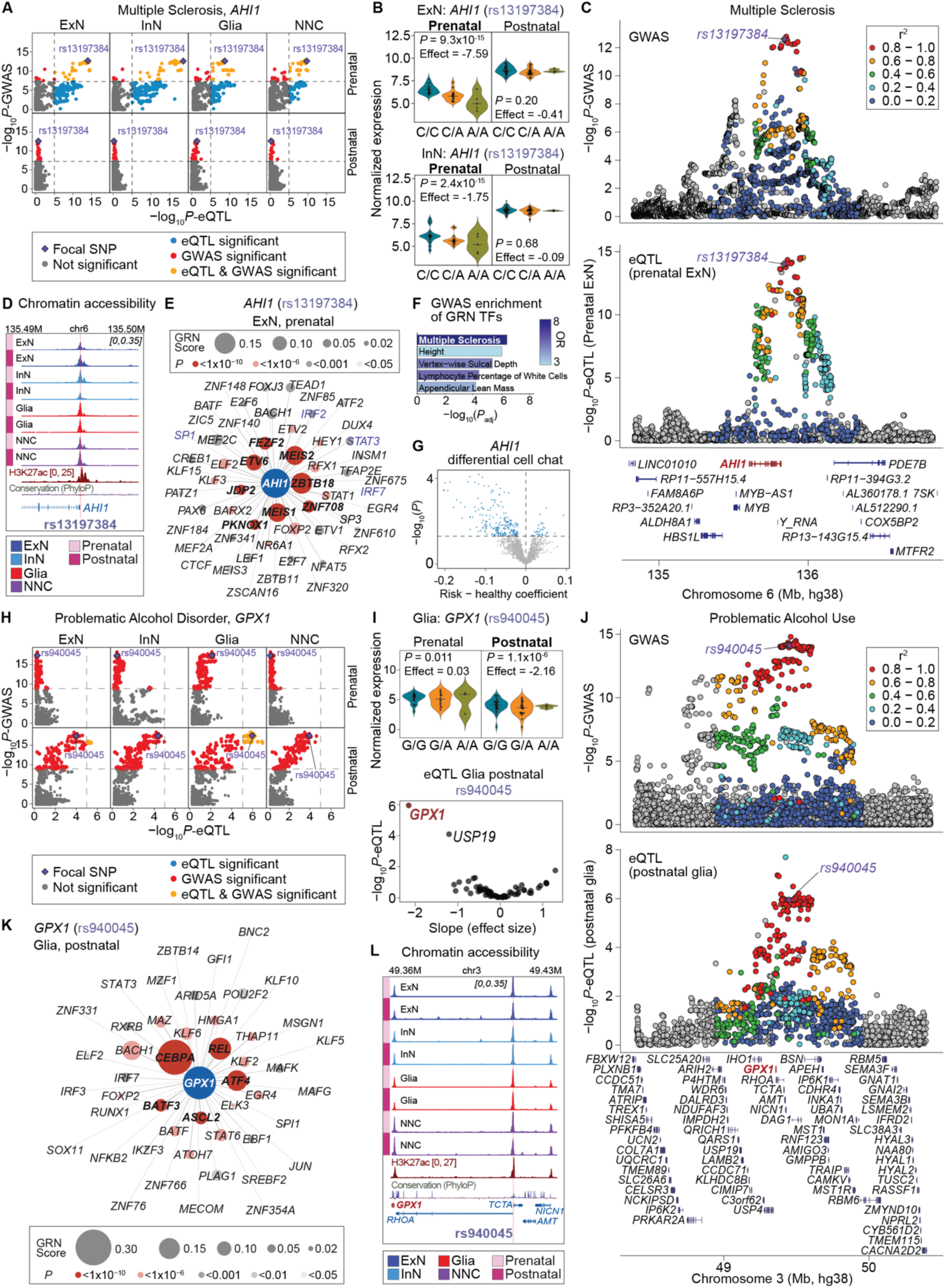
Cell-type- and developmental-stage-predominant colocalization of common variants underlying MS and problematic alcohol use disorder. **(A)** Scatterplots showing colocalization of MS GWAS variants at the AHI1 locus with eQTLs across excitatory (ExN), Inhibitory (InN), Glia, and non-neuronal cells (NNC) during prenatal and postnatal development stages. Scatterplots compare GWAS and eQTL association significance for variants within the locus (1e6 up and downstream of the GWAS lead variant). Variants are colored according to GWAS/eQTL significance overlap. The candidate causal variant is highlighted in purple. The genome-wide significance threshold for GWAS associations is *p* < 5e-8 and the significance threshold for eQTL associations is *p* < 1e-5. Explore more in BrainVar website (www.brainvar.org/). **(B)** Genotype-stratified *AHI1* expression for the candidate causal variant rs13197384 in prenatal and post-natal excitatory and inhibitory neuron populations. Violin plots show raw normalized expression (log_2_CPM+1; not adjusted for covariates) across genotypes. Genotypes are shown as homozygous reference (blue), heterozygous (yellow), and homozygous alternate (green). *P* values and effect sizes were calculated using TensorQTL. **(C)** LocusZoom plot showing MS GWAS and prenatal excitatory eQTL association signals at the *AHI1* locus. Variants are colored according to linkage disequilibrium (r²) with the lead/causal SNP rs13197384. **(D)** Chromatin accessibility UCSC tracks across developmental stages and cortical cell types at the *AHI1* locus. Tracks show ReadsInTSS normalized pseudobulk ATAC-seq accessibility. The candidate causal variant rs13197384 is highlighted. H3K27ac is ChIP-seq on dlPFC tissue female adult (89 years).^36^ Conservation score shows phyloP conservation visualized using UCSC genome browser. **(E)** Gene regulatory network of *AHI1,* rs13197384 variant in prenatal excitatory. Node color indicates statistical significance (calculated from ordinary least squares regression) and node size reflects median network score (**Table S20**, see also **Method** section for more information). **(F)** Enrichment analysis of genes within the *AHI1*, rs13197384 prenatal excitatory neuron gene regulatory network against the GWAS Catalog 2023 using Enrichr.^42^ Enrichment results are sorted based on -log10 adjusted *p* and colored based on odd ratio. **(G)** Volcano plot of differential CellChat analysis showing ligand–receptor interactions differentially affected in rs13197384 MS genetic risk donors compared with healthy donors. The x-axis shows the coefficient, indicating a decrease or increase in signaling, while the y-axis shows the −log10 *p* calculated by the Cell-Chat differential analysis (**Table S23**). **(H)** Scatterplots showing colocalization of problematic alcohol disorder GWAS variants at the *GPX1* locus with eQTLs across different cell types and developmental stages. Scatterplots compare GWAS and eQTL association significance for variants within the locus (1e6 up and downstream of the GWAS lead variant). The candidate causal variant is highlighted in purple. The genome-wide significance threshold for GWAS associations is *p* < 5e-8 and the significance threshold for eQTL associations is *p* < 1e-5. **(I)** Genotype-stratified *GPX1* transcription for the candidate causal variant rs940045 in prenatal and postnatal glial populations. Violin plots show raw normalized expression (log_2_CPM+1; not adjusted for covariates) across genotypes. *p* values and effect sizes were calculated using TensorQTL. Bottom: eQTL effect sizes and *p* values for all genes associated with rs940045 within the postnatal glial *GPX1* locus, demonstrating that *GPX1* is the primary significantly affected gene with the strongest effect size. Each point represents an individual gene. See also **Figure S4C**. **(J)** LocusZoom plot showing problematic alcohol disorder GWAS and postnatal glial eQTL association signals at the *GPX1* locus. Variants are colored according to linkage disequilibrium (r²) with rs940045. **(K)** Gene regulatory network of *GPX1,* rs940045 variant in postnatal glial populations. Node color indicates statistical significance and node size reflects median network score (**Table S20**, see also **Method** section for more information). **(L)** Chromatin accessibility profiles across developmental stages and cortical cell types at the *GPX1* locus. Track shows pseudobulk ATAC-seq accessibility (ReadsInTSS normalized). The candidate causal variant rs940045 is highlighted. See all candidates in **Table S19**. All genomic coordinates correspond to the hg38 human reference genome assembly. See also **Figure S4** and BrainVar website (www.brainvar.org/). **Abbreviations**: ChIP-seq, chromatin immunoprecipitation sequencing; CPM, counts per million; dlPFC, dorsolateral prefrontal cortex; eQTLs, expression quantitative trait loci; MS, multiple sclerosis.

### *AHI1*-mediated risk for multiple sclerosis in prenatal neurons

An MS GWAS with over 700,000 individuals identified 236 genome-wide significant loci.^40^ Analy-sis of gene expression from the resulting gene list implicated both microglia and neurons,^40^ a result we replicate (**Figure S3B/C**). At the MS-associated rs4896153 locus (chr6:135,833,463, hg38), we identified a robust prenatal-predominant colocalization signal (PPH4 = 0.98) to the gene *AHI1* (Abelson helper integration site 1), especially in neurons (**Figure 4A**). *AHI1* encodes Jouberin, a scaffold protein associated with primary cilia implicated in neuronal differentiation, axonal guidance, and cerebellar and cortical development. Complete knockout of *AHI1* leads to Joubert Syndrome,^41^ a recessive neurodevelopmental disorders characterized by characterized by hypoplasia of the cerebellar vermis. *AHI1* is highly expressed, particularly in neuronal populations, with expression increasing across cortical development (**Figure S4A**). No other gene in the locus had a high colocalization posterior probability in prenatal or postnatal cells (**Figure S4B**). Genotype-stratified analyses demonstrated that the MS risk allele (odds ratio = 1.14, *p* GWAS = 2.4x10^-13^) was associated with reduced prenatal *AHI1* expression (effect size = -7.59, *p* eQTL = 9.3x10^-15^; **Figure 4B, 4C**). Integrating our chromatin accessibility data with cross-species conservation, we highlighted rs13197384 as a likely casual variant (**Table S19**) residing in an active prenatal neuronal regulatory element in the *AHI1* promoter (**Figure 4D**).

To investigate the broader regulatory environment surrounding *AHI1*, we assessed gene regulatory networks within prenatal excitatory neurons (**Figure 4E; Table S20**). *AHI1* was regulated by a densely connected developmental module enriched with transcription factors implicated in neuronal differentiation and cortical patterning, including *MEIS1*, *MEIS2*, *ETV6*, *KLF3*, and *FOXP2*. Comparing the genes in this regulatory network to the GWAS Catalog, the most significant enrichment was for MS (adjusted *p* = 2.7x10^-4^, Odd ratio = 8.30, Enrichr^42^ (**Figure 4F**; **Table S21**). Assessing our putative causal allele (rs13197384), we performed a motif break analysis. The alternative allele (C>A) was predicted to impact four genes in the regulatory network, strengthening binding for STAT3 and IRF7 (ΔPWM scores = 1.58 and 1.30, respectively) and weakening SP1 binding (two overlapping motifs: ΔPWM scores = −0.73 and −0.72) (**Figure 4E; Table S22**). The gene *STAT3* has previously been implicated as a genome-wide significant peak in multiple sclerosis potentially acting via neurons.^40^ Given the role of primary cilia in cell-cell signaling, we performed a differential CellChat analysis^43^ in prenatal cells from donors homozygous for the rs13197384 risk allele (A/A; 7 donors) or non-risk allele (C/C; 22 donors). As a result, 23 ligand/receptor interactions met an FDR threshold < 0.1 and the risk allele was associated with substantial decrease in predicted interactions (**Figure 4G**; **Table S23**), especially related to microglia and neurons and microglia and oligodendrocyte precursor cells acting via glutamate signaling (e.g., SLC1A2) and trans-synaptic adhesion (e.g., NRXN1).

Collectively, these results suggest a mechanism of MS risk in which increased STAT3 binding leads to decreased *AHI1* expression in prenatal neurons, resulting in impaired cell-cell signaling between neurons, microglia and oligodendrocyte precursor cells, presumably increasing susceptibility to neuroimmune dysfunction later in life.

### *GPX1*-mediated risk for problematic alcohol use in postnatal glia

At the rs66518420 locus (chr3:49,564,316; hg38) associated with problematic alcohol use (PAU) from over 1 million individuals,^44^ we identified a colocalization signal (PPH4 = 0.96) to the gene *GPX1* centered on rs940045 and strongest in postnatal glia (**Figure 4H-J; Table S19**). The risk allele was associated with reduced *GPX1* (beta = -0.015, *p* GWAS*=* 7.7x10^-15^) expression in postnatal glial cells (effect size = -2.16, *p* eQTL = 1.1x10^-6^) (**Figure 4I**). Within the locus, the gene *USP19* also showed a trend towards an eQTL (**Figure 4I; S4C**). The regulatory relationship between rs940045 and *GPX1*, but not *USP19*, was further supported by eQTLs from GTEx frontal cortex,^16^ as well as a single cell amygdala dataset that identified a microglia-specific cRE within which this SNP resides.^45^ *GPX1* encodes glutathione peroxidase 1, a key antioxidant enzyme that protects cells from oxidative stress. Because chronic alcohol exposure induces oxidative damage and inflammatory responses in the brain, reduced *GPX1* expression may impair the ability of glial cells to maintain redox homeostasis.^46–48^ Consistent with this mechanism, *GPX1* is highly expressed in glial populations during postnatal cortical maturation (**Figure S4A**), and rs940045 resides in an ATAC-seq peak between the TSS of the genes *RHOA* and *TCTA*, about 50kbp up-stream of the *GPX1* TSS (**Figure 4J**). Gene regulatory network analysis further placed *GPX1* within a glial transcriptional module enriched for inflammatory and stress-response regulators, including *BATF*, *IRF7*, *STAT6*, and *NFKB2* (**Figure 4K; Table S20**). Together, these findings suggest that genetic susceptibility at the *GPX1* locus may contribute to PAU disorder through disruption of postnatal glial oxidative stress and neuroimmune pathways.

### *FADS1*-mediated risk for mood disorders in postnatal neurons

A tandem gene cluster of three fatty acid desaturase genes (*FADS1*, *FADS2*, *FADS3*) has been associated with both MDD and bipolar disorder.^14,49^ We identified colocalization signals (MDD PPH4 = 0.97, BIP PPH4 = 0.96) with *FADS1*, but not *FADS2* or *FADS3* in postnatal neurons (**Figure S4E–G; Table S19**). Genotype-stratified analyses demonstrated that the risk-associated allele (odds ratio = 0.94, *p* GWAS = 3.2x10^-16^) was linked to reduced *FADS1* expression (effect size = -2.35, *p* eQTL = 1.3x10^-6^) in postnatal neuronal populations. *FADS1* encodes a key enzyme involved in polyunsaturated fatty acid and lipid metabolism, pathways essential for neuronal membrane composition, synaptic function, and neuroinflammatory regulation. Altered lipid metabolism has been repeatedly implicated in psychiatric disorders, including depression and bipolar disorder.^14,49,50^ We implicate rs174562 as a putative casual variant. This SNP is located in the *FADS1* promoter overlapping an ATAC-seq peak across all cell types and developmental stages (**Figure S4H**), suggesting prolonged regulatory activity throughout cortical maturation. Together, these findings suggest that mood disorder-associated variation at the *FADS1* locus may influence psychiatric susceptibility through persistent disruption of neuronal lipid metabolic and neuroinflammatory pathways across development.

### Defining therapeutic targets for genetic medicines

Advances in genetic medicines raise the potential to treat rare genetic disorders by modifying the expression or translation of genes in the human brain. To assess the potential of our atlas to nominate therapeutic targets, we assessed whether the developmental and cell type DARs we identified (**Figure 1C, E**) might predict critical cis-regulatory elements (cREs) that can influence gene expression.^51,52^ Distal cREs (e.g., enhancers) are more likely to have cell type specificity that proximal cREs (e.g., promoters) making a potential therapy more specific. However, a recent high-throughput CRISPRa screen of 337 NDD-associated genes in iPSC-derived neurons found that predicting suitable distal cRE targets was substantially harder than for proximal cREs.^53^ Considering the 100 distal cREs they identified as CRISPRa targets,^53^ all of them had neuronal ATAC-seq peaks in our data and 74 of them overlapped our neuronal DARs (**Figure S5A**). For example, an excitatory neuron-specific DAR in a *SATB2* intron was a CRISPRa-responsive cRE for *SATB2*^53^ (**Figure S5B**).

Given this relationship between DARs and CRISPRa-responsive cREs, we considered whether we could leverage our DARs to predict novel CRISPRa-responsive cREs in genes associated with NDD. To achieve this, we created a comprehensive list of putative cREs in 164 NDD-associated genes (∼75 per gene). Each cRE was annotated for developmental/cell-type DAR overlap, known^37,54–56^ and predicted regulatory-elements,^57^ H3K27ac,^36^ Activity-by-Contact (ABC) scores,^53,58^ sequence conservation across species,^59^ and chromatin accessibility in SH-SY5Y cells obtained from GEO [GSM2700775]^60^ (**Figure S5C; Table S24**). We also calculated co-accessibility using cicero (**Table S24; Figure S5C**), identifying developmental changes in co-accessible peaks (**Figure S5D**) and both direct distal-proximal co-accessibility and membership in cis-co-accessibility networks (CCANs) for candidate cREs (**Table S24**). Within these cREs, we focused on co-accessible neuronal DARs with open chromatin in SH-SY5Y cells and selected a distal cRE for *SOX5* for validation (**Figure 5C–F; Table S24**). This cRE was differentially accessible in prenatal excitatory neurons (log_2_FC=1.08, FDR=9.2x10^-20^) but retained substantial excitatory neuron accessibility postnatally (**Figure 5C-E**) and was accessible in SH-SY5Y cells for experimental validation. The ABC score (0.044) was consistent with the cRE acting as a distal cRE to the *SOX5* promoter (**Figure 5F; Table S24**).

**Figure 5.**
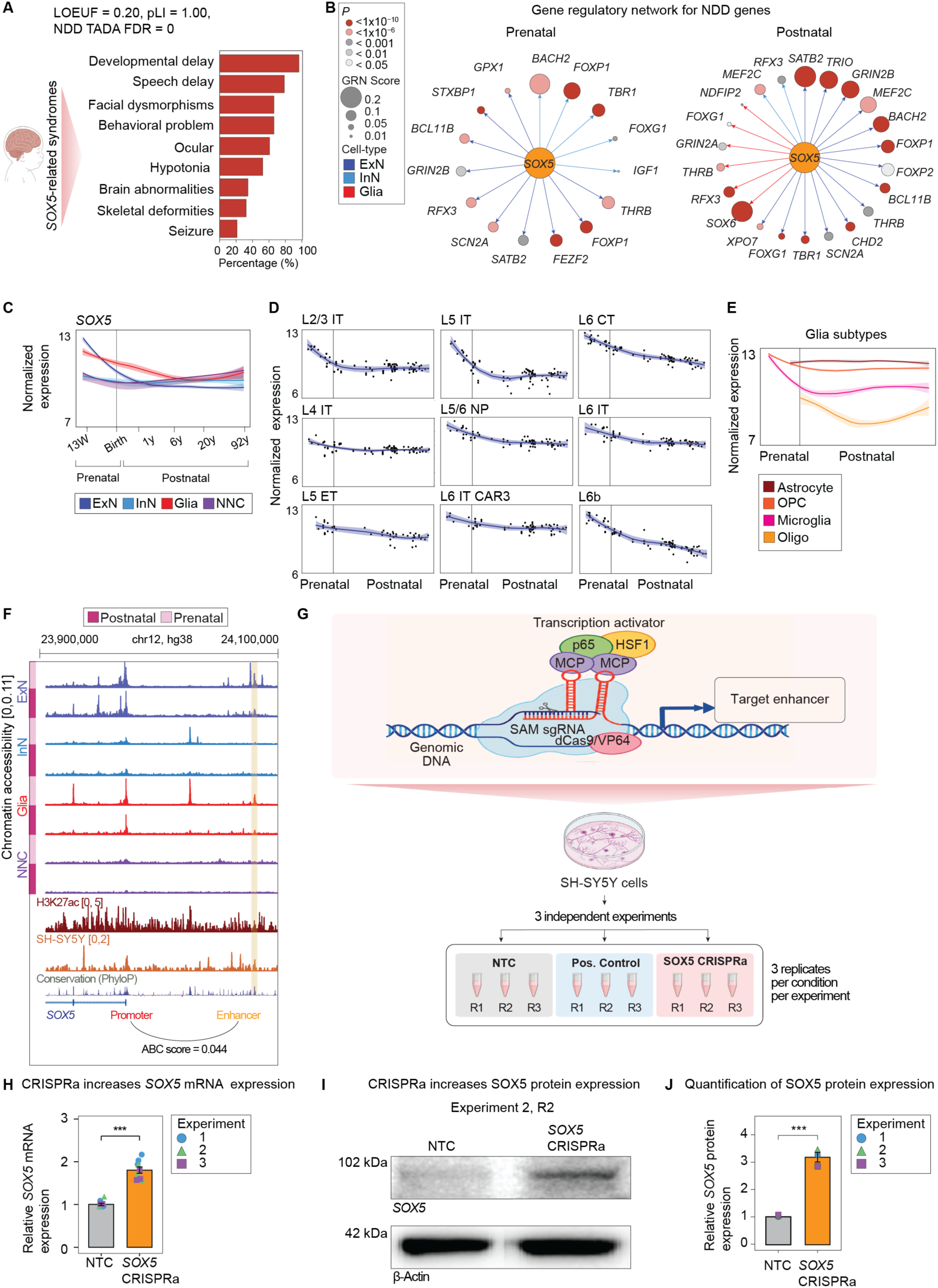
CRISPRa-mediated targeting of an enhancer element to increase SOX5 protein expression in SOX5 haploinsufficiency syndromes. **(A)** Clinical features associated with *SOX5*-related syndrome/Lamb–Shaffer syndrome. Bars indicate the percentage of affected individuals exhibiting each reported phenotype based on DECIPHER (https://www.deciphergenomics.org). Gene-level metrics shown above indicate strong intolerance of *SOX5* to loss-of-function variation and genetic association with neurodevelopmental disorders (NDDs): LOEUF = 0.20, where lower values indicate greater depletion of predicted loss-of-function variants relative to expectation; pLI (probability of loss-of-function intolerance) = 1.00, indicating strong intolerance to heterozygous loss-of-function variation; and NDD TADA FDR = 0, indicating strong statistical evidence for association of *SOX5*-disrupting variants with NDDs. **(B)** Gene regulatory networks of *SOX5* during prenatal and postnatal brain development in, glia, excitatory and inhibitory neurons. Nodes represent NDD-associated genes regulated by *SOX5* in the inferred regulatory network. **(C)** Smoothed *SOX5* expression across major brain cell types from 13 weeks post-conception through 92 years of age. Lines indicate normalized (log_2_CPM+1) *SOX5* expression trajectories in excitatory and inhibitory neurons, glia, and non-neuronal cells. **(D)** Smoothed expression of *SOX5* across excitatory neuronal subtypes. Normalized (log_2_CPM+1) SOX5 expression is shown across prenatal and postnatal development for L2/3 IT, L4 IT, L5 IT, L5/6 NP, L6 CT, L5 ET, L6 IT CAR3, and L6b populations. Points represent individual observations, and curves indicate fitted developmental expression trajectories. **(E)** Smoothed *SOX5* expression (log_2_CPM+1) across glial subtypes, including astrocytes, oligodendrocyte precursor cells, microglia, and oligodendrocytes. **(F)** Chromatin accessibility and regulatory annotation of the *SOX5* enhancer. Normalized chromatin accessibility (ReadsInTSS normalized) profiles are shown for prenatal and postnatal excitatory and inhibitory neurons, glia, and non-neuronal cell populations, together with H3K27ac and chromatin accessibility profiles from SH-SY5Y cells. The *SOX5* promoter and candidate enhancer at chr12:24081200–24081530 (hg38) are highlighted. The predicted enhancer–promoter connection has an ABC score of 0.044. **(G)** Experimental design for CRISPR activation of the candidate SOX5 enhancer. A dCas9-based transcriptional activation complex is recruited to the candidate enhancer by sgRNAs to increase endogenous *SOX5* transcription. SH-SY5Y cells were subjected to three independent experiments comprising non-targeting control (NTC), SCN2A CRISPRa as positive control (Pos. Ctrl),^52^ and SOX5 enhancer-targeting CRISPRa conditions, followed by assessment of *SOX5* mRNA and protein abundance. **(H)** CRISPRa targeting of the *SOX5* enhancer increases *SOX5* mRNA expression (qPCR) relative to the NTC condition in SH-SY5Y cells. Points denote measurements from three independent experiments, and bars summarize relative *SOX5* expression. See also **Figure S5E**. **(I)** CRISPRa targeting of the *SOX5* enhancer increases *SOX5* protein abundance. Representative (Experiment 2, R2) immunoblots of *SOX5* in NTC and *SOX5* enhancer-targeting CRISPRa conditions. β-Actin staining is shown as a protein-loading control. See also **Figure S5F**, **J**. **(J)** Quantification of *SOX5* protein expression. Densitometric analysis of *SOX5* protein expression in NTC and *SOX5*-transduced cells. *SOX5* band intensities were normalized to β-actin and expressed relative to the NTC condition. Three independent experimental runs were performed, with three replicate wells analysed per condition in each experiment. Replicate wells were averaged within each experimental run, and each point represents the mean of one experimental run (n = 3). Bars represent mean ± SEM. Statistical significance was assessed using a linear model with experimental run included as a blocking factor. **Abbreviations:** ABC, activity-by-contact; CPM, counts per million; CRISPRa, CRISPR activation; CT, corticothalamic; ET, extratelencephalic; FDR, false discovery rate; IT, intratelencephalic; LOEUF, loss-of-function observed/expected upper bound fraction; NDDs, neurodevelopmental disorders; NP, near-projecting; NTC, non-targeting control; pLI, probability of loss-of-function intolerance; SEM, standard error of the mean; TADA, transmission and de novo association.

Haploinsufficiency of *SOX5* causes a neurodevelopmental disorder characterized by developmental delay, speech and language impairment, and facial dysmorphism (SOX5-syndrome, eponym Lamb Shaffer) (**Figure 5A**).^61,62^ The *SOX5* gene encodes an SRY-related HMG-box transcription factor with critical roles in cortical development, neuronal differentiation, and neuronal specification.^62,63^ Consistent with these roles, our GRN analysis shows *SOX5* participating in developmental transcriptional programs involving multiple important NDD-associated genes, including *SCN2A*, across prenatal and postnatal stages (**Figure 5B**). Thus, upregulation of *SOX5* would potentially improve symptoms of *SOX5*-syndrome.

CRISPRa targeting of this distal cRE reproducibly increased *SOX5* mRNA expression in SH-SY5Y cells across three independent experiments (**Figure 5G**). Relative to the non-targeting control, *SOX5* expression increased by 2.04-, 1.70-, and 1.66-fold in experiments 1, 2, and 3, respectively (n = 3 biological replicates per experiment; n = 9 biological replicates in total), corresponding to a mean 1.80-fold increase across experiments (linear model with experimental run included as a blocking factor, *p* = 8.4x10^-4^, (**Figure 5H; Figure S5E; Table S25**). Importantly, the increase in *SOX5* transcript abundance was accompanied by increased *SOX5* protein levels. Immunoblotting followed by normalization of *SOX5* signal to total protein (β-actin) revealed 3.21, 3.37, and 2.8-fold increases in experiments 1, 2, and 3, respectively, relative to the non-targeting control (n = 3 biological replicates per experiment; n = 9 biological replicates in total), corresponding to a mean 3.13-fold increase across experiments (linear model with experimental run included as a blocking factor, *p* = 1.0x10^-3^; **Figure 5I,J; Figure S5F, J**; **Table S26**). Together, these results provide proof-of-concept that the cell typeand developmental stage-resolved regulatory architecture captured by our atlas can prioritize functional cis-regulatory elements capable of increasing the mRNA and protein expression of haploinsufficient NDD genes.

## DISCUSSION

Disruption of the structure or function of the brain, the most complex organ in the known universe, is “the leading cause of overall disease burden in the world”.^64,65^ We are far from understanding the development, physiology, and pathology of the brain. Here, we present a step on the journey by producing a comprehensive genetic and cellular atlas of the developing human dlPFC.

We observe substantial developmental changes across gene regulation, gene expression, and gene splicing with many such changes varying across cell types. In general, genes associated with human traits are more likely to show dynamic regulatory patterns compared to those without, in keeping with evolutionary need to ‘control’ critical genes. We can use these dynamic patterns to provide insights into when, where, and how gene disruption leads to human traits. Such approaches have been performed before with other datasets, especially transcriptomic datasets.^7,19,22,23,66^ With this more comprehensive dataset, we largely draw the same conclusions: neurodevelopmental disorders are enriched in neuronal programs, with strong signals in prenatal neurons but also postnatal signal; psychiatric disorders are also enriched in neurons, with prenatal vs. postnatal enrichment varying across disorders; and neurodegenerative disorders are enriched in postnatal microglia programs (**Figure 2**). Such analyses rest on the assumption that high gene expression is causal. At the extremes (expressed vs. not expressed) this assumption is logical; however, enrichment analyses rely on subtler quantitative changes. Our dataset enables us to assess enrichment patterns across three orthogonal approaches: gene enrichment in expression patterns (**Figure 2A,D**), gene enrichment in chromatin peaks (**Figure 2A,D**), and polygenic score correlation in chromatin peaks (**Figure 2C**). The consistent observed patterns are similar across these approaches and robust to how dynamic genes are determined (**Figure 2A,D**), lending some support to the conclusions. Of note, increasing the cell type resolution did little to further resolve neurodevelopmental or neuropsychiatric disorders, possibly indicating a pan-neuronal effect (e.g. synaptic plasticity) or that larger cross-regional atlases and more accurate gene lists will be required. Cellular resolution did provide some additional insight in neurodegenerative disorders, with enrichment in prenatal progenitor cells, neurons and microglia in multiple sclerosis (MS) (**Figure S2B,C**).

Within the cellular and developmental compartments, we saw evidence of shared gene function and developmental trajectories, supporting this notion of a coherent biological context relevant to specific disorders. For example, prenatal inhibitory neurons were highly enriched for NDD-associated genes and these genes converged on the SWI/SNF complex and related chromatin regulators (**Figure 2E, S2D,E**). We also noted that gene function predicted different expression trajectories within a cell type. For example, ASD-associated genes related to transcriptional regulation were expressed at a higher level in prenatal neurons, while genes related to neuronal communication were expressed at a higher level in postnatal neurons (**Figure 2B, S2F**); it remains to be seen whether this predicts therapeutic response at a given age. This trajectory analysis also identified a schizophrenia-specific enrichment (**Figure S2F**) in non-neuronal cells, especially vascular leptomeningeal cells. To further explore these relationships, we performed a cross-trait factorization analysis within enriched cell types and developmental stages. Factorization identified five domains across brain-related traits, highlighting shared genetic vulnerabilities (**Figure 2F**).

We also noted dynamic effects in molecular QTLs across development and/or cell type (**Figure 3**). While sample size is the main determinant for QTL discovery, resolution across cell-type, developmental stage, brain region, and cell state (e.g., neuronal activity) are likely to be necessary to interpret QTLs. To this end, we highlighted three dynamic QTLs that colocalize with brain-related disorders. Of note, we find that the strongest colocalization results at the rs4896153 locus in MS^25^ were obtained for the gene *AHI1* in prenatal excitatory and inhibitory neurons (**Figure 4**). Integrating the modalities of our dataset, we were able to identify a potential mechanism, with rs13197384, a putative causal variant in the *AHI1* promoter, enhancing the binding of STAT3, a MS-associated transcription factor,^67^ leading to reduced *AHI1* expression and decreased ligand-receptor signaling between prenatal neurons and microglia. Of note, the predicted gene regulatory network targeting *AHI1* was also enriched for MS-associated genes. We note that substantial validation would be required to confirm this mechanism but that these data provide the basis with which to form and test such hypotheses. We also provide increased resolution for relationships between eQTLs targeting *GPX1* in problematic alcohol use in postnatal glia (**Figure 4**), lending support to the role of oxidative stress in this trait, and *FADS1* in depression and bipolar disorder in postnatal excitatory neurons (**Figure S4**), highlighting the role of neuronal regulation and lipid metabolism in these disorders, both of which are important for synapse function^68^ and neuroinflammatory balance.^69,70^

Finally, we propose that our identified dynamic regulatory regions can be applied to guide therapeutic insights. Notably, enhancer elements that could be targeted to upregulate genes in a CRISPR activation genomic screen of NDD-associated genes in iPSC-derived neurons were sub-stantially more likely to vary across cell type and/or development (**Figure S5**). Based on this result, we assessed whether the dynamic regulatory regions we had identified could guide the identification of further CRISPR activation NDD gene targets. Integrating additional genomic data including co-accessibility in our dataset, we selected a putative enhancer of *SOX5*, the causal gene of haploinsufficient *SOX5* syndrome. We demonstrated that CRISPR activation of this enhancer element in SH-SY5Y cells increased *SOX5* RNA expression and SOX5 protein levels. This is an important proof of principle that suggests that developmental regulatory maps can help identify therapeutic target in addition to disorder mechanisms.

### Limitations of the study

Our study has several limitations. The interpretation of enrichment analyses requires caution. This methodology assumes that enrichment of disorder-associated genes within stronger open chromatin peaks or higher expressed genes of a particular cell type or developmental stage reflect the biological context most relevant to genetic risks. While this is a useful approach for prioritizing candidate cellular and developmental mechanisms, it does not establish causality. Furthermore, disorder-associated genes may function across multiple cell types and developmental periods, and the context showing the strongest enrichment may not necessarily represent the primary site of disorder pathogenesis. Therefore, our enrichment results should be viewed as hypotheses that identify candidate disorder-relevant programs for future experimental investigation. Next, although our atlas spans a broad developmental range, some developmental periods and cell populations remain underpowered. To maximize statistical robustness, our QTL analyses focused on broad cell types and prenatal versus postnatal stages rather than finer developmental intervals. Larger cohorts with increased sample size across developmental periods will enable higher-resolution analyses of genetic regulation and may reveal additional patterns. Also, colocalization does not establish causality. Although colocalization identifies loci where molecular QTL and GWAS signals are consistent with a shared underlying variant, it cannot definitively identify the causal variant, target gene, or molecular mechanism. We integrated multiple lines of evidence, including developmental specificity, chromatin accessibility, regulatory annotation, and prior literature, to prioritize candidate causal genes and variants. However, direct experimental validation will be required to confirm these regulatory mechanisms. Furthermore, we only considered dlPFC; the relative strength of evidence between different brain regions (or brain states) might lead to different conclusions (e.g., enrichment, colocalization). Finally, although we demonstrate that enhancer activation can increase expression of selected neurodevelopmental disorder genes, these experiments represent proof-of-concept studies. Substantial further work on both the payload and a vector would be required to make, dose, and deliver a safe and efficacious therapy. Despite these limitations, our study provides a developmental and cell-type-resolved resource for understanding how genetic variation influences molecular regulation in the human brain and contributes to brain disorder.

### Conclusion

Overall, our study shows that genetic liability in the human brain is shaped by both cell type and developmental timing. The same variant can act differently depending on when and where it is active. This makes developmental context essential for understanding brain disorders. It also provides a framework for moving from genetic association to mechanism and, eventually, to therapy.

## Extended Figures

**Figure S1.**
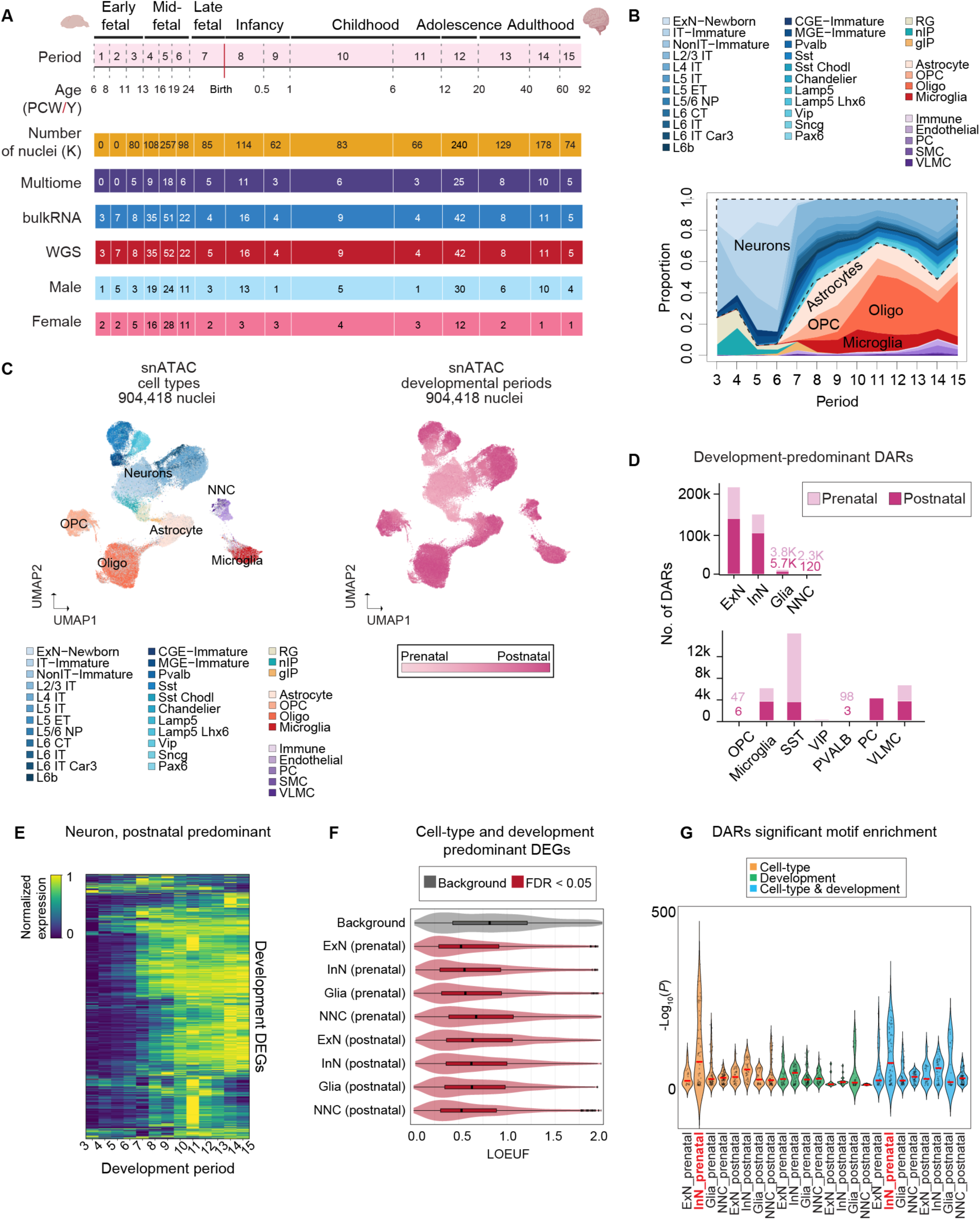
Cohort composition and developmental dynamics of cellular and regulatory programs, related to Figure 1. **(A)** Donor distribution across developmental periods (6 PCW–92 years) and modality (snMultiome, bulk RNA sequencing, and WGS) after preprocessing. Prenatal ages are reported in PCW, and postnatal ages in years. The heatmap displays nuclei counts (×10³) and donor numbers per developmental period, modality, and sex. In the snMultiome dataset, Period 1 donors were not included due to the lack of available tissue. Period 2 donors were also excluded for snMultiome analyses because they were determined to contain predominantly non-cortical cells, indicating inaccurate regional dissection (**Table S1** and **S2**). **(B)** Developmental shifts in major cortical cell subtypes, illustrating postnatal expansion of glial populations. The y-axis represents the proportion of cells, and the x-axis indicates developmental periods. **(C)** Top left: ATAC UMAP colored by annotated broad cell types. Top right: ATAC UMAP colored by developmental periods (Periods 3–15). **(D)** Numbers of prenatal- and postnatal-predominant DARs across broad cell types (adjusted *p* < 0.05, average log₂FC > 0.5) calculated by Wilcoxon rank sum test (**Data S2**). **(E)** Transcriptional changes of neuronal postnatal DEGs during development. The normalized expression shows min-max scaled pseudobulk average expression across periods for neurons (including both excitatory and inhibitory neurons). **(F)** Loss-of-function observed/expected upper bound fraction (LOEUF) distributions across cell type- and developmental stage-predominant DEGs. LOEUF scores from the gnomAD v2.1.1 gene constraint dataset^71^ were used to assess loss-of-function constraint across DEG sets. Distributions were compared with a background comprising all genes tested in the DEG analyses and represented in the gnomAD constraint dataset using two-sided Wilcoxon rank-sum tests. *P* values were corrected for multiple testing using the Benjamini–Hochberg procedure. DEG sets with LOEUF distributions significantly different from the background (FDR < 0.05) are highlighted in red (**Table S6**). **(G)** HOMER^72^ motif discovery results for cell type (orange), developmental stage (green), and joint cell type and development (blue) DARs. The y-axis shows the −log_10_ *p* value calculated by HOMER, and each dot represents a significantly enriched motif. Prenatal inhibitory neurons are highlighted in red because they exhibited the highest relative −log10 *p* values among the identified motifs (**Table S4**). **Abbreviations:** DARs, differentially accessible regions; DEGs, differentially expressed genes; FDR, false discovery rate; gnomAD, Genome Aggregation Database; HOMER, Hypergeometric Optimization of Motif EnRichment; LOEUF, loss-of-function observed/expected upper bound fraction; PCW, post-conception weeks; snMultiome, single-nucleus multiome; UMAP, uniform manifold approximation and projection.

**Figure S2.**
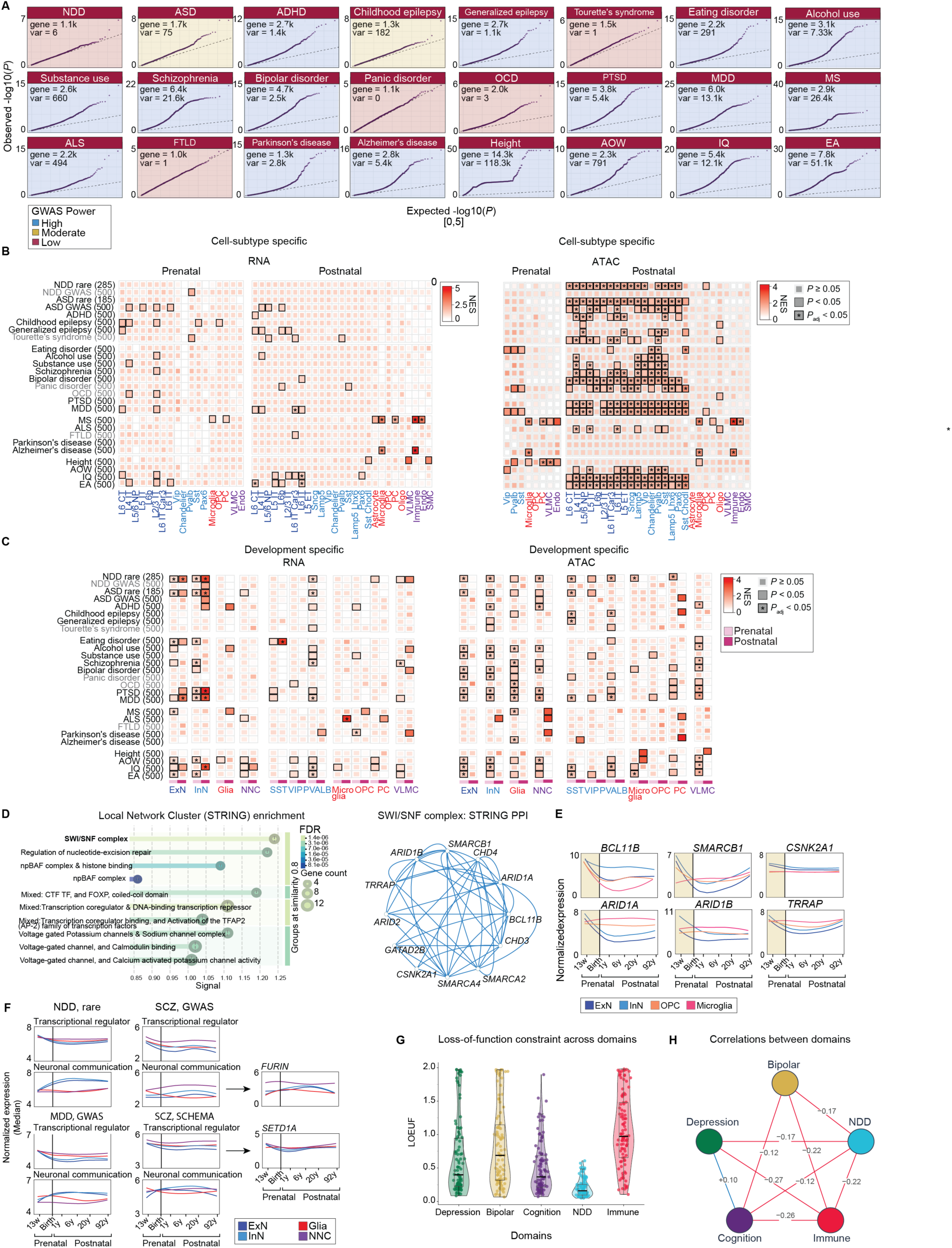
Extended analyses of cell-type and developmental enrichment across brain disorders related to Figure 2. **(A)** QQ plots of MAGMA gene association results for brain disorder GWAS datasets (see also **Table S7** and **DATA S5**). The number of genome-wide significant variants (*p* < 5 × 10⁻⁸) and MAGMA-significant genes (*p* < 0.05) are indicated for each trait (labelled as var and gene). Colors represent estimated GWAS statistical power, with blue indicating high power, yellow moderate power, and red low power. **(B)** Cell subtype–level enrichment analysis of brain disorder–associated genes using RNA expression and ATAC-derived gene markers (see also **Table S8**). Enrichment was performed separately for prenatal and postnatal stages for cell subtypes. Cell subtypes are color-coded as follows: excitatory neurons, dark blue; inhibitory neurons, light blue; Glia, red; non-neuronal cells, purple. Traits shown in grey represent those with low-powered GWAS data **(C)** Developmental stage–specific enrichment of brain disorder–associated genes using stage-resolved marker genes derived from RNA and ATAC datasets. Traits shown in grey represent those with low-powered GWAS data **(D)** Left: STRING protein–protein interaction analysis of genes enriched in prenatal inhibitory neurons for NDD (see also **Table S12**). Right: Interaction network highlighting convergence of NDD-associated genes enriched in prenatal inhibitory neurons on components of the SWI/SNF chromatin remodeling complex. **(E)** Expression patterns of SWI/SNF-enriched NDD genes showing high prenatal expression in inhibitory neurons. Expression values represent the median log₂-normalized CPM + 1. **(F)** Expression patterns of NDD-, MDD-, and SCZ-associated genes grouped by functional categories, including transcriptional regulation and neuronal communication. Two representative schizophrenia-associated genes are shown on the right. Expression values represent the median log₂-normalized CPM + 1 across genes within each developmental period and cell type (see also **Table S10**). **(G)** LOEUF score distributions among the top 100 genes in each biological domain (**Table S13**). Lower LOEUF scores indicate stronger intolerance to loss-of-function variants. An overall gene-level permutation test, accounting for genes shared across domains, revealed significant differences in LOEUF distributions (*p* = 0.000020; 50,000 permutations). **(H)** Pearson correlations between gene-loading scores across biological domains (**Table S13**). **Abbreviations**: CPM, counts per million; LOEUF, loss-of-function observed/expected upper bound fraction; MAGMA, multi-marker analysis of genomic annotation; MDD, major depressive disorder; NDD, neurodevelopmental disorder; SCZ, schizophrenia; STRING, Search Tool for the Retrieval of Interact-ing Genes/Proteins; SWI/SNF, switch/sucrose non-fermentable.

**Figure S3.**
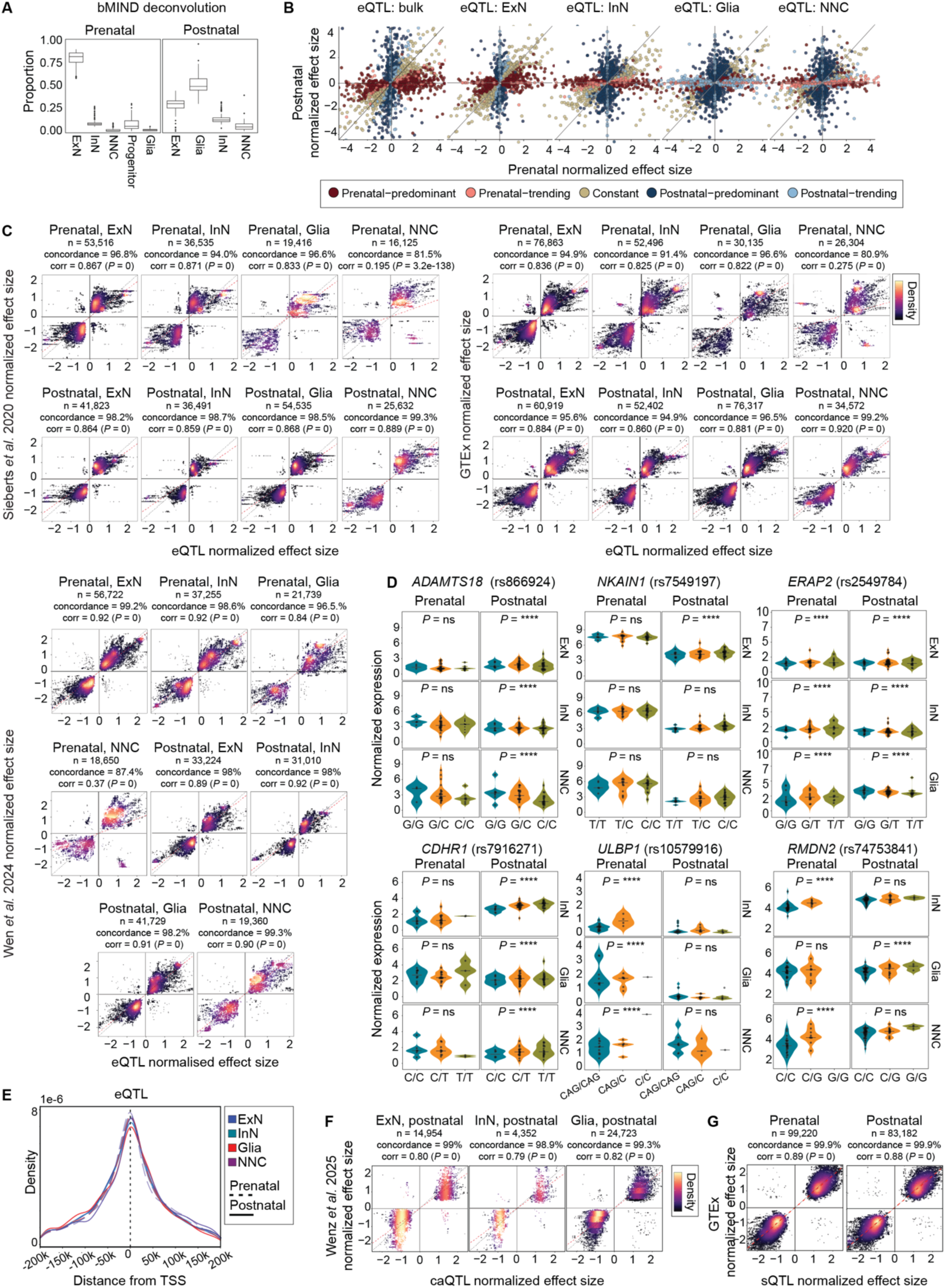
eQTL characterization and replication across public datasets and published studies related to Figure 3. **(A)** bMIND deconvolution of prenatal and postnatal bulk RNA-seq samples showing inferred proportions of excitatory neurons (ExN), inhibitory neurons (InN), glial cells, progenitors, and non-neuronal cells (NNC). Boxplots represent donor-level distributions for each developmental stage (see also **Table S14**). **(B)** Comparison of prenatal and postnatal eQTL effect sizes across bulk tissue and individual cell types. eQTLs are colored according to developmental specificity categories: prenatal-predominant, prenatal-trending, constant, postnatal-predominant, and postnatal-trending (see also **Table S15**). **(C)** Validation of significant eQTLs (*p* < e-5) by comparison of effect sizes with independent datasets from previous studies.^9,16,17,34^ Density scatterplots show concordance of effect directions and Spearman correlation coefficients between datasets. Effect size from each dataset were independently standardized by z-score normalization prior to correlation analysis. Color intensity represents local point density, where warmer colors indicate regions with higher concentrations of overlapping eQTLs. n denotes the number of overlapping eQTLs between datasets. The GTEx^16^ and Sieberts et al.^17^ datasets were derived from adult prefrontal cortex tissue, whereas the Wen et al.^9^ dataset was derived from prenatal prefrontal cortex tissue. **(D)** Representative examples of developmental-stage- and cell-type-predominant eQTLs related to Figure 3C. Violin plots show raw normalized gene expression (log_2_CPM+1; not adjusted for covariates) stratified by genotype across prenatal and postnatal cell populations. Genotypes are shown as homozygous reference (blue), heterozygous (yellow), and homozygous alternate (green). The *P* shown on plot is nominal significant score calculated by TensorQTL pipeline. **(E)** Distribution of distances between eQTL variants and TSSs across broad cell types and developmental stages. **(F)** Replication of postnatal ExN, InN, and Glia caQTL effect sizes with Wenz et al*.-*derived caQTL dataset.^35^ Density scatterplots show concordance rates and Spearman correlation coefficients between datasets. Effect size from each dataset were independently standardized by z-score normalization prior to correlation analysis. **(G)** Replication of prenatal and postnatal sQTL effect sizes with GTEx splicing QTL dataset for human prefrontal cortex tissue.^34^ Density scatterplots show concordance of effect directions and Spearman correlation coefficients between datasets. All genomic coordinates correspond to the hg38 human reference genome assembly. Effect size from each dataset were independently standardized by z-score normalization prior to correlation analysis. **Abbreviations:** bMIND, Bayesian model for inferring cell-type-specific gene expression; caQTLs, chromatin accessibility quantitative trait loci; CPM, counts per million; eQTLs, expression quantitative trait loci; GTEx, Genotype-Tissue Expression; sQTLs, splicing quantitative trait loci; TSSs, transcription start sites.

**Figure S4.**
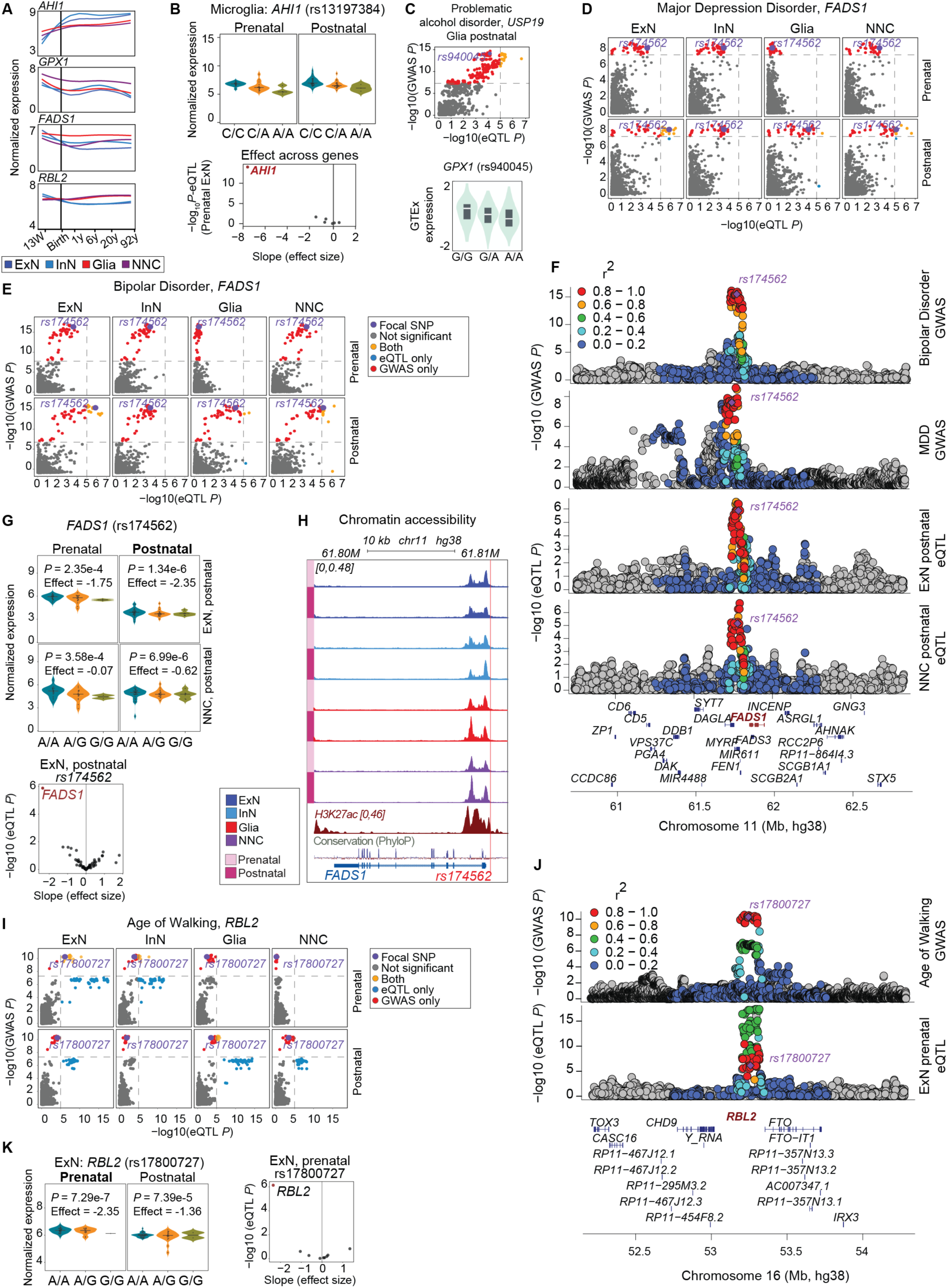
Developmental expression trajectories and additional colocalization signals across cell types and developmental stages related to Figure 4. **(A)** Smoothed log_2_CPM+1-normalized expression of *AHI1*, *GPX1*, *FADS1*, and *RBL2* across broad cell types and developmental periods. **(B)** Up: Genotype-stratified *AHI1* expression for the candidate causal variant rs13197384 in prenatal and postnatal Microglia. Violin plots show raw normalized expression (log_2_CPM+1; not adjusted for covariates) across genotypes. Genotypes are shown as homozygous reference (blue), heterozygous (yellow), and homozygous alternate (green). Bottom: eQTL effect sizes and *p* values for all genes associated with rs13197384 within the prenatal excitatory *AHI1* locus, demonstrating that *AHI1* is the primary significantly affected gene with the strongest effect size. Each point represents an individual gene. **(C)** Scatterplots of problematic alcohol disorder GWAS variants at the *USP19* locus with eQTLs across glia postnatal. Variants are colored according to GWAS/eQTL significance overlap. The candidate causal variant according to Figure 4H is highlighted in purple. **(D)** Scatterplots showing colocalization of MDD GWAS variants at the *FADS1* locus with eQTLs across broad cell types and developmental stages. Variants are colored according to GWAS/eQTL significance overlap. The candidate causal variant is highlighted in purple. **(E)** Scatterplots showing colocalization of BIP GWAS variants at the *FADS1* locus with cis-eQTLs across cortical cell types and developmental stages. Variants are colored according to GWAS/eQTL significance overlap. The candidate causal variant is highlighted in purple. **(F)** LocusZoom plots showing MDD GWAS, BIP GWAS, and excitatory neuronal and non-neuronal eQTL association signals at the *FADS1* locus. Variants are colored according to linkage disequilibrium (LD) (r²) with the lead SNP rs174562.| **(G)** Genotype-stratified *FADS1* expression for rs174562 across prenatal and postnatal excitatory and non-neuronal cell populations. Violin plots show raw normalized expression (log_2_CPM+1; not adjusted for covariates) across genotypes. *P* values and effect sizes were calculated using TensorQTL. Bottom: eQTL effect sizes and *p* values for all genes associated with rs174562 within the *FADS1* locus. Each point represents an individual gene. **(H)** Chromatin accessibility profiles across developmental stages and cortical cell types at the *FADS1* locus. Track shows pseudobulk ATAC-seq accessibility (ReadsInTSS normalized). Candidate causal variant is highlighted. **(I)** Scatterplots showing colocalization of AOW GWAS variants at the *RBL2* locus with eQTLs across cell types and developmental stages. Variants are colored according to GWAS/eQTL significance overlap. The candidate causal variant is highlighted in purple. **(J)** LocusZoom plot showing AOW GWAS and prenatal excitatory neuron eQTL association signals at the *RBL2* locus. Variants are colored according to LD (r²) with the lead SNP rs17800727. **(K)** Left: Genotype-stratified *RBL2* expression for rs17800727 across prenatal and postnatal excitatory neurons. Violin plots show raw normalized expression (log_2_CPM+1; not adjusted for covariates) across genotypes. *P* values and effect sizes were calculated using TensorQTL. Right: eQTL effect sizes and *p* values for all genes associated with rs17800727 within the *RBL2* locus. Each point represents an individual gene. The genome-wide significance threshold for GWAS associations is p < 5e-8 and the significance threshold for eQTL associations is p < 1e-5. All genomic coordinates correspond to the hg38 human reference genome assembly. Also see Figure 4 and BrainVar website (www.brainvar.org/). **Abbreviations:** AOW, age of onset of weekly drinking; BIP, bipolar disorder; CPM, counts per million; eQTLs, expression quantitative trait loci; LD, linkage disequilibrium; MDD, major depressive disorder

**Figure S5.**
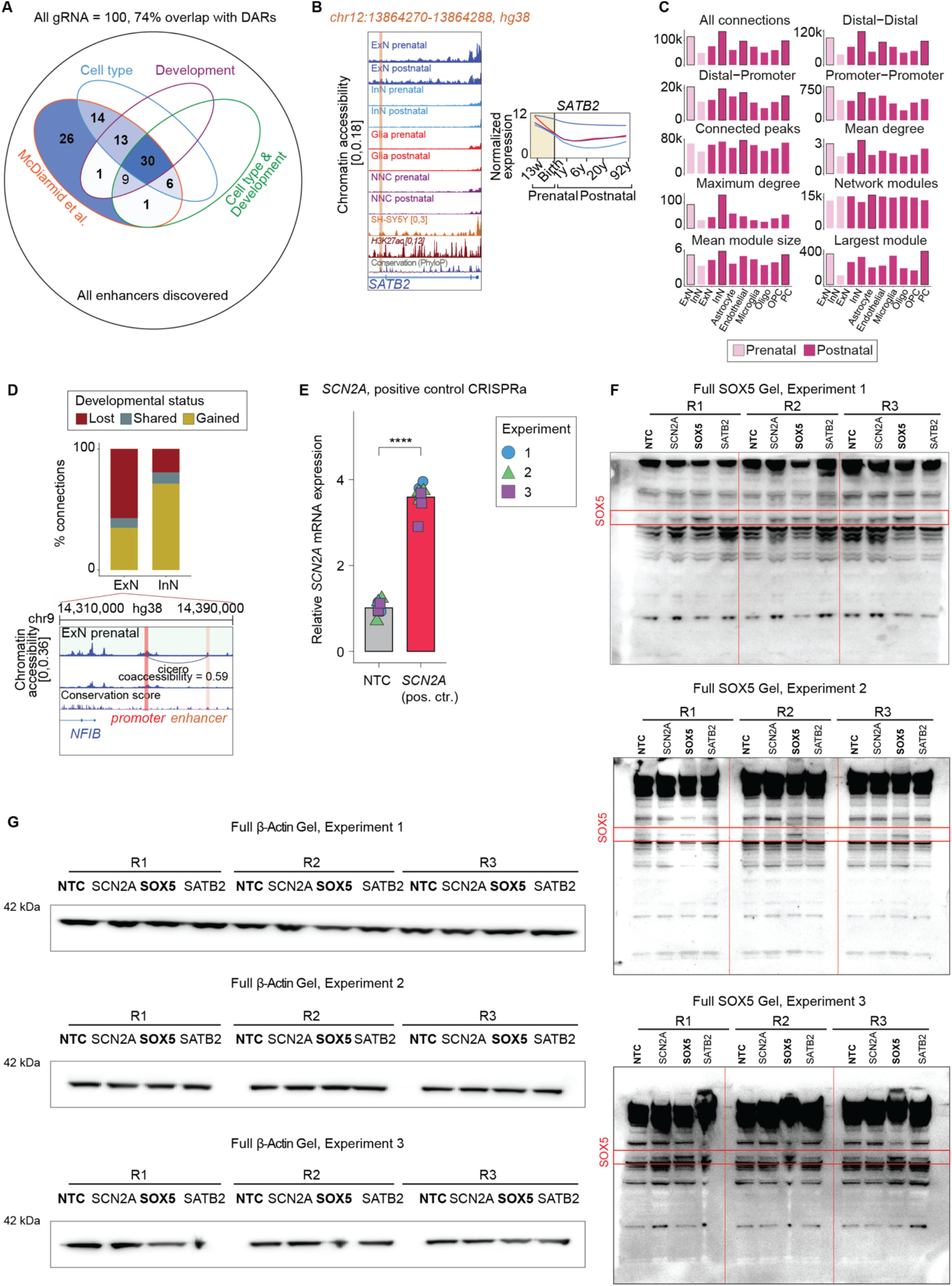
Coaccessibility analysis and enhancer discovery related to Figure 5. **(A)** Overlap between discovered DARs and enhancers identified based on ATAC data with McDarmid et al. study targeting 100 enhancers associated with NDD haploinsufficient genes. **(B)** Example of the overlapped DARs with McDarmid et al.^53^ study gRNA for *SATB2.* Normalized chromatin accessibility (ReadsInTSS normalized) profiles are shown for prenatal and postnatal excitatory neurons, inhibitory neurons, glia, and non-neuronal cell populations, together with H3K27ac and chromatin accessibility profiles from SH-SY5Y cells. The *SATB2* promoter and the enhancer at chr12:13864270–13864288 (hg38) are highlighted. Conservation score shows phyloP conservation visualized using UCSC genome browser. Right: Smoothed normalized (log_2_CPM+1) *SATB2* expression trajectories in excitatory neurons, inhibitory neurons, glia, and non-neuronal cells. **(C)** Statistics of co-accessibility connections and networks identified by Cicero for each developmental stage and cell type. “All connections” indicates the total number of co-accessibility connections; “distal-to-distal” indicates the number of connections between two distal peaks; “distal-to-promoter” indicates the number of connections between distal and promoter regions; and “promoter-to-promoter” indicates the number of connections between two promoter regions. Additional statistics include the number of unique connected peaks, the mean and maximum degrees of the co-accessibility networks, the number of network modules, the mean module size, and the size of the largest module. **(D)** Percentage of co-accessibility connections lost, shared, or gained during development in excitatory and inhibitory neurons. The *NFIB* locus is shown as an example in excitatory neurons, illustrating a prenatal-specific connection that is lost postnatally. **(E)** CRISPRa targeting of the *SCN2A* positive control region increases *SCN2A* mRNA expression (qPCR) relative to the NTC condition in SH-SY5Y cells. Points denote measurements from three independent experiments, and bars summarize relative *SCN2A* expression. **(F)** Full *SOX5* immunoblot for Experiment 1, 2, and 3. *SOX5* protein levels were assessed under non-targeting control (NTC), SCN2A, and *SOX5* enhancer-targeting CRISPRa conditions. A *SATB2*-targeting gRNA from McDarmid et al.^53^ was also included in the immunoblot; however, our qPCR analysis did not show statistically significant upregulation of *SATB2* in SH-SY5Y cells in all experiments. **(G)** Full β-actin gel images for all experiments and biological replicates, showing consistent protein loading across all samples. **Abbreviations:** CPM, counts per million; CRISPRa, CRISPR activation; DARs, differentially accessible regions; NDD, neurodevelopmental disorder; NTC, non-targeting control; qPCR, quantitative polymerase chain reaction

## RESOURCE AVAILABILITY

### Lead contact

Requests for further information and resources should be directed to and will be fulfilled by the lead contact, Stephan Sanders.

### Materials availability

The unique reagent generated in this study for SOX5-targeting CRISPR activation (CRISPRa) guide RNA and associated plasmid constructs, are available from the lead contact upon reasonable request. No additional unique biological materials were generated in this study.

### Data and code availability

#### Data

Raw sequencing data generated in this study, including snRNA-seq and snATAC-seq, bulk RNA-seq, and whole-genome sequencing, together with large-scale processed datasets, including structural variant calls, eQTLs, caQTLs, sQTLs, and chromatin co-accessibility results, will be available through the NIMH Data Archive (NDA; https://nda.nih.gov/) under the BrainVar collection as of the date of publication, subject to applicable data-access requirements.

Processed snRNA-seq and ATAC-seq HDF5 objects, including gene expression and chromatin accessibility matrices and cell-type annotations, and genome-wide chromatin accessibility tracks (bigWig files) are publicly available through the UCSC Cell (brainvar.cells.ucsc.edu) and Genome Browser (https://genome.ucsc.edu/cgi-bin/hgTrackUi?db=hg38&g=singleCellSignalsPeaks). An interactive platform for exploring eQTL associations and GWAS–eQTL colocalization results are available at https://brainvar.org. Additional processed datasets and supporting results are provided in the supplemental information accompanying this paper.

This study also reanalyzes previously published human brain transcriptomic, genomic, and QTL datasets, including those from Werling et al. (2020),^7^ Trevino et al. (2021),^57^ Ma et al. (2022),^73^ Herring et al. (2022),^26^ Wang et al. (2025),^74^ the GTEx Consortium,^16^ Sieberts et al. (2020),^17^ Wen et al. (2024),^9^ and Wenz et al. (2025).^35^ Previously published GWAS summary statistics, rare-variant gene sets, and functional genomic resources, including enhancer–promoter predictions and ABC scores, were also analyzed. All external datasets and their corresponding source references, accession numbers, and identifiers are listed in **Key Resources Table**.

#### Code

All original code used for data processing, computational analyses, statistical testing, and figure generation is publicly available through GitHub at https://github.com/sanderslab/brainvar2-public/ and https://github.com/SestanLab/BrainVar

#### Additional information

Any additional information required to reanalyze the data reported in this paper is available from the lead contact upon request.

## ACKNOWLEDGMENTS

The authors thank Xiaozheng Yao, Brian Park, Kai-Ling Chen and Mitchell J. Machiela for assistance with LDlink concurrent access. LDlink is a publicly available webtool hosted at ldlink.nih.gov by the Integrative Tumor Epidemiology Branch of the Division of Cancer Epidemiology and Genetics and maintained by the Center for Biomedical Informatics and Information Technology. LDlink is funded by the intramural research program of National Cancer Institute within the US National Institutes of Health. We also thank the UCSC Cell Browser team for developing and maintaining the Cell Browser, which facilitate data visualization and exploration. We are deeply grateful to the brain donors and their families whose generous tissue donations made this research possible. Some illustrations in the figures were created with BioRender.

## AUTHOR CONTRIBUTIONS

Conceptualization, N.R., S.J.S., Y.L., R.K., and N.S.; Experimental design: N.R., R.K., Y.L., T.M., S.P., K.R., B.D., N.S., C.R., A.G.-G, and S.J.S.; Functional assays and experimental validation: A.G.-G., T.M., S.P., S.C., J.K., R.M., and C.R.; Data generation: R.K., S.P., S.C., J.K., A.D.-F., A.-N.S., H.A.-B., and M.J.G.; Data processing: N.R., R.K., Y.L., E.M.W., J.L.L., S.D., A.N., A.D.-F., M.G., K.J., and X.Z.; Data analysis: N.R., Y.L., S.-W.K., Y.K., Q.X., E.M.W., J.W., J.L.L., S.D., A.N., L.K., X.W., H.C., M.L., D.L., A.L., S.M., M.Z., K.J., and X.Z.; Statistical analysis: N.R., Y.L., S.-W.K., Y.K., J.W., L.K., X.W., K.R., and B.D.; Clinical data and interpretation: N.R., R.K., P.L.D.J., K.R., B.D., N.S., and S.J.S.; Scientific and technical support: D.S.-P., G.G., P.K., M.G., N.F.P., and C.J.Y.; Visualization, N.R.; Writing – original draft, N.R.; Project administration, Y.L., R.K., and N.R.; Funding acquisition, S.J.S and N.S.; Supervision and scientific oversight: S.J.S, N.S., H.B., M.E.T., D.M.W., C.R., K.R., and B.D.

N.R., R.K., Y.L., N.S. and S.J.S. jointly led the BrainVar project. N.R. coordinated the project, designed and supervised computational analyses, interpreted results, prepared figures, and led manuscript preparation. R.K. led BrainVar experimental data generation, metadata preparation, and manuscript preparation. Y.L. contributed to study design, quality control, computational analyses, and metadata and manuscript preparation. S.J.S. provided overall scientific leadership, directed the study, supervised the analytical strategy, and oversaw manuscript development. N.S. directed the experimental data-generation, supervised experimental research, and contributed to study design and interpretation. Manuscript preparation: All authors contributed to writing, reviewing, and editing the manuscript and approved the final version.

## DECLARATION OF GENERATIVE AI AND AI-ASSISTED TECHNOLOGIES

During the preparation of this work, the authors used ChatGPT (OpenAI) and Claude AI (Anthropic) to assist with code refinement and manuscript grammar and writing editing. After using these tools, the authors reviewed and edited the content as needed and take full responsibility for the content of the publication.

## SUPPLEMENTAL INFORMATION

### Supplementary Tables

**Table S1.** Donor metadata and analysis inclusion information, related to **Figure 1**

**Table S2.** Sequencing library metadata, related to **Figure 1**

**Table S3.** Single-nucleus multiome cell metadata, related to **Figure 1**

**Table S4.** Motif discovery results for the differentially accessible regions, related to **Figure 1** and **Figure S1**

**Table S5.** Pathway enrichment for the differentially expressed genes, related to **Figure 1** and **Figure S1**

**Table S6.** LOEUF enrichment results for the differentially expressed gene, related to **Figure 1** and **Figure S1**

**Table S7.** GWAS summary statistics information for traits, related to **Figure 2** and **Figure S2**

**Table S8.** FGSEA enrichment results across cell types and developmental stages, related to **Figure 2** and **Figure S2**

**Table S9.** ASD gene functional classification, related to **Figure 2**

**Table S10.** ASD, NDD, MDD, and SCZ gene functional classification, related to **Figure 2** and **Figure S2**

**Table S11.** LDSC partitioned heritability enrichment results, related to **Figure 2**

**Table S12.** Pathway enrichment and STRING network results, related to **Figure 2**

**Table S13.** NMF results and inputs for shared biological domains, related to **Figure 2**

**Table S14.** Marker genes used for deconvolution using bMIND, related to **Figure S3**

**Table S15.** eQTL temporal category classification, related to **Figure 3**

**Table S16.** Motif discovery results for the caQTL peaks, related to **Figure 3**

**Table S17.** Overlap of caQTL peaks with differentially accessible regions, related to **Figure 3**

**Table S18.** Overlap of caQTL results with eQTL results, related to **Figure 3**

**Table S19.** Colocalization results and highlighted loci, related to **Figure 4** and **Figure S4**

**Table S20.** Gene regulatory network results, related to **Figure 4** and **Figure S4**

**Table S21.** GWAS enrichment results for GRN of *AHI1* candidate causal variant in prenatal excitatory neurons, related to **Figure 4**

**Table S22.** MotifbreakR analysis results for AHI1 candidate causal variant, related to **Figure 4**

**Table S23.** Cellchat results for *AHI1* candidate causal variant, related to **Figure 4**

**Table S24.** List of CREs for NDD haploinsufficient genes, related to **Figure 5** and **Figure S5**

**Table S25.** qPCR results for CRISPR-a experiments targeting the identified *SOX5* enhancer, related to **Figure 5**

**Table S26** Quantified protein expression for CRISPR-a experiments targeting the identified *SOX5* enhancer, related to **Figure 5** and **Figure S5**

## Supplementary Data

**Data S1.** Cell-type marker DEGs and DARs, related to **Figure 1** and **Figure S1**

**Data S2.** Developmental marker DARs and DEGs, related to **Figure 1** and **Figure S1**

**Data S3.** Joint developmental and cell-type marker DARs and DEGs, related to **Figure 1** and **Figure S1**

**Data S4.** DARs based on gene activity scores, related to **Figure 2**

**Data S5.** MAGMA gene-level results for GWAS traits, related **to Figure 2** and **Figure S1**

## Supplementary Documents

**Document S1.** CNV analysis results related to **Figure 1**.

## STAR★METHODS

### KEY RESOURCES TABLE

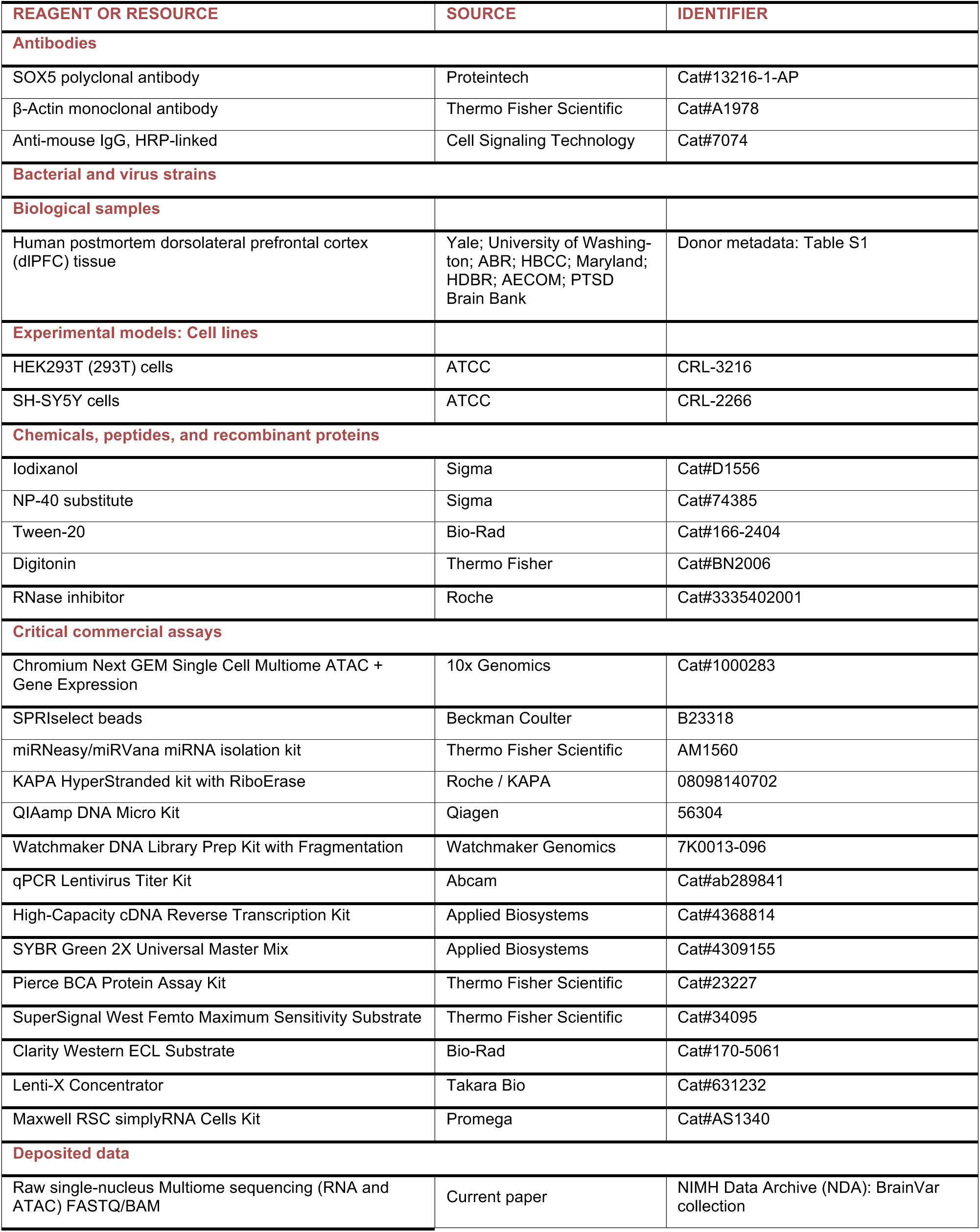
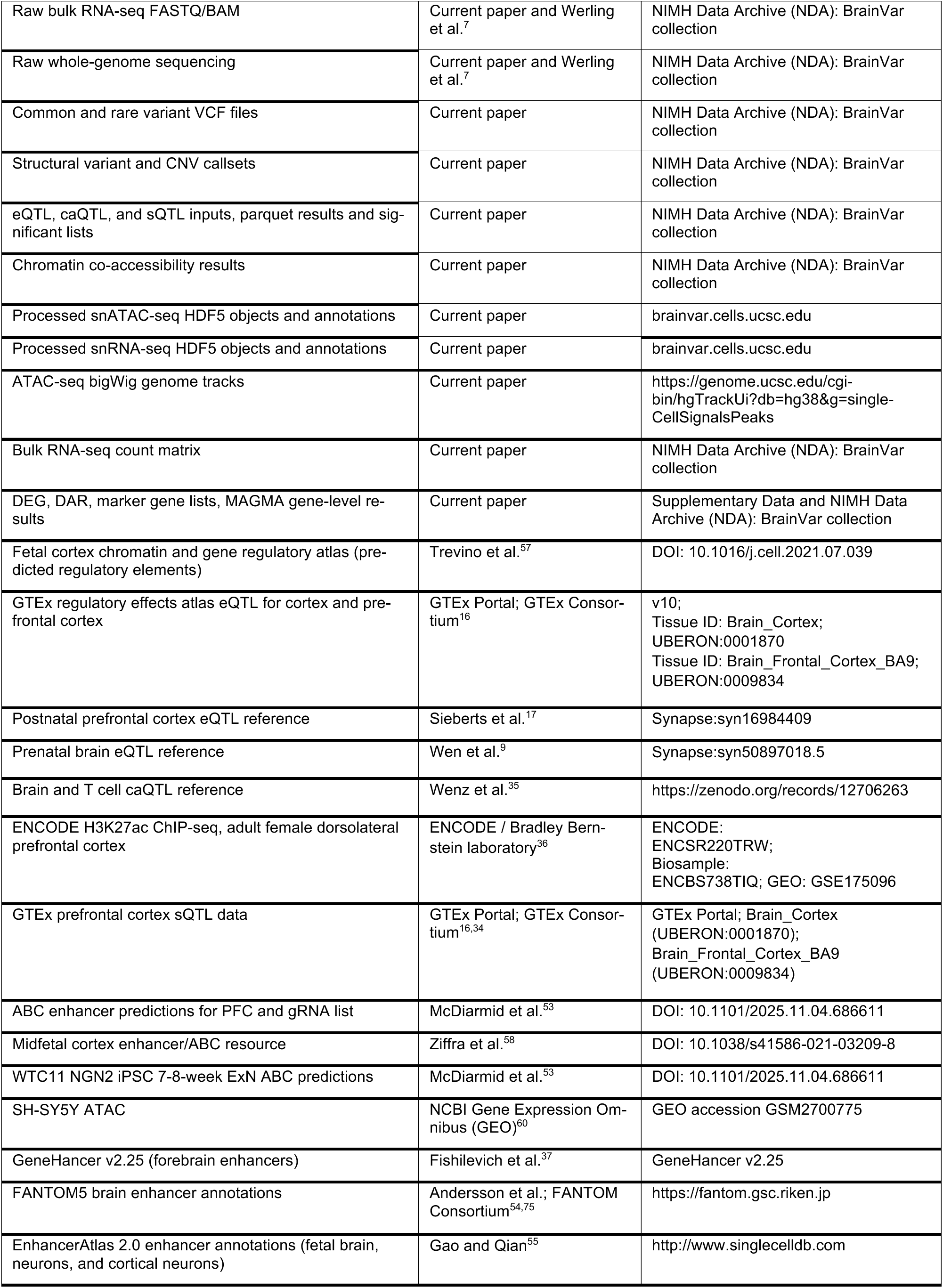
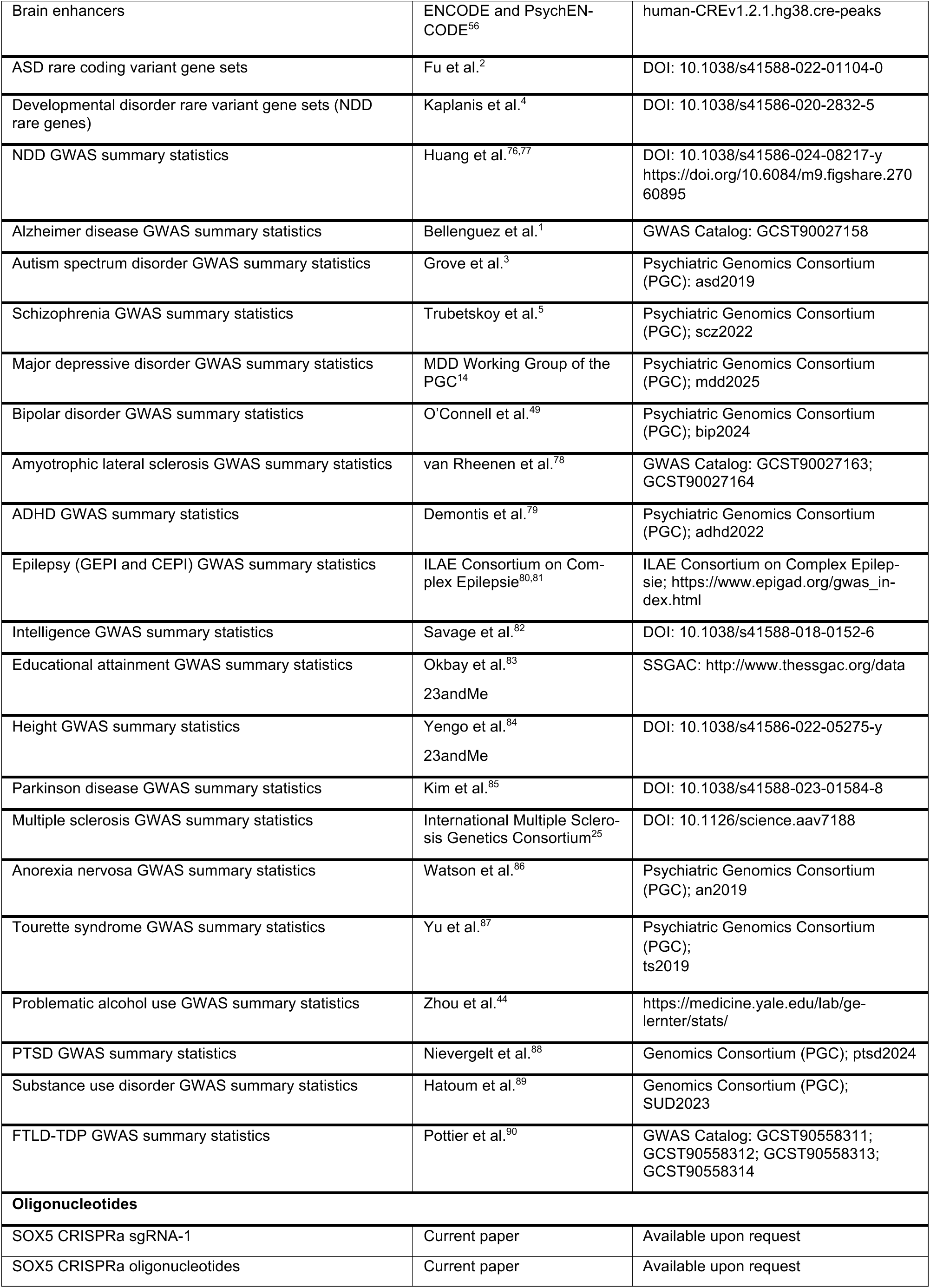
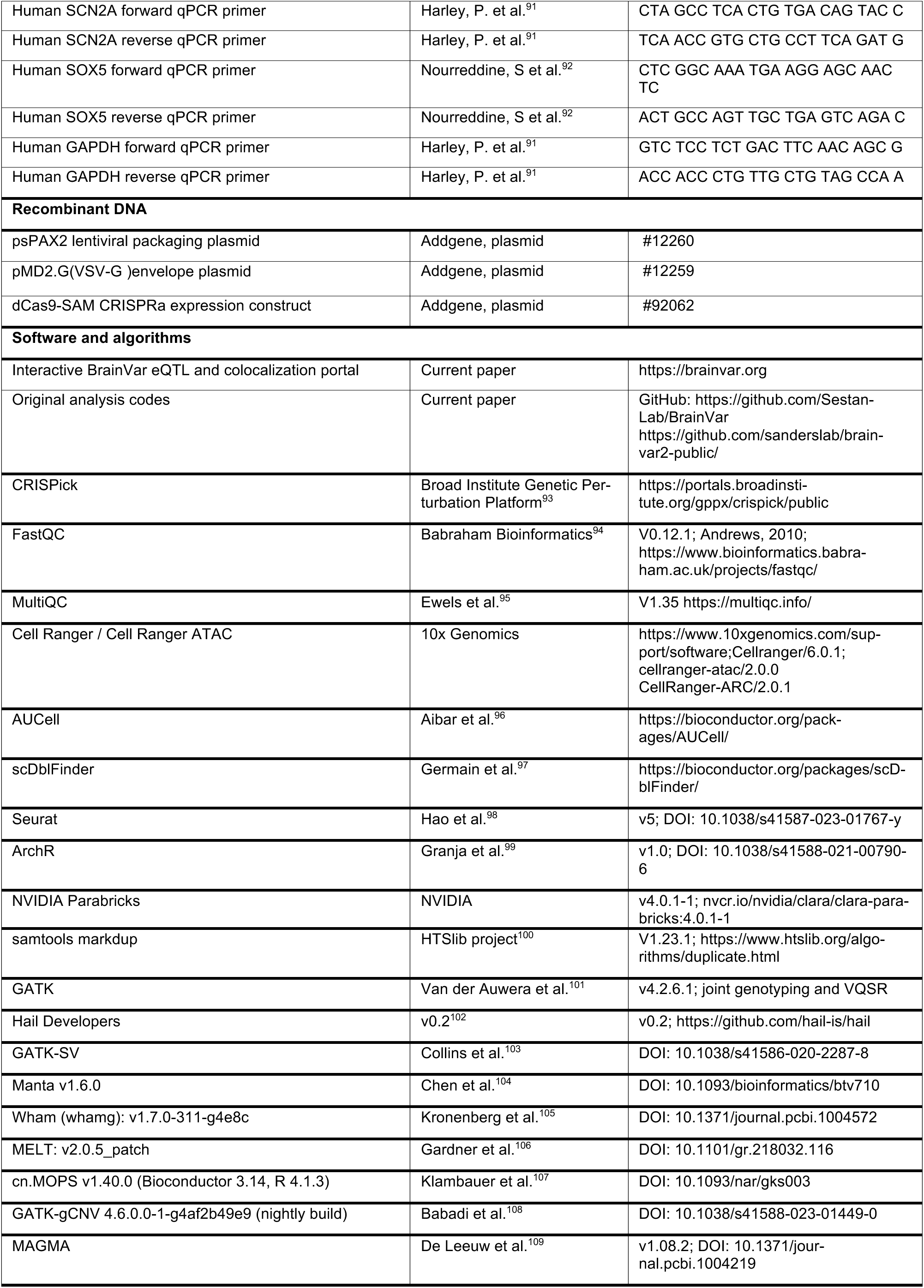
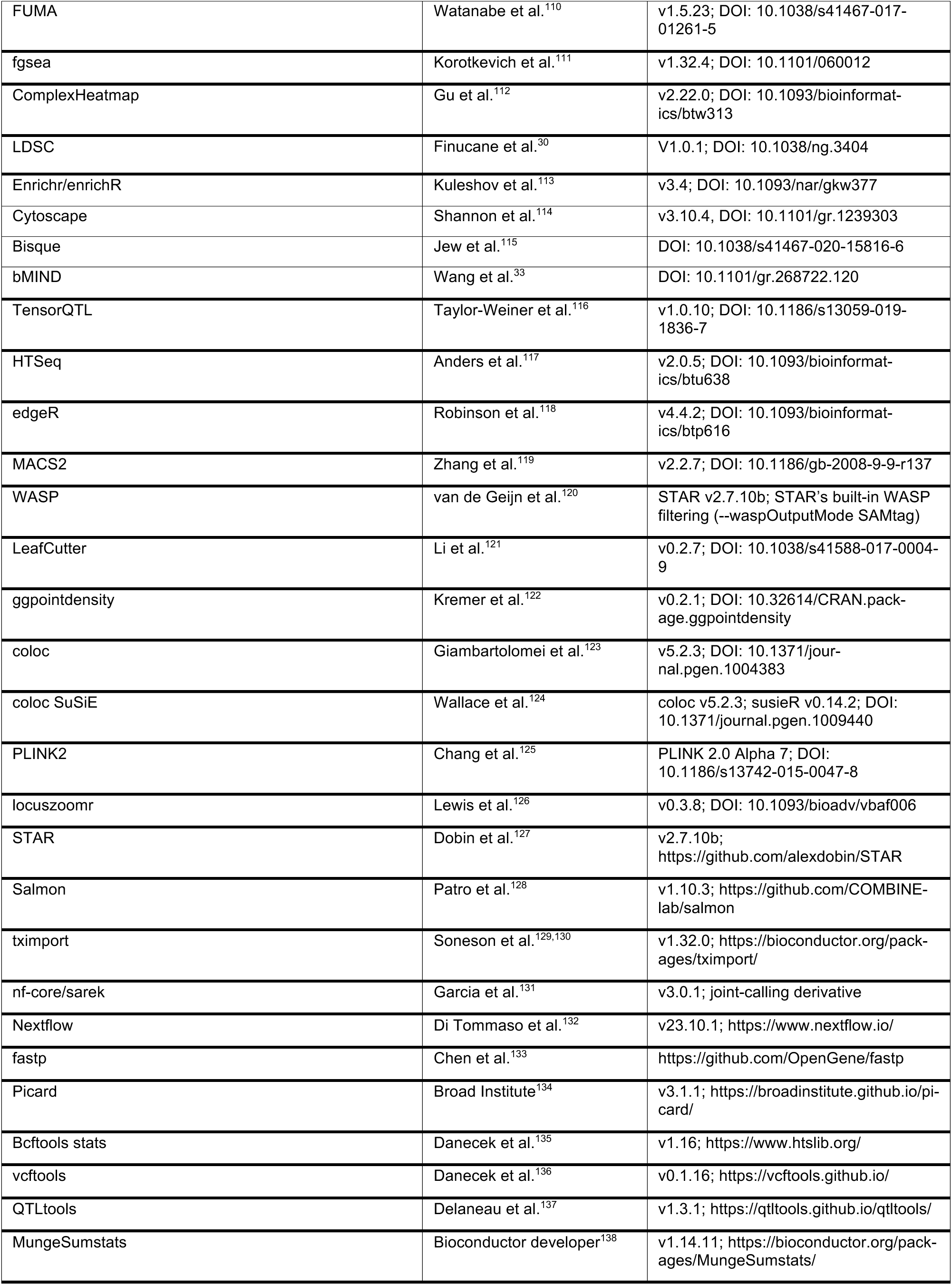
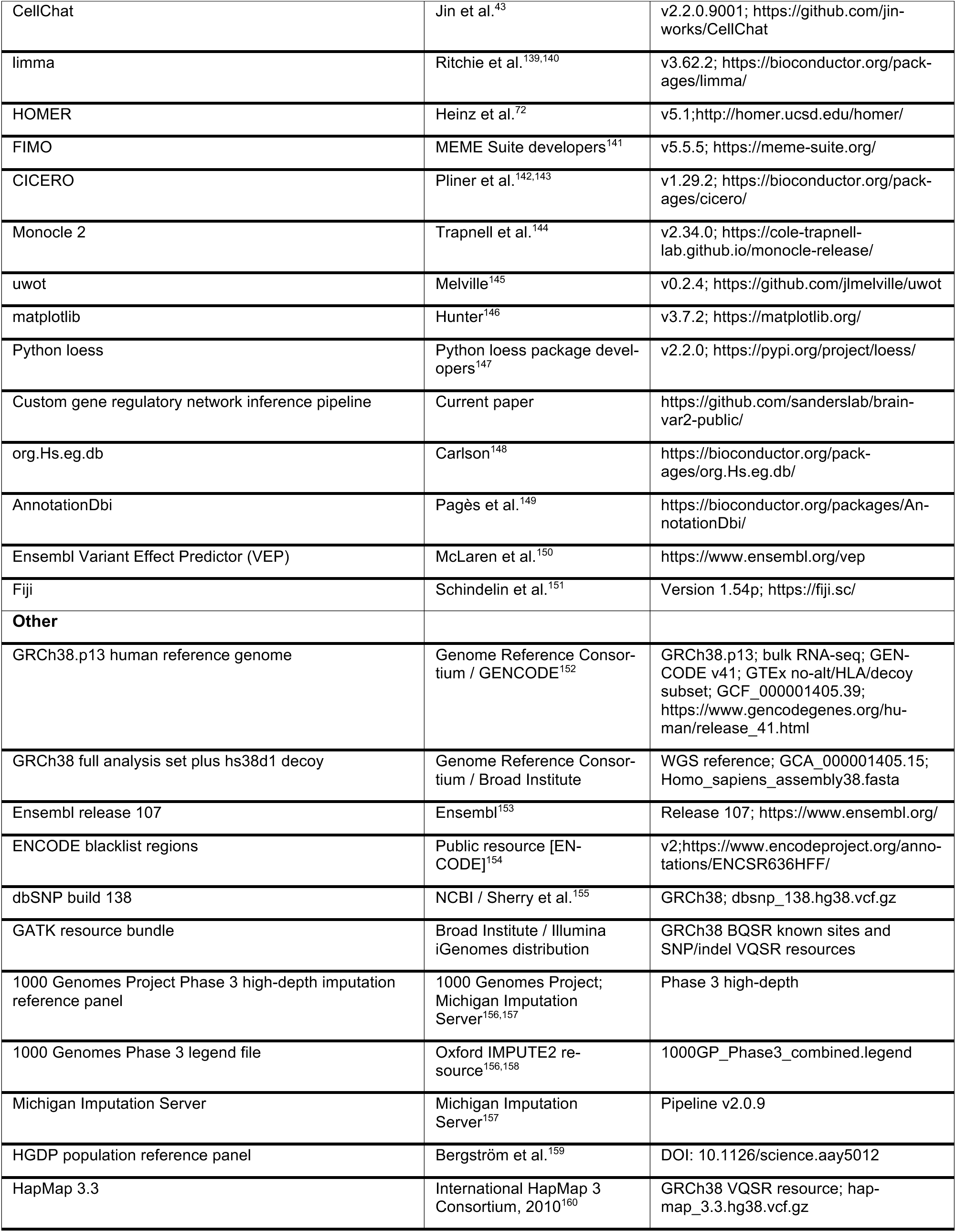
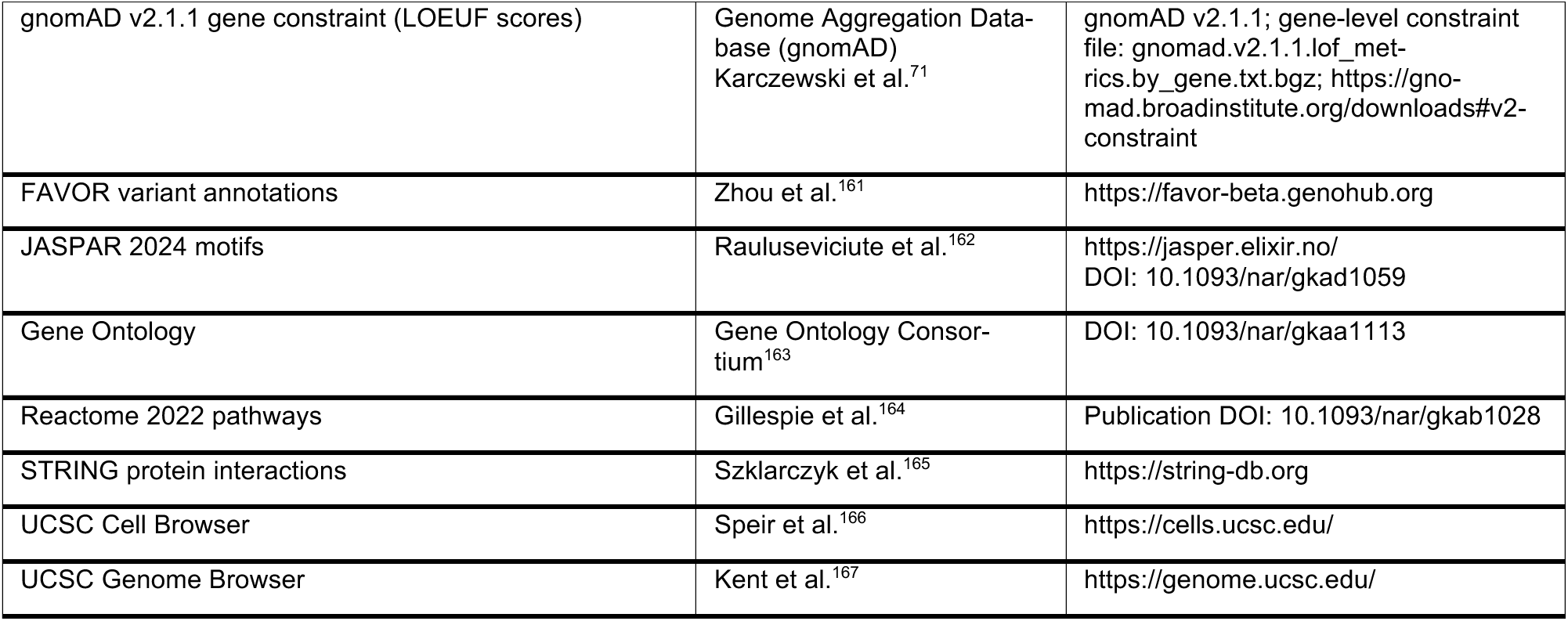

## EXPERIMENTAL MODEL AND STUDY PARTICIPANT DETAILS

### Human postmortem tissue

Postmortem human brain specimens were obtained from multiple established tissue repositories, including the Department of Neuroscience at Yale University School of Medicine, the Birth Defects Research Laboratory at the University of Washington, Advanced Bioscience Resources Inc. (ABR), the Human Brain Collection Core (HBCC), the Brain and Tissue Bank at the University of Maryland, the MRC–Wellcome Trust Human Developmental Biology Resource at the University of Newcastle (UK), the Human Fetal Tissue Repository at Albert Einstein College of Medicine (AECOM), and the National Center for PTSD Brain Bank. All tissues were collected following informed consent from parents or next of kin and with approval from the institutional review boards at Yale University School of Medicine, the National Institutes of Health, and the respective contributing institutions. All procedures complied with ethical standards for research involving human tissue, in accordance with National Institutes of Health (NIH) guidelines and the World Medical Association (WMA) Declaration of Helsinki.^168^

In total, 118 postmortem brain specimens (female = 46, male = 71, unconfirmed = 1) were used for 10x Genomics snMultiome profiling, and 240 specimens (female = 95, male = 139, unconfirmed = 8) were used for bulk RNA sequencing. WGS was performed for 262 donors (female = 104, male = 150, unconfirmed = 8), including donors previously published.^7^ Donor ages spanned 15 developmental stages, ranging from 6 PCW to 92 years, as defined by established developmental frameworks.^18^ Post-conception age was calculated as gestational age in weeks minus two weeks. Postmortem interval (PMI) was defined as the time in hours between death and tissue freezing. After quality control and preprocessing, a total of 231 donors were included in the analysis, of whom 114 had 10x multiome data, 229 had bulk RNA-seq data, and 231 had WGS data.

## METHOD DETAILS

### Tissue dissection and processing

Tissue dissection was performed as previously described.^18^ The dlPFC (Brodmann areas 9/46) was isolated based on established anatomical landmarks. For specimens at 9–10 PCW, dissections encompassed the full thickness of the cerebral wall. Samples from 12–22 PCW primarily included the cortical plate and, in some cases, portions of the underlying subplate. For specimens spanning 28 PCW to 92 postnatal years, dissections were performed to capture the entire cortical gray matter (layers 1-6); in samples from 4–12 postnatal months, this additionally included portions of the subplate. The majority of samples used for snMultiome profiling overlap with those previously reported,^7^ with additional samples added across developmental stages. Period 1 samples were not included in the snMultiome dataset due to the lack of available tissue. Period 2 donors (3 donors) were excluded because they were determined to contain predominantly noncortical cells, indicating inaccurate regional dissection (see **Table S1**). Therefore, snMultiome libraries of period 2 were not included in the following processing steps.

### Nuclear isolation, multiome capture, library generation, and sequencing

Nuclei were isolated using a protocol adapted from our previous work.^73^ To minimize experimental bias, frozen brain tissue was pulverized into a fine powder in liquid nitrogen using a mortar and pestle (Coorstek #60316, #60317) for samples in which the dlPFC region weighed >40 mg. For tissues weighing <40 mg, samples were sectioned on a cryostat and leaflets approximately 100– 300 µm thick were collected, or intact tissue fragments were used. Pulverized material was aliquoted and weighed (10–30 mg) prior to the day of nuclei isolation. All buffers were prepared using molecular-grade reagents unless otherwise specified and maintained at 4°C until use. All plastic consumables and glass Dounce homogenizers were RNase-free. Samples were processed in batches, with each experimenter handling 1–3 samples per experiment.

For lysis, 1 mL of lysis buffer [250 mM sucrose (Sigma, S0389), 25 mM KCl (Sigma, 60142), 5 mM MgCl₂ (Sigma, M1028), 20 mM Tris-HCl pH 7.5 (Invitrogen, 15567-027), 0.1% Igepal (Sigma, I8896), 1× protease inhibitor (Roche, 11836170001 or 5056489001), 1 mM Dithiothreitol (DTT: Sigma, 43816), and 0.08 U/µL RNase inhibitor (Roche, 3335402001) was added directly to each tissue aliquot and vortexed. The suspension was transferred to a pre-chilled Dounce homogenizer on wet ice containing 1 mL of lysis buffer. An additional 1 mL of lysis buffer was added to the original tube, vortexed to recover residual tissue, and transferred to the homogenizer. Tissue was homogenized using loose and tight pestles (30–50 strokes each) with consistent pressure and without introducing air bubbles. The homogenate was filtered through a 40 µm tube-top strainer pre-wetted with 250 µL lysis buffer. For samples from donors <20 years of age, the filtrate was transferred to a 15 mL tube containing 3 mL gradient buffer [25 mM KCl, 5 mM MgCl₂, 20 mM Tris-HCl pH 7.5, 50% iodixanol (Sigma, D1556), 1% Bovine Serum Albumin (BSA) with 10× protease inhibitor (premade stock), 1 mM DTT, and 0.08 U/µL RNase inhibitor] and mixed by gentle inversion (10 times). These samples were centrifuged at 1,000 × g for 30 min at 4°C in a swinging-bucket rotor. After centrifugation, debris was removed by vacuum, and the supernatant was care-fully aspirated, leaving ≤50 µL above the nuclei pellet. Samples from donors >20 years of age (**Table S1**) were processed in 2 mL tubes. Samples were resuspended in 0.4 mL lysis buffer and combined with 0.4 mL 50% iodixanol buffer [50% (v/v) iodixanol, 25 mM KCl, 5 mM MgCl₂, 20 mM Tricine-KOH (pH 7.8), Ethylenediaminetetraacetic Acid (EDTA) free protease inhibitors, RNase inhibitor (80 U/mL), 1 mM DTT, and water], followed by mixing (10 inversions). The mixture was underlaid with 0.6 mL of 40% iodixanol cushion (prepared by mixing lysis buffer and iodixanol buffer) and overlaid with 0.6 mL of 30% iodixanol (prepared similarly), with the sample suspension layered on top. Tubes were centrifuged at 3,000 × g for 30 min at 4°C without braking. Following centrifugation, the 30%/40% interphase was collected into a new tube.

Nuclei pellets or interphase fractions were resuspended by pipette mixing in 0.5 mL nuclear permeabilization buffer [10 mM NaCl, 3 mM MgCl₂, 10 mM Tris-HCl pH 7.5, 0.01% NP-40 substitute (Sigma, 74385), 0.01% Tween-20 (Bio-Rad, 166-2404), 0.001% digitonin (Thermo Fisher, BN2006), 1× protease inhibitor, 1 mM DTT, 1% BSA, and 1 U/µL RNase inhibitor] and incubated on ice for 5 min. To dilute the permeabilization buffer, 0.5 mL wash buffer (10 mM NaCl, 3 mM MgCl₂, 10 mM Tris-HCl pH 7.5, 0.1% Tween-20, 1 U/µL RNase inhibitor, 1 mM DTT, and 1% BSA) was added, and nuclei were filtered through a 30 µm cell strainer into a new 1.5 mL tube. An aliquot of the nuclei suspension was removed for counting, and the remaining sample was centrifuged at 500 × g for 10 min at 4°C. During centrifugation, nuclei were counted using a manual hemocytometer at 20× magnification, with samples diluted 1:2 (postnatal) or 1:4 (prenatal). If centrifugation completed during counting, one experimenter proceeded with supernatant removal while another completed counting. Based on calculated concentrations, nuclei were resuspended in resuspension buffer [1× Nuclei Buffer (10x Genomics ATAC Kit A), 1 mM DTT, and 1 U/µL RNase inhibitor] to achieve ∼8–10 million nuclei/mL.

Following resuspension, samples were filtered through a 40 µm cell strainer pre-wetted with resuspension buffer. A second aliquot was counted to confirm a final concentration of 3.3–8 million nuclei/mL, consistent with 10x Genomics Gene expression (GEX) and ATAC guidelines (CG000338) for targeting 10,000 nuclei. Single nuclei were subjected to transposition, capture, and library preparation according to the 10x Genomics Multiome protocol. GEX and ATAC libraries were sequenced on an Illumina NovaSeq 6000 at the Yale Center for Genomic Research to an initial depth of 250 million reads. Following initial quality control, libraries flagged for insufficient depth were re-sequenced to achieve a minimum of 25,000 reads per cell.

### Sample ID naming Terminology

For each donor, a region of the dlPFC was dissected and subdivided into multiple aliquots (tubes), each designated for distinct assays. Donor identifiers typically begin with the prefix human subject brain (HSB), for example, HSB107. Biospecimens used for bulk RNA sequencing are labeled using the numeric donor ID (excluding the HSB prefix), followed by R1 or R2 to indicate the first or second aliquot used for RNA extraction and sequencing (e.g., 107_R1, 107_R2). In some cases, additional bulk RNA sequencing aliquots were generated due to contamination or donor mismatches identified during downstream analyses (**Table S1, S2**). WGS samples follow a similar convention, with DNA-derived biospecimens labeled using the donor ID and a D suffix (e.g., 107_D1). When discrepancies were detected in downstream analyses, DNA was re-extracted from an alternative biospecimen from the same donor (e.g., 107_D2).

Biospecimens processed for snMultiome assays are labeled with H (Human), followed by the donor ID, brain region (dlPFC), and assay designation (m for multiome), for example, H107DFCm. Most biospecimens were processed once and do not include a numerical suffix. However, when a biospecimen was captured multiple times within the same experiment, identifiers include a suffix (1 or 2) to denote distinct libraries, which are considered technical replicates. In cases where libraries were repeated due to technical issues (e.g., clogging or low-quality output), the initial sample ID does not include a suffix, whereas the repeated sample includes a numerical suffix. These are treated as biological replicates from the same donor. All replicate types can be further distinguished based on capture dates, as detailed in **Table S1**. In **Table S1**, the SubjectID column was filled in only when a match was identified across all modalities (WGS, bulk RNA, snMultiome GEX, and snMultiome ATAC). If the libraries could not be matched, no SubjectID was assigned.

### Bulk RNA extraction, library preparation, and sequencing

The majority of donor samples used for bulk RNA sequencing (RNA-seq) (n = 176) had been previously processed and sequenced as described earlier.^7^ dlPFC tissue samples from 33 extra donors were included to provide bulk RNA-seq data for donors that also underwent snMultiome profiling. For a subset of donors (n = 5), RNA was re-extracted from a separate tissue aliquot and re-sequenced because of either discrepancies identified between RNA and WGS datasets or in-adequate library quality. In addition, RNA from these additional samples was isolated using the miRNeasy/miRVana miRNA Isolation Kit with phenol (Thermo Fisher Scientific, cat# AM1560). Sequencing libraries were generated using the KAPA HyperStranded kit with RiboErase and sequenced at the Yale Center for Genomic Research on the Illumina NovaSeq 6000 platform, achieving an average depth of approximately 43 million reads per library.

### Bulk DNA extraction, library preparation, and sequencing

Most donor DNA samples (n = 219) had also been previously extracted and sequenced in the earlier study.^7^ Additional DNA samples were obtained from 33 donors incorporated into the snMultiome cohort, as well as from 1 donor from the prior study,^7^ whose bulk RNA sample had been independently reprocessed and sequenced. Genomic DNA from these donors was isolated using the QIAamp DNA Micro Kit (Qiagen, cat# 56304). Library construction was carried out with the Watchmaker DNA Library Prep Kit with Fragmentation (Cat. #7K0013-096). Sequencing was performed at the Yale Center for Genomic Research using using the Illumina NovaSeq 6000 system, with each sample sequenced to an average coverage depth of 30x.

### CRISPR activation for SOX5 haploinsufficiency

#### sgRNA design and selection

Single-guide RNAs (sgRNAs) targeting the genomic region of interest were designed using the Broad Institute CRISPick platform (Genetic Perturbation Platform) against the human GRCh38 reference genome. Guides were designed for CRISPR-mediated transcriptional activation (CRIS-PRa) using the SpCas9 NGG PAM system. Predicted on-target activity was assessed using the RS3seq-Chen2013 model, and candidate sgRNAs were evaluated based on predicted activity, position relative to the intended regulatory region, guide ranking, and spatial distribution. Default CRISPick guide-selection parameters were retained unless otherwise indicated. Population-variation filtering using gnomAD was enabled to reduce selection of guides overlapping common genetic variants, whereas target-local mode and selection based on maximum off-target matches were disabled. Candidate sgRNAs were subsequently prioritized according to their predicted on-target activity and suitability for the intended CRISPRa application. A single lead sgRNA exhibiting the strongest overall on-target prediction among the evaluated candidates was selected for experimental validation. For expression from the U6 promoter, complementary DNA oligonucleotides corresponding to sgRNA-1 were synthesized for cloning. An additional guanine nucleotide was incorporated at the 5′ end of the guide where required to support efficient U6 promoter-driven transcription. See also **Key Resources Table**.

#### Lentivirus cloning

Guides sequences were ordered from Integrated DNA Technologies (IDT) as duplexed DNA, and cloned into a LentiSam V2 plasmid (Addgene #92062). Duplexed DNA contained the overhangs necessary to be compatible with LentiSam V2. Both plasmid and guide RNA duplexes were cloned together after golden gate cloning with Esp3I (NEB, R0734) T4 DNA ligase (NEB, M0202), and ATP-containing T4 DNA ligase buffer. Golden gate cycling conditions were chosen following the manufacturer’s suggestions. Cloned plasmids were transformed into NEB Stable competent cells (NEB, C3040); adding no more golden gate reaction than 10% of the total bacterial volume used. We followed the high efficiency transformation protocol provided by the manufacturer for these cells. Colonies were verified through nanopore sequencing (Source BioScience).

#### Lentivirus production, purification and quantification

Lentiviruses were produced by transfecting HEK293T. 293T cells were plated at 5x10^6^ cells per 15 cm plate (CyoOne, Starlab, CC7682-3614) in DMEM GlutaMAX (Gibco, 10566016) supplemented 10% with fetal bovine serum (FBS) (Gibco; A5256701) and 1% antibiotic-antimycotic (Sigma-Aldrich, A5955), 5 plates per guide were used. Cells were transfected the next day. In a tube containing 0.5 mL of Opti-MEM medium (Gibco, 31985070), 6 µg of PSPAX2, 6 µg of VSV-G, and 8 µg of the GOI were mixed and incubated for 5 minutes at room temperature. In another tube with 0.5 mL Opti-MEM, 48 µg of PEI MAX (Kyfora Bio, 24765) were mixed and incubated for 5 minutes at room temperature. After the 5-minute incubation, the DNA is added dropwise into the PEI-containing mixture and left incubating at room temperature for 20 minutes to form polyplexes. Once the polyplexes are formed, they are added dropwise onto their respective dishes.

The next morning the medium is changed to fresh complete DMEM, and left for another 48h before collection. Then, the supernatant is collected and spun for 5’ at 500xg to remove cells. The supernatant is then spun again at 2,000xg for 10 minutes to remove debris. The supernatant is then filtered through a pre-wetted 0.45 µm syringe filter. The filtrate is then concentrated using Lenti-X concentrator (Takara Bio, 631232) following the manufacturer’s protocol. Lentiviral pellets were resuspended in 0.5 mL sterile 1X Phosphate Buffer Saline (Gibco, 10010023), and viral genomes were quantified using a qPCR Lentivirus titer kit (Abcam, ab289841).

#### Stable cell line generation

5x10^5^ Wild-type SH-SY5Y cells (ATCC, CRL-2266) were plated in a 6 well plate (CytoOne, Starlab, CC7682-7506) with 2 mL of complete DMEM. The next day, 60 viral genomes per cell were added per well. After 72h, the medium is refreshed and 10 µg of Blasticidin (Gibco, A1113903) per mL of complete DMEM medium are added as selection marker. Cells were selected in Blasticidin for 10 days while changing the medium and selection marker every 48h.

#### RNA extraction and qPCR

5x10^5^ of each SH-SY5Y stable cell line expressing their respective dCas9-SAM and guide RNA was plated in 6 well plates in triplicates. After 48h, the RNA of each well was isolated using a Promega Maxwell RSC 16 instrument with a Maxwell RSC simplyRNA kit (Promega, AS1340). RNA concentration was measured in a nanodrop instrument. 1 µg of RNA for each sample was used to generate cDNA with a High-capacity cDNA Reverse Transcription kit (Applied Biosystems, 4368814), following the manufacturer’s protocol. The final cDNA was diluted 5 times with nuclease free water. qPCRs were run using SYBR Green 2X Universal Master Mix (Applied Biosystems, 4309155) in an Applied Biosystems StepOne Plus instrument. Each reaction was 20 µl with 2 µl of cDNA per well in MicroAmp optical plates (Applied Biosystems, 4346906). Samples were run with technical duplicates using the Quantitation Comparative CT (ΔΔCT) program for SYBR Green reagents with melt curve at standard speed. Results were analysed using the ΔΔCT and with GAPDH as housekeeping gene. The primer sequences used in this study are provided in the **Key Resources Table**.

#### Protein extraction and western blot

5x10^5^ of each SH-SY5Y stable cell line expressing their respective dCas9-SAM and guide RNA was plated in 6 well plates in triplicates. After 48h, the cells were washed with sterile PBS and 150 µl of 1X Radioimmunoprecipitation assay (RIPA) Lysis and Extraction Buffer (Thermo, 89901) supplemented with 1 tablet of cOmplete Protease Inhibitor Cocktail Tablets (Roche, 04574834001). Then, each lysate was sonicated for 15 s using a Qsonica Q500 instrument at 50% amplitude in pulse mode with 5 s On and 5 s OFF in ice with a 1/16 inch microtip (Qsonica, 417-A). Then, the resulting lysate was spun down for 10 minutes in a microcentrifuge at 10,000xg, and the pellet was discarded. Protein concentration was quantified using a Pierce BCA kit (Thermo, 23227) and measured in a plate reader (BMG Labtech, CLARIOstar). 20 µg of protein were mixed with LDS (Thermo, 84788) and NuPAGE Sample reducing agent (Thermo, NP0004) to a 1X concentration and incubate at 95 °C for 5 minutes and quenched in ice. The samples were loaded into a NuPAGE 4-12%, 1 mm, Bis-Tris Midi gel (Thermo, WG1403) and run in 1X MOPS SDS (Thermo, NP0001) at 120V for 1h and 45 minutes, using 10 µl of the Amersham ECL Rainbow protein ladder (Merck, GERPN800E). The gel was then wet transferred into a 0.45 µm PVDF transfer membrane (Immobilon-FL) (Millipore, IPFL00010) after activation in methanol. The transfer took place in 1X NuPAGE Transfer Buffer (Thermo, NP00061) with 10% methanol. The transfer was run at 70V for 1h 45 minutes. After transfer, total protein was measured by staining the membrane in Fast Green FCF (Merck, F7252). Then, the membrane was blocked in 5% milk in Tris Buffer Saline Tween buffer (TBST) for 1 h at room temperature. Primary antibodies were incubated overnight at 4 °C. After primary antibody incubation, the membranes were washed with TBST 5 times for 5 minutes at room temperature, and incubated at room temperature for 1 h with the secondary antibody in 5% milk TBST. All secondary antibodies are conjugated to HRP. Prior to imaging, all blots were washed 5 times for 10 minutes in TBST buffer. SOX5 was imaged using the ECL substrate SuperSignal West Femto Maximum Sensitivity Substrate (Thermo, 34095), and the β-Actin loading control with Clarity Western ECL Substrate (Bio-Rad, 170-5061). Blots were imaged in a Bio-Rad Chemidoc MP system using the auto exposure mode. Gel densitometry was performed using Fiji. Antibody details, including suppliers and catalogue numbers, are provided in the **Key Resources Table**.

## QUANTIFICATION AND STATISTICAL ANALYSIS

### Data processing of single-nucleus RNA sequencing

Due to inconsistencies in data quality between transcriptomic and chromatin accessibility modalities across several libraries, each modality was processed independently to maximize data retention and ensure robust downstream analyses. Raw sequencing reads were aligned to the human reference genome (GRCh38; Ensembl release 107, Gene Annotation of the ENCODE Project GENCODE v41) using Cell Ranger count with the *--include-introns* option. Each library was evaluated based on the total number of recovered cells, sequencing depth, median genes detected per cell, median Unique Molecular Identifier (UMI) counts per cell, and the distribution of mitochondrial read percentages, as well as overall distributions of reads and genes per cell. Libraries that were under-sequenced relative to the number of recovered cells were subjected to additional sequencing to achieve a median depth of approximately 20,000 reads per cell. In total, 10 of 147 RNA libraries were excluded due to poor sequencing quality or insufficient cell recovery (see **Table S3**). All remaining libraries were retained for library-level quality control, as described below. See also **Key Resources Table**.

### Filtering of low-quality cells and doublets

Quality control was performed at both library and cross-library levels to remove low-quality nuclei and doublets.^73^ Doublets were identified using a combination of computational scoring and marker gene expression. Doublet scores were computed with scDblFinder. To minimize false positives and preserve biologically meaningful clusters, preliminary cell-type identities were inferred using AUCell based on canonical marker genes. AUCell scores were calculated for each cell, and clusters were assigned to a given cell type when the 75th percentile of scores exceeded a cell-type– specific threshold. Clusters exhibiting assignments to multiple cell types and enrichment for high doublet scores (75th percentile exceeding the scDblFinder threshold) were classified as doublets and excluded. Low-quality nuclei were identified based on mitochondrial content and expression of housekeeping genes. Cells with mitochondrial transcript content greater than 5% were removed. Consistent with our previous study,^169^ low-quality nuclei exhibited reduced expression of housekeeping genes that are typically stable in high-quality cortical cells. To define this gene set, six representative libraries were sampled and integrated with previously annotated datasets containing low-quality cell labels. Genes enriched in high-quality cells, as identified by FindMarkers, were designated as housekeeping genes. Clusters showing depletion of the top 50 of these genes were manually annotated as low quality and excluded. All filtering steps were applied both within individual libraries and across libraries to ensure consistent data quality.

### Cell type annotation

To derive consensus cell-type annotations across each developmental period, we first identified major cell classes within each developmental period and subsequently integrated these classes across periods for refinement. Within each developmental period, libraries were integrated using the IntegrateLayers function in Seurat (v5)^98^ with method = “CCAIntegration”. Gene expression matrices were normalized by total counts per cell, scaled to 10,000, and log-transformed. The top 2,000 highly variable genes (HVGs) were identified per library and ranked by variance, from which a shared set of 2,000 HVGs was selected for canonical correlation analysis (CCA)–based integration. PCA was first performed for dimensionality reduction, and CCA integration was applied in PCA space. The top 30 CCA dimensions were used for clustering at a resolution of 0.2. Clusters were annotated to major cell classes based on canonical marker gene expression, including excitatory neuronal lineages (excitatory progenitors, intratelencephalic [IT], and non-IT subtypes), inhibitory neurons (medial ganglionic eminence [MGE] and caudal ganglionic eminence [CGE] lineages), and non-neural cell types (oligodendrocyte precursor cells [OPCs], oligodendrocytes, astrocytes, microglia, and vascular cells). To harmonize and refine annotations across developmental periods, each major cell class was re-integrated using IntegrateLayers with method = “FastMNNIntegration”. The top 30 Mutual Nearest Neighbors (MNN) dimensions were used for clustering at a resolution of 0.2 to define refined subclusters. Final cell-type identities were assigned manually based on canonical marker gene expression. To validate these annotations, labels were transferred from published single-nucleus RNA-seq reference datasets using the TransferData function in Seurat,^98^ with the weight.reduction parameter optimized for each integration space. Prenatal datasets were validated against previously published references,^57,74,170^ while postnatal datasets were compared with independent references.^26,73^ Concordance between transferred labels and manually assigned annotations was used to assess accuracy of the annotation.

### Data processing of single-nucleus ATAC sequencing

To preserve as many nuclei as possible for downstream analyses, the snRNA-seq and snATAC-seq datasets were analyzed independently. Because snATAC-seq data are inherently sparse, matched snRNA-seq profiles from the same nuclei were used to support doublet identification and preliminary cell-type classification.

Raw snATAC-seq sequencing reads were mapped to the human reference genome (hg38; Ensembl release 107, GENCODE v41) using the cellranger-atac count pipeline from 10x Genomics. Fragment files generated from this step were subsequently analyzed with ArchR (v1.0)^99^ for quality assessment, dimensionality reduction, and downstream cell-type annotation.

To reduce the influence of poor-quality libraries on downstream analyses, each library was assessed based on sequencing depth, TSS enrichment, and representation of diverse cell populations. Libraries containing fewer than 100 high-quality nuclei; defined by a TSS enrichment score greater than 4 and more than 1,000 fragments per nucleus, or exhibiting limited cellular heterogeneity were removed from further analysis. For quality evaluation, a genome-wide chromatin accessibility matrix using 500 bp bins was generated and subjected to iterative latent semantic indexing (LSI) for dimensionality reduction. Cell-type diversity within each library was assessed using gene activity scores from established marker genes. Overall, 21 low-quality libraries were excluded from a total of 147 libraries.

Quality-control procedures were carried out at both the individual-library and combined-library levels using an approach consistent with the snRNA-seq workflow. Nuclei with TSS enrichment scores below 4 or fewer than 100 detected fragments were filtered out. Clusters displaying sub-stantially reduced average TSS enrichment or fragment counts were manually inspected and excluded when necessary. Doublet annotations inferred from matched snRNA-seq data were transferred to nuclei shared between modalities. Clusters containing more than 50% predicted doublets were removed prior to downstream analyses.

### Cell type annotation

To achieve concordant cell-type annotations across modalities, labels from both snRNA-seq and snATAC-seq datasets were integrated. For nuclei represented in both datasets, annotations derived from snRNA-seq were prioritized, whereas nuclei detected only in the snATAC-seq dataset were annotated using chromatin accessibility information. Cell-type classification from snATAC-seq data followed the same hierarchical framework applied to the snRNA-seq analysis, in which broad cellular categories were first identified within each developmental stage and then merged across stages for further refinement. Cross-period integration was conducted in ArchR using the addHarmony function applied to the 500 bp genomic tile matrix. Clustering parameters were set to a resolution of 1.0 for excitatory neuronal populations and 0.2 for inhibitory neuronal and non-neural populations. Final annotations were assigned through manual evaluation of gene activity scores for established marker genes. See also **Key Resources Table**.

### Bulk RNA data processing

Sequencing reads were aligned to the human reference genome (GRCh38.p13; GENCODE v41) using STAR (v2.7.10b). To maintain consistency with the GENCODE annotation used for snRNA-seq in our previous work,^7^ the full genome assembly (rather than the primary assembly) and corresponding GENCODE v41 annotations were used. Following the Broad Institute Genotype-Tissue Expression (GTEx) pipeline,^16,171,172^ decoy, alternate, and Human Leukocyte Antigen (HLA) contigs were excluded prior to alignment. Gene-level read counts were generated using htseqcount (v2.0.5) in union mode with strand-specific settings (-s yes) based on GENCODE v41 annotations. In parallel, transcript-level and gene-level expression estimates were obtained using Salmon (v1.10.3), with selective alignment and validated mappings, followed by summarization with tximport (v1.32.0).

Quality control was performed at the raw read, alignment, and gene expression levels. Raw reads were evaluated using FastQC and summarized with MultiQC. Post-alignment metrics, including alignment rate and duplication levels, were assessed using Picard (https://broadinstitute.github.io/picard/). Gene-level QC was conducted on normalized count matrices (CPM and Transcripts Per Million [TPM]), evaluating library complexity, gene detection, and RNA composition metrics, including ribosomal and hemoglobin content, as well as sex concordance. Sample relationships were further examined using PCA, hierarchical clustering, and correlation analyses to identify outliers and batch effects. Samples failing QC criteria were excluded from downstream analyses (see **Table S1)**.

### Whole Genome Sequencing (WGS) Data Processing

WGS data from 271 postmortem brain DNA libraries were processed using NVIDIA Parabricks (v4.0.1-1).^131,173,174^ The fq2bam workflow aligned reads with BWA-MEM to the GRCh38 full analysis set supplemented with the hs38d1 decoy (NCBI assembly accession GCA_000001405.15; Homo_sapiens_assembly38.fasta).^175^ The workflow performed mate fixing and Picard-compatible duplicate marking with an optical duplicate distance of 2,500 pixels (http://broadinstitute.github.io/picard). Base quality score recalibration used dbSNP build 138,^155^ the Mills and 1000 Genomes gold-standard indel set, and the Axiom Exome Plus genotype resource as known variant sites.

### Single-sample variant discovery and joint genotyping

Variant discovery was performed independently for each WGS library using the Parabricks implementation of GATK HaplotypeCaller in genomic variant call format (gVCF) mode.^101,176^ One whole-genome gVCF was generated per library, with variants annotated using the StandardAnnotation and StandardHCAnnotation groups. The same whole-genome gVCF was supplied to each contig-specific joint-genotyping step.

Joint genotyping was performed using a workflow derived from nf-core/sarek (v3.0.1), implemented in Nextflow (v23.10.1).^132^ Per-library whole-genome gVCFs were imported into contig-specific GATK GenomicsDB workspaces using GenomicsDBImport (v4.2.6.1; batch size = 50) with workspace consolidation. GATK GenotypeGVCFs (v4.2.6.1) jointly genotyped the samples using StandardAnnotation and AS_StandardAnnotation annotations. Raw contig-level calls were combined for genome-wide variant quality score recalibration model training, and recalibrated contig-level VCFs were merged in reference contig order.

### Variant quality recalibration

Variant quality score recalibration (VQSR) was performed using GATK VariantRecalibrator^101^ and ApplyVQSR.^101^ Indels were recalibrated using the GATK resource bundle, including the Mills and 1000 Genomes gold-standard indel set^177^ and the Axiom Exome Plus genotype dataset as training resources, with dbSNP build 138^155^ used as a known variant resource. Single nucleotide polymorphism (SNP) recalibration was performed using the GATK resource bundle, including the HapMap 3.3 callset,^160^ the 1000 Genomes Omni 2.5M array, and 1000 Genomes Phase 1 high-confidence SNPs as training resources, with dbSNP build 138 used as a known-sites resource.^155,177,178^ SNP recalibration models incorporated standard GATK annotation metrics, including QualByDepth (QD), MappingQuality (MQ), MappingQualityRankSumTest (MQRankSum), ReadPosRankSumTest (ReadPosRankSum), FisherStrand (FS), StrandOddsRatio (SOR), and depth (DP). Indel models used the same annotations except MQ. Final variant filtering was performed using ApplyVQSR at truth-sensitivity thresholds of 99.0% for indels and 99.8% for SNPs.

### Variant call quality assessment

Variant call quality was evaluated using multiple complementary metrics. Picard CollectVariant-CallingMetrics (v3.1.1) (http://broadinstitute.github.io/picard). was used to summarize variant call characteristics relative to dbSNP build 138. Additional variant statistics were computed using bcftools stats (v1.16),^135^ and transition-to-transversion (Ti/Tv) ratios were calculated using vcftools (v0.1.16).^136^ Quality metrics were assessed both genome-wide and across specific genomic contexts, including segmental duplications, tandem repeats, homopolymer regions, centromeric regions, low-mappability regions, GC-extreme regions, and annotated functional genomic elements. Aggregate quality control metrics were summarized using MultiQC.

### Genetic sex identification

Genetic sex was inferred using both RNA-seq and WGS data. For RNA-seq libraries, X-Inactive Specific Transcript (XIST) expression levels were used to assign XX or XY status, whereas Y chromosome coverage was used for WGS-based sex determination. Among samples passing all other RNA-seq and WGS quality control criteria, both XIST expression and Y chromosome coverage exhibited clear bimodal distributions when visualized as histograms. Genetic sex was therefore assigned by visual inspection of these distributions. Borderline cases were compared with recorded sex, and one discordant case was resolved after confirmation from the originating provider.

### Sample modality matching

Genotype-based matching of bulk RNA-seq libraries with WGS libraries was performed using QTLtools Match BAM to VCF (MBV) v1.3.1.^137,179^ Each RNA-seq binary alignment/map (BAM) file was compared to our previous study joint-called WGS genotype panel,^7^ and diagnostic plots of heterozygous genotype concordance versus homozygous genotype concordance were generated for all bulk RNA–DNA comparisons. Plots were manually inspected to determine match quality; correct donor matches formed clear outliers with high concordance across both metrics, whereas cross-sample contamination was indicated by high heterozygous concordance but reduced homozygous concordance.

When sample swaps were detected, donor identity could not be directly inferred from genotype concordance alone (e.g., if donor HSB515 bulk RNA matched donor HSB516 WGS). Where possible, additional metadata were used to resolve donor identity. Genetic sex was determined as described above, and developmental stage was inferred from RNA-seq data, with fetal and postnatal samples clearly distinguishable by PCA of gene expression. These approaches enabled the identification of matching libraries for 25 donors that had previously been excluded from our previous study.^7^

After pairing libraries generated as part of our previous study,^7^ samples with evidence of contamination or lacking a confident RNA–DNA pairing were flagged for resequencing. When sufficient tissue was available, these samples were re-extracted and resequenced alongside newly generated donor samples, as described above. Specifically, six bulk RNA libraries were regenerated due to cross-contamination concerns, 12 donors had new bulk RNA libraries generated, and 8 donors had new WGS libraries generated. Following resequencing and alignment, QTLtools MBV was applied as described above to match libraries from all three modalities (bulk RNA-seq, snRNA-seq, and snATAC-seq) to the new dataset joint-called WGS genotype panel. For single-nucleus libraries, MBV was applied to full library-level BAM files, which aggregate reads across all barcodes.

No cross-contamination was detected in the new bulk RNA-seq libraries. However, snRNA-seq libraries from one donor showed evidence of cross-sample contamination and were excluded from downstream analyses. In addition, three donors were excluded due to the inability to identify genetically matched RNA and WGS libraries. Libraries were included in downstream analyses only if they (i) passed all quality control criteria, (ii) had a genetically matched WGS library, (iii) showed no evidence of cross-sample contamination, and (iv) had concordance between genetically inferred sex and reported donor sex (see **Table S1**).

### Generic ancestry inference

To infer ancestry (population) labels for study samples, we employed a supervised classification approach using reference populations from the 1000 Genomes Project and the Human Genome Diversity Project (HGDP).^159^ Together, these datasets comprised 4,119 individuals with known population assignments: African (AFR; n = 996), Admixed American (AMR; n = 547), East Asian (EAS; n = 808), Finnish (FIN; n = 92), Middle Eastern (MID; n = 128), Non-Finnish European (NFE; n = 682), South Asian (SAS; n = 788), and Other (OTH; n = 79). The study sample variant dataset was filtered to a common set of 5,847 SNPs mapped to the GRCh38 reference genome, corresponding to the Purcell 5k loci widely used for ancestry inference. Genotypes at these loci were extracted using Hail^102^ and merged with the reference datasets at matching variant positions. PCA was performed on the combined genotype matrix. The top 10 principal components (PCs) derived from the reference individuals were used to capture the major axes of genetic variation across global populations. A random forest classifier (implemented in R) comprising 1,000 decision trees was trained on reference individuals with known population labels to predict ancestry from these 10 PCs. Individuals labeled as “Other” (n = 79) were excluded from model training but retained for visualization in PCA space. The trained classifier was applied to the study samples to infer population membership. For each sample, predicted class probabilities across all population labels were computed, and the label with the highest posterior probability was assigned as the final ancestry designation.

### Structural variant discovery

Prior to structural variant (SV) discovery, we performed stringent sample-level quality control on whole-genome sequencing data, including assessments of sequencing coverage, estimated autosomal and sex chromosome ploidy, and whole-genome dosage (WGD) scores, which quantify deviations in genome-wide sequencing depth from the expected baseline (**Document S1**).

SV discovery was performed using the GATK-SV pipeline,^103^ which integrates multiple complementary algorithms to maximize sensitivity across diverse SV classes. Specifically, Manta^104^ and

Wham^105^ were used for broad SV detection, including deletions, duplications, inversions, and complex rearrangements; Scramble was used for mobile element insertion (MEI) discovery;^180^ and cn.MOPS^107^ together with GATK-gCNV^108^ were applied for CNV detection. Candidate variants identified across samples were integrated and filtered using a random forest classifier incorporating evidence from discordant read pairs, split reads, read-depth changes, and B-allele frequency patterns. High-confidence variants were subsequently jointly genotyped across all samples to improve cohort-wide sensitivity and harmonization. Overlapping variants within individual genomes were further evaluated to identify and resolve potential complex SVs (cxSVs).

Additional refinement procedures were applied to improve callset precision. These included machine learning–based filtering of low-quality genotype calls and assessment of per-sample SV burden to identify outlier samples. One sample (HSB112), which exhibited an excess of singleton deletions within the 5–20 kb size range, was excluded during this refinement stage. The finalized SV callset was functionally annotated using GATK-SVAnnotate to predict gene-disruptive and dosage-altering consequences.

To further assess callset quality, we manually inspected sequencing depth profiles for 15 CNVs larger than 1 Mb and confirmed support for all events. We additionally reviewed a 40 kb deletion affecting the 3′ region of *PACS2*, a constrained Online Mendelian Inheritance in Man (OMIM) gene (loss-of-function observed/expected upper bound fraction [LoEUF] = 0.32), in two samples (**Document S1**). Finally, we manually evaluated 41 predicted loss-of-function (pLoF) CNVs larger than 5 kb affecting moderately constrained genes, observing clear depth support for all but four deletions located within repetitive genomic regions known to be difficult to resolve using short-read sequencing technologies.^181^

The final SV callset comprised 57,745 SV sites across the cohort, with an average of 6,972 SVs identified per genome (**Document S1**). Most autosomal biallelic SVs (83%) satisfied Hardy–Wein-berg equilibrium (HWE) following Bonferroni correction, supporting overall callset quality.

### Common variants filtering and PCA analysis

Common variant filtering was performed using Hail (v0.2)(https://github.com/hail-is/hai)l^102^ on joint-called WGS data. VCF files were imported with GRCh38 as the reference genome and split into bi-allelic records. To reduce false-positive calls arising from problematic genomic regions, variants overlapping annotated low-complexity regions, centromeres, or telomeres were excluded. Variant-level quality control was performed using Hail’s variant_qc module. Variants were required to have a call rate >95% and to pass VQSR filtering. In addition, GATK-recommended hard filters for germline short variants were applied, including QD ≥ 2.0, FS ≤ 60.0, SOR ≤ 3.0, MQ ≥ 40, MQRankSum ≥ –12.5, and ReadPosRankSum ≥ –8.0.

Genotype-level quality control excluded entries with sequencing depth <10X or >200x, as well as those failing expected allele balance thresholds for homozygous reference (≤0.1), heterozygous (0.25–0.75), or homozygous alternate (≥0.9) genotypes. To restrict analyses to common variation, only variants with minor allele frequency (MAF) >2% and HWE *p*-value >1 × 10⁻⁶ were retained. The resulting high-quality matrix table was used for downstream analyses.

For PCA, variants were further restricted to biallelic SNPs and pruned for LD (r² < 0.1 within a 500 kb window) to obtain a set of approximately independent variants. PCA was performed using Hail’s hwe_normalized_pca function with k = 20 components, and per-sample eigenvectors were exported for downstream population structure analyses.

### GWAS summary statistics and variant to gene mapping

We analyzed summary statistics from multiple GWAS data spanning psychiatric disorder, neuro-developmental disorders, neurodegenerative disease, and other traits (see **Table S7** and **Key Resources Table** for full details), including ASD,^3^ SCZ,^5^ ALZ,^1^ ALS,^78^ BIP,^49^ MDD,^14^ attention-deficit/hyperactivity disorder (ADHD),^79^ epilepsy (GEPI and CEPI),^81^ intelligence(IQ),^82^ EA,^83^ height,^84^ Parkinson’s disease,^85^ MS,^25^ anorexia nervosa (AN)/eating disorder,^86^ Tourette’s syn-drome,^87^ PAU,^44^ PTSD,^88^ substance use disorder (SUD),^89^ NDD,^4,77^ and frontotemporal lobar degeneration (FTLD-TDP).^90^

All datasets were harmonized using MungeSumstats (v1.14.11), including removal of indels, lift-over to GRCh37 (hg19), and filtering of variants with imputation INFO scores <0.7 when available. Gene-level association testing was performed using MAGMA (v1.08.2) implemented in FUMA (v1.5.23).^109,110^ SNPs were mapped to genes based on physical proximity, assigning variants located within gene bodies as well as within ±1 kb upstream and downstream of each gene. Only protein-coding genes with at least one mapped SNP were included in the analysis. LD structure was accounted for using population-matched reference panels (e.g., UK Biobank European or 1000 Genomes Phase 3, depending on the GWAS). MAGMA computed gene-level association statistics by aggregating SNP-level effects while accounting for LD between variants. Genes were subsequently ranked by their MAGMA *p*-values to identify trait-associated loci.

Quantile–quantile (Q–Q) plots were generated to evaluate the distribution of observed gene-level association statistics relative to the null expectation for each phenotype. Gene-level *p* values obtained from MAGMA analyses were filtered to exclude missing or zero values prior to analysis. For each phenotype, *p* values were ranked in ascending order and transformed to observed −log10(*p*) values. Expected −log10(*p*) values were calculated using the theoretical quantiles derived from a uniform distribution with the ppoints function in R. Observed and expected distributions were visualized.

### Differentially expressed genes (DEGs)

To characterize transcriptional signatures across brain cell types and developmental stages, we performed a series of differential gene expression analyses on snRNA-seq data using FindMarkers by Seurat package in R. Cells annotated as “Unknown” in the fine-grained cell type classification were excluded prior to all analyses. A uniform quality control filter was applied across all datasets, retaining only cells with ≤15% mitochondrial gene content and ≥0.25% ribosomal gene content to remove low-quality, stressed, or poorly captured cells. Developmental stage was defined from the period metadata field, with periods 3–6 classified as prenatal, period 7 as transitional, and periods 8 and above as postnatal.

#### Cell-type marker identification

To define transcriptional markers of each cell population, we performed one-versus-all differential expression analysis independently within the prenatal and postnatal datasets using FindAllMarkers. Cell type annotations were examined at three levels of resolution: (i) broad categories (ExN, InN, glia, NNC, and progenitors; (ii) fine-grained subtypes from celltype_v1. For all comparisons, a minimum of 10 cells per group and 10 cells expressing each feature were required (min.cells.group = 10, min.cells.feature = 10). The Wilcoxon rank-sum test was used as the default statistical framework throughout.

#### Neuron versus non-neuronal cell comparisons

To characterize transcriptional differences between neuronal and non-neuronal brain cell populations, we constructed a binary classification variable distinguishing neurons from specific non-neuronal types including astrocytes, microglia, endothelial cells, oligodendrocyte precursor cells, vascular leptomeningeal cells (VLMCs), smooth muscle cells (SMCs), and oligodendrocytes. FindAllMarkers was then applied to this grouping independently in the prenatal and postnatal datasets, comparing each non-neuronal cell type and the pooled neuronal population against all others, using the same minimum cell and feature thresholds described above.

#### Prenatal versus postnatal DEG analysis, cell-type stratified

To identify genes with stage-predominant expression within each cell population, we performed directed pairwise comparisons between prenatal and postnatal cells using FindMarkers. This analysis was conducted in two passes: first across four broad cell type categories (ExN, InN, NNC, Glia) and second across all fine-grained cell subtypes defined by celltype_v1. For each cell type, the object was subset to cells of that identity prior to comparison, and only cell types represented in both developmental groups were tested. The same minimum cell and feature thresholds were applied throughout. Cell types lacking sufficient group representation were skipped.

#### Combined cell type and developmental stage analysis

To simultaneously capture cell-type-predominant and stage-predominant transcriptional variation, we constructed a combined identity variable by concatenating each cell’s unified cell type label and developmental stage designation (e.g., excitatory neurons, ExN_prenatal; inhibitory neurons, InN_postnatal). This combined labelling was applied to excitatory and inhibitory neurons, microglia, and oligodendrocyte precursor cell populations from the merged prenatal–postnatal Seurat object. FindAllMarkers function was used for the analysis.

An adjusted *p*-value threshold of < 0.05 and an average log₂ fold change (log₂FC) > 0.5 were applied in all analyses to identify marker genes. Progenitor cells were excluded from the main focus of the study due to the limited number of cells available; however, the results were included in supplementary materials. Top 10 genes per cell type and developmental stage based on adjusted significance and fold-change were visualized using heatmap plot. Protein-coding genes were retained through annotation with the org.Hs.eg.db database, and the pseudobulk expression profiles were generated using AggregateExpression, followed by row-wise min–max scaling. Heatmaps were constructed with the ComplexHeatmap package using scaled expression values across prenatal and postnatal cell populations, with annotations indicating developmental stage and cell type.

### Differentially accessible regions (DARs)

To identify DARs across cell types and between developmental stages, we performed differential accessibility analysis on snATAC-seq data using the ArchR R package (v1.0).^99^ Cell type annotations transferred from the paired RNA modality were used throughout to ensure consistency between modalities. A broad cell type classification (broad.type) was derived by collapsing rna_majorclass_v1 labels into five categories, ExN, InN, glia, NNC, and progenitors, using the same mapping applied in the RNA analyses. Cells annotated as “Unknown” were excluded prior to all differential analyses.

#### Cell-type marker accessibility

To identify chromatin regions that distinguish cell types within each developmental period, we applied getMarkerFeatures to the PeakMatrix of each stage-pre-dominant ArchR project separately, grouping cells by fine-grained (rna_celltype_v1and broad (cell_broad) cell type annotations in turn. Cell types with fewer than 50 cells were excluded to ensure statistical stability; up to 10,000 cells per group were subsampled for computational efficiency. The Wilcoxon rank-sum test was used throughout, with bias correction applied for TSS enrichment score and log10-transformed fragment count to account for technical variation in ATAC-seq data quality.

#### Neuron versus non-neuronal cell accessibility

To characterise broad chromatin differences between neuronal and non-neuronal populations, inhibitory and excitatory neurons were grouped as “Neuron.” The getMarkerFeatures function was then applied to this grouping, comparing each non-neuronal type against all others including the Neuron class, using the same Wilcoxon test and bias correction parameters described above.

#### Prenatal versus postnatal DARs analysis, cell-type stratified

To identify developmentally regulated chromatin regions within each cell type, prenatal and postnatal ArchR projects were merged into a single combined project. Developmental stage was assigned period labels: periods 3–6 were designated prenatal, period 7 (transition) was excluded, and periods 8 and above were designated postnatal. The analysis restricted to excitatory neurons (ExN), inhibitory neurons (InN), Microglia, and oligodendrocyte precursor cells (OPC) due to having those cell types both prenatal and postnatal. Group-level pseudobulk coverages were computed with addGroupCoverages, peaks were called jointly across all celltype_dev groups using MACS2 via addReproduciblePeakSet, and a PeakMatrix was constructed with addPeakMatrix. Differential accessibility between prenatal and postnatal within each cell type was then assessed using getMarkerFeatures applied, with the same Wilcoxon test, bias correction, and subsampling parameters as above. The analysis was run in parallel at both the broad cell type level (ExN, InN, Glia, NNC) and the fine cell subtype.

#### Combined cell type and developmental stage analysis

As a complementary analysis, differential accessibility across both cell types and developmental stages was assessed. The combined ArchR project was filtered to excitatory and inhibitory neurons, Microglia, and oligodendrocyte precursor cell populations, and a unified cell type annotation was constructed. A combined grouping variable (celltype_dev) was created by concatenating unified cell type and developmental stage (e.g., excitory neuron, ExN_prenatal; inhibitory_neuron, InN_postnatal), and getMarker-Features was applied with the same Wilcoxon test, bias correction parameters, and subsampling as above.

A False Discovery Rate (FDR) threshold of < 0.05 and an average log₂ fold change (log₂FC) > 0.5 were applied in all analyses to identify marker genes.

### Gene activity score

Gene activity scores for broad cell types (ExN, InN, Glia, NNC, and Progenitor) were computed from the snATAC-seq data using ArchR’s addGeneScoreMatrix^99^ function with an exponential distance-decay model, exp(−∣ *x* ∣/10000) + exp(−1), which weights accessibility at regulatory elements according to their distance from gene bodies. The resulting gene activity scores were then used to identify differentially activated genes from the ATAC data using the same analysis pipeline described in the previous section, with the gene activity score matrix serving as the input. This approach helped the assignment of regulatory regions to their corresponding genes for trait gene enrichment analysis using ATAC data.

### Trait gene enrichment analysis

To assess the enrichment of brain disorder-associated genes within cell-type-predominant and/or development-predominant markers, we performed GSEA using the fgsea R package (v1)^111^ with the fgseaMultilevel function. Disorder gene sets were defined for 24 traits spanning neurodevelopmental, psychiatric, neurodegenerative, and other traits as control. Rare variant gene sets for ASD^2^ and NDD^4^ and common variant gene sets derived from MAGMA gene-level association analysis of published GWAS summary statistics (see **Table S7**; **Data S5;** and **Key Resources Table**). For common variant traits, the top 500 genes reaching MAGMA significance (*p* < 0.05) were retained per trait. Rare variant gene sets^2^ were filtered to genes meeting study-specific significance thresholds (FDR-TADA ≤ 0.05 for ASD; significant = TRUE for NDD).

Enrichment analysis was performed for both RNA differential expression and ATAC-seq gene activity marker genes across three gene sets: cell-type-predominant, developmental stage-pre-dominant, and cell-type-and-stage DEGs. Genes were ranked by sign(log2FC) × (−log10(adjusted *P*)). Genes detected in fewer than 20% of cells in both compared groups were excluded prior to ranking. GSEA was run using fgseaMultilevel with gene set sizes between 5 and 500 and eps = 0 for precise *p* value estimation. Normalized enrichment scores (NES), nominal *p* values, and FDR-adjusted *p* values were computed for each trait-by-cell-type combination and visualized as heatmaps using ComplexHeatmap,^112^ with disorder traits ordered by age of onset along rows and cell types or developmental stages along columns in **Figure 2A,D**.

### Heritability estimated from SNPs in open chromatin regions

The open chromatin regions, calculated before, were amalgamated into prenatal and postnatal sets for our analyses. We matched SNPs found in our samples to the set of SNPs reported by each GWAS described earlier (see **Table S7**), limiting analyses to the set of SNPs shared by each GWAS and was also characterized in our samples. However, we removed SNPs with MAF < 0.05 in our sample; if they fell within the Major Histocompatibility Complex (MHC) region (chr6: 28,510,120-33,480,577, GRCh38); or if their association chi-square test statistic from the GWAS was large and therefore too influential (max(80,0.001×max(N)). These steps yielded a set of SNPs for heritability analysis for each GWAS, ranging from 1,043,916 SNPs for the GWAS of height to 5,017,797 for the GWAS of Parkinson’s disease (see **Table S11**). LD between alleles of SNPs, measured as r, was estimated from our full sample. Next, following the default settings and recommendations of the previous study,^30^ we estimated cell-type specific heritability from SNPs falling in the set of open chromatin regions of cell types by stratified LD score regression (LDSC).^30^ We did, however, set a lower bound for total heritability z-score to 5.1, versus the recommended 7, to allow for the analysis of CEPI; z-scores for all other traits exceeded 7. Also, although the sLDSC program only outputs z-scores, for illustration we translated z-scores to *p* values. sLDSC uses a method-of-moments solution that allows negative estimates of heritability, which should be noted because heritability must be ≥ 0.

### Identification of shared regulatory domains using non-negative matrix factorization

Leading-edge genes from significant (adjusted *p* < 0.05) FGSEA enrichments (both RNA and ATAC) were used to construct a binary trait-by-gene matrix, indicating the presence or absence of each gene (**Table S13**). Only major cell-type enrichments were included, and traits with fewer than 10 leading-edge genes were excluded. The final matrix included 22 traits and 1,980 genes.

NMF decomposed the input trait-by-gene matrix into trait-by-domain (*W*) and domain-by-gene (*H*) loading matrices. This approach allows multiple traits and genes to contribute to multiple latent domains. Five domains (*k* = 5) were used based on biological interpretability and reconstruction error. To assess the stability of the decomposition, 100 random initializations were performed (maximum 1,000 iterations per run), and the solution with the lowest reconstruction error was retained. The best reconstruction error was 43.38, compared with a mean of 43.54 across restarts. Domain stability was evaluated by matching components across restarts using the Hungarian algorithm based on cosine similarity. The mean matched cosine similarity ranged from 0.90 to 0.98 across the five domains. Trait-level stability was assessed as the frequency of dominant-domain assignment across restarts. Most traits showed stable assignments, although TS (0.53), panic disorder (0.58), NDD GWAS (0.66), and ADHD (0.66) showed lower stability. Domain contributions were normalized within each trait to facilitate comparison across disorders. GO Biological Process enrichment was tested using the top 25, 50, and 100 genes ranked by their NMF loadings, with the full set of 1,980 input genes used as the background. The top 100 genes from each domain were retained for visualization and further characterization.

To assess the influence of MHC genes, NMF was performed both with and without MHC-associated genes as a sensitivity analysis. After MHC exclusion, the input matrix contained 1,919 genes, and the mean matched cosine similarity ranged from 0.53 to 0.98. A large proportion of the immune domain involved MHC genes, and this domain was sensitive to MHC exclusion. This is consistent with the contribution of this region to multiple sclerosis and Alzheimer’s disease.

### Expression and chromatin accessibility trajectory

Cell type-predominant gene expression trajectories across brain development were visualized using pseudobulk CPM values. For each gene, CPM values were log₂-transformed following addition of a pseudocount of 1 (log₂[CPM + 1]) to stabilize variance and accommodate zero values. Developmental stage was expressed as post-conceptional days (PCD) on a logarithmic. Smoothed expression trajectories were estimated for each cell type independently using locally estimated scatterplot smoothing (LOESS) regression, as implemented in the Python loess package,^147^ with 95% confidence intervals computed from the standard errors of the local fits. All figures were generated in Python using matplotlib. Explore the data here: www.brainvar.org.

Genome-wide ATAC-seq coverage tracks were generated using the getGroupBW() function implemented in the ArchR package. Pseudobulk BigWig tracks were computed for each cell type and development group from the ArchR project using a tile size of 100bp and a maximum of 10,000 cells per group.^99^ Signal intensities were normalized using the ReadsInTSSmethod to account for sequencing depth and TSS enrichment differences across samples. Generated BigWig files were visualized in the UCSC Genome Browser for inspection of chromatin accessibility profiles across genomic loci (https://genome-test.gi.ucsc.edu/cgi-bin/hgTrackUi?db=hg38&g=single-CellSignalsPeaks).

### Pathway and STRING enrichment

For all pathway enrichments, the enrichR R package^113^ was used against the Reactome_2022 database.^164^ Gene sets were defined for groups representing leading-edge genes from the GSEA analyses, including enriched NDD rare-associated inhibitory neuron genes (prenatal), ASD- and NDD-associated excitatory neuron genes (prenatal), and MS/ALZ-associated glia genes (postnatal). For each gene set, enrichment was assessed against all Reactome pathways and terms were ranked by a combined score computed as −log10(adjusted p value) × log(gene count + 1). The top 3 terms by combined score with Benjamini-Hochberg adjusted p value < 0.05 were retained for visualization. Later, for NDD_rare genes enriched in prenatal inhibitory neuron markers, we performed enrichment analysis using the STRING database through the STRING web platform^165^ and identified protein–protein interaction complexes involving these genes. The resulting network was visualized using Cytoscape software (see **Table S12, Figure 2E**, and **Figure S2D**).

### Motif discovery

Motif discovery analysis was performed using HOMER^72^ on the DARs and caQTL genomic regions to identify transcription factor binding motifs enriched relative to background sequences. Known transcription factor motifs were evaluated for enrichment by comparing their occurrence in the target and background sequences. Motifs were considered significantly enriched when the Benjamini-Hochberg adjusted *p* value was <0.05. The distributions of motif significance across analysis runs were visualized using violin plots, with individual motifs represented as points and the median value indicated for each run.

### Gene Ontology–based functional categorization of genes in brain disorders

Gene Ontology (GO)-based functional categorization^163^ was performed for ASD,^2^ NDD,^76,77^ MDD,^14^ and SCZ genes^5^ (see **Table S9; S10**). Genes were classified into three biologically interpretable categories based on cellular compartment GO annotations. “Neuronal Communication” genes were defined as genes annotated with GO:0005886 (plasma membrane) and/or GO:0045202 (synapse), but lacking GO:0005634 (nucleus) annotation. “Transcriptional Regulation” genes were defined as genes annotated with GO:0005634 (nucleus) but lacking plasma membrane and synapse annotations. Genes not meeting either criterion were classified as “None/Other.” GO annotations were retrieved from the org.Hs.eg.db database^148^ using AnnotationDbi.^149^ Developmental and cell type–specific expression differences between functional groups were visualized using violin plots of normalized expression values [log2(CPM + 1)] across prenatal and postnatal developmental stages.

### BMIND cell type deconvolution

Cell-type proportions in bulk RNA-seq samples were estimated using *Bisque*,^115^ with matched snRNA-seq data serving as the reference. To improve analysis power, closely related cell types were aggregated into broader categories: ExN, InN, Glia, NNC, and progenitors. To maintain consistency with developmental differences observed in single-nucleus data, deconvolution was performed separately for prenatal and postnatal samples. Marker genes were initially identified using FindMarkers (Seurat v5.0.3) and subsequently refined using MarkerPen^182^ to enhance cell-type specificity (**Table S14**). Cell type–predominant gene expression profiles for each bulk sample were estimated using bMIND,^33^ incorporating bulk RNA-seq read counts and inferred cell type proportions. Models were adjusted for relevant covariates, including sex, RIN, developmental period, batch, and ancestry components. As in the deconvolution step, analyses were performed separately for prenatal and postnatal samples. The resulting estimates were treated as pseudo-counts and used for downstream cell type–specific eQTL analyses.

### Expression QTL

cis-eQTL mapping was performed using TensorQTL v1.0.10^116^ with matched WGS genotype data and bulk RNA-seq, pseudobulk RNA-seq, and bMIND-derived expression matrices for broad cell types during developmental stages (prenatal and postnatal). Genotype data were obtained from joint-called and quality-controlled WGS variant call files. Samples lacking developmental annotations, duplicate samples, and samples failing genotype or RNA-seq quality control were excluded prior to analysis (see **Table S1**). Bulk RNA-seq gene expression matrices were generated from HTSeq^117^ read counts and normalized using trimmed mean of M values (TMM) normalization implemented in edgeR.^118^ Genes were retained if they were expressed in at least 23 samples and had an average CPM > 1. For pseudobulk snRNA-seq analyses, donor-level pseudobulk expression matrices were generated separately for each broad cell type. TMM normalization was applied independently for each cell type. Individuals contributing fewer than 10 cells to a given cell type were excluded. Genes were retained if they were expressed in at least 10 samples with average CPM > 1, and genes with zero variance were removed separately within prenatal and postnatal groups prior to eQTL mapping. Cell-type-predominant genes were retained if they passed these filtering criteria. After preprocessing, 223 donors included in the analysis.

For bMIND-derived expression analyses, prenatal and postnatal samples were processed separately. Cell-type-predominant expression estimates were TMM normalized, and genes were retained if expressed in at least 13 prenatal samples or at least 10 postnatal samples with average CPM > 1. Covariates included sex, developmental age or developmental period, RNA integrity number (RIN), PMI, tissue pH (when available), sequencing batch, XIST RNA depth, genotype PCs, and expression PCs derived from PCA of normalized expression matrices. Genotype PCs were computed using variants filtered at MAF ≥ 0.02. cis-eQTL mapping was performed using TensorQTL testing variants located within ±1 Mb of each gene TSS. For bulk RNA-seq and bMIND analyses, variants with MAF ≥ 0.05 were included. For pseudobulk analyses, variants with MAF ≥ 0.1 were included. The same thresholds were applied for interaction analyses (maf_threshold_interaction). Nominal association testing was performed using TensorQTL, and variant-gene pairs with nominal *p* < 1 × 10^5^ were retained for downstream analyses. The eQTL results can be explored in the designed interactive website: www.brainvar.org.

The π1^183^ method was used to quantify replication of eQTL signals between cell types and developmental stages. For each pairwise comparison, one dataset was designated as the test dataset and the other as the discovery dataset. Significant gene–SNP associations were first identified in the test dataset using a nominal *p-*value threshold of 1 × 10^5^. The same gene–SNP pairs were then retrieved from the discovery dataset, and their *p-*value distribution was used to evaluate evidence for replication. For eGene analyses, only the lead SNP per gene in the test dataset was retained. Replication was quantified by estimating the proportion of true non-null associations (*π*_1_) from the discovery *p-*values corresponding to significant test associations. The proportion of true null hypotheses (*π*_0_) was estimated using Storey’s spline-based approach across *λ* values ranging from 0.05 to 0.95, and *π*_1_ was calculated as 1 − *π*_0_. Higher π1 values indicate stronger replication and greater sharing of genetic regulatory effects between datasets.

Similar to previous study,^7^ scatter plot was used to visualize eQTL results categorized in five groups: prenatal predominant, prenatal trending, constant, postnatal predominant, and postnatal trending. For each developmental stage, the lead variant per gene was defined as the SNP with the minimum *p* value across all tested cis variants. Lead associations from prenatal and postnatal datasets were merged, and for each gene the most significant association across both developmental periods was retained. Genes were included for downstream analyses if the lead association reached a nominal significance threshold of *p* < 1 × 10⁻⁵ in at least one developmental stage.

To compare developmental-stage-predominant effects, the corresponding SNP effect sizes (slope), standard errors, and nominal *p*-values were extracted from both prenatal and postnatal datasets for each retained lead variant. Missing associations in one developmental stage were assigned *p* = 1 and beta = 0. Differential effect sizes between prenatal and postnatal stages were assessed using a two-sided Z-test based on the difference in beta coefficients normalized by the combined standard error. Two-sided *p*-values were calculated from the standard normal distribution. Associations were subsequently classified into five developmental categories adapted from the previous work.^7^ Prenatal-predominant eQTLs were defined as associations significant in the prenatal stage (*p* < 1 × 10⁻⁵) but absent postnatally (*p* ≥ 1 × 10⁻⁵), with significantly different effect sizes between stages based on the Z-test (*p* < 0.05). Postnatal-predominant eQTLs were defined analogously, with significance restricted to the postnatal stage. Constant eQTLs were defined as associations significant in at least one developmental stage, nominally significant in both stages (*p* < 0.05), and showing concordant effect directions. Remaining associations were categorized as prenatal-trending or postnatal-trending according to whether the absolute prenatal or postnatal effect size was larger, respectively. For visualization purposes, effect sizes were normalized using a robust Z-score transformation based on the median and interquartile range (IQR). Robust scaling was performed separately for prenatal and postnatal effect sizes within each cell type to reduce the influence of outlier associations and facilitate cross-condition comparison.

### Chromatin accessibility QTL

Reproducible peaks were identified for each cell type and developmental group using MACS2^119^ through the addReproduciblePeakSet function in ArchR. Pseudobulk ATAC-seq profiles were generated by aggregating read counts at the donor level within each cell type. Peaks overlapping ENCODE blacklist regions and samples failing predefined quality control criteria were excluded (see **Table S1**). Donor-level count matrices were normalized using library size scaling, followed by TMM normalization and transformation to log-transformed counts per million (logCPM). For caQTL analyses, logCPM values were further quantile normalized across donors. PCA was performed within each cell type to identify and exclude outlier donors prior to downstream analyses. After preprocessing, 90 donors included in the analysis. For each prenatal and postnatal cell type, quantile-normalized donor-level profiles were combined with peak genomic coordinates to generate TensorQTL-compatible donor-by-peak phenotype matrices. Peaks with zero counts in more than 20% of donors were excluded. Covariates included sequencing batch, sex, developmental period, PMI, the top five ATAC-seq PCs, and the top three genotype principal components. Genotype PCs were computed using variants MAF ≥ 0.02. After preprocessing and removing outliers, the number of donors retained for each cell type was as follows: **prenatal** (n = 26): ExN (n = 24), glia (n = 25), InN (n = 26), NNC (n = 20), and progenitors (n = 26); **postnatal** (n = 64): ExN (n = 63), glia (n = 64), InN (n = 61), and NNC (n = 62). cis-caQTL mapping was performed using TensorQTL^116^ testing variants located within ±1 Mb of each peak. Nominal association testing was performed using TensorQTL, and variant-peak pairs with nominal *p*-values <1e-5 were retained for downstream analyses.

### Splicing QTL

Paired-end RNA-seq reads were aligned to the GRCh38 reference genome using GENCODE v41 annotations with STAR in a two-pass mapping strategy. In the first pass, splice junctions detected across all samples were aggregated and filtered to remove low-confidence junctions, including mitochondrial junctions, non-canonical splice motifs, junctions supported only by multi-mapping reads, and those with insufficient read support (<2). The resulting high-confidence set of junctions was incorporated into the splice-junction database for the second pass to improve alignment accuracy and sensitivity for unannotated introns. During the second pass, allele-specific mapping bias was reduced using STAR’s implementation of WASP.^120^ Reads flagged by WASP as potentially biased were removed using samtools, retaining only reads that passed the WASP remapping consistency check. The resulting WASP-filtered, coordinate-sorted BAM files were used for downstream intron usage quantification.

### Intron quantification and normalization

Intron excision was quantified using LeafCutter.^121^ Splice junctions extracted from the WASP-filtered BAM files were summarized into intron-level counts and grouped into clusters defined by shared splice donor or acceptor sites. Clustering was performed using the LeafCutter pipeline with the following filtering parameters: a minimum of 50 reads per cluster, a maximum intron length of 500 kb, a requirement that each intron contribute at least 0.1% of total cluster reads, and at least 5 supporting reads per intron. In total, 383,851 introns from 80,199 clusters, mapping to 20,364 genes, were identified.

Intron usage was calculated as the proportion of reads supporting each intron relative to the total cluster counts. Additional quality-control filtering removed non-autosomal introns, introns observed in too few individuals, and introns with minimal variability across samples. Usage values were then standardized per intron and transformed using rank-based inverse normal (quantile) normalization. After filtering and normalization, 285,600 introns from 55,268 clusters, corresponding to 14,542 genes, were retained for downstream sQTL analysis. To account for broad technical and biological variation in intron usage, PCA was performed separately for prenatal and postnatal samples on the normalized intron usage matrices.

### Intron annotation

To classify introns detected by LeafCutter and to distinguish annotated splice junctions from cryptic splicing events, we generated a comprehensive intron annotation reference from the GEN-CODE v41 gene transfer format (GTF). The reference annotation was converted into intron- and splice-site–level databases containing all known introns, 5′ splice sites, 3′ splice sites, and exon coordinates. Detected LeafCutter introns were then matched against these reference features by comparing both splice sites of each intron to annotated donor and acceptor positions.

For each intron, the corresponding gene was assigned by identifying the gene most consistently represented across all matching splice-site annotations within the intron’s cluster. Classification of introns was based on the concordance between the detected donor and acceptor sites and the known splice sites from the reference. Introns whose cluster lacked reliable strand information were designated as unknown strand. Introns with neither splice site matching any annotated gene were classified as cryptic unanchored, whereas introns with only one annotated splice site were categorized according to whether the annotated site occurred at the 5′ or 3′ end of the intron, taking into account the gene’s strand orientation. Introns with a known splice site on only one end but unclear strand assignment for the cluster were labeled as cryptic. Introns for which both splice sites matched a known annotated intron were considered annotated, while introns whose donor and acceptor sites were individually annotated but had never been observed together as a pair were classified as novel annotated pairs. After preprocessing, 219 donors included in the analysis. Finally, TensorQTL pipeline^116^ was used to identify sQTLs for prenatal and postnatal stages separately. Genotype PCs were computed using variants filtered at MAF ≥ 0.02. cis-caQTL mapping was performed using TensorQTL testing variants located within ±1 Mb of TSS region. Nominal association testing was performed using TensorQTL, and variant-splicing pairs with nominal *p*-values < 1e-5 were retained for downstream analyses.

We visualized sQTL events using LeafCutter-derived intron excision and junction count data from RNA-sequencing analyses. Sashimi plots were generated in R using the make_differential_splicing_plot() function from the LeafCutter/leafviz framework.^121^

### QTL overlapping with previous studies

Overlap analyses between our QTLs and external datasets^9,16,17^ were performed to validate our results. Significant (*p* <5 × 10^-8) prenatal and postnatal eQTLs were matched to GTEx v10 prefrontal cortex eQTLs,^16,34^ prenatal cortex eQTLs,^9^ and Siebert postnatal dIPFC eQTLs^17^ (see **Key Resources Table**). For overlapping eQTLs, regression slope coefficients from each dataset were independently standardized by z-score normalization prior to correlation analysis. Pearson and Spearman correlations, allelic concordance rates, and ordinary least squares regression slopes were calculated using the standardized effect sizes. Scatter plots were visualized using ggplot2 and ggpointdensity,^122^ with axis limits scaled according to the 95th percentile of the absolute z-score distributions. For large overlap sets, up to 20,000 points were randomly sub-sampled for visualization, while all statistical analyses were performed using the complete dataset.

For sQTL analyses, our sQTLs and GTEx v10 sQTLs for prefrontal cortex^34^ were harmonized using matched Reference SNP Cluster Identifiers (rsIDs) and splice-cluster coordinates, and overlaps between prenatal and postnatal datasets were visualized using Venn diagrams and scatter plot. Splicing start site distance distributions of overlapping sQTLs were assessed using kernel density estimation plots. For caQTL analyses, overlapping caQTLs and bulk brain and T cell caQ-TLs^35^ were identified using shared variant–peak pairs, and effect-size concordance was evaluated using z-score–standardized regression slopes and correlation analyses as described for eQTLs.

### Colocalization and fine mapping

To assess whether GWAS signals and cis-eQTL signals at overlapping loci were driven by shared causal variants, we performed Bayesian colocalization analysis using the coloc R package (v5.2.3).^123^ GWAS summary statistics were harmonized to a common allele coding, and cis-eQTL summary statistics were restricted to genes within ±1 Mb of each genome-wide significant GWAS lead variant (*p* < 5 × 10⁻⁸). For every gene–locus pair, GWAS and QTL datasets were intersected on chromosome, position, and allele identity, and minor allele frequencies were folded to the minor allele in both datasets. Colocalization analyses were performed separately using bulk prenatal and postnatal eQTL datasets, as well as BMIND-derived cell type–specific eQTL datasets across ExN, InN, Glia, and NNC contexts. Tests were conducted with study type = “quant” for both traits and retaining default prior probabilities. Posterior probabilities for the five competing hypotheses (PP.H0–PP.H4) were extracted for each test. Signals with PP.H4 ≥ 0.8 were considered evidence of colocalization, consistent with thresholds adopted in prior brain QTL–GWAS integration stud-ies,^16,184^ while signals with 0.5 ≤ PP.H4 < 0.8 were retained as suggestive and evaluated separately.

To relax the single-causal-variant assumption underlying coloc.abf at loci exhibiting allelic heterogeneity, we additionally applied the SuSiE extension of coloc^124^ using in-sample LD matrices calculated from our WGS data. For each GWAS locus, the LD matrix was computed using plink2,^125^ specifically with options r-unphased, ref-based, and square, to obtain correlations between SNPs that were based on the reference allele count and relative frequency. For each gene– locus pair, fine-mapping was first performed independently on the GWAS and QTL signals using runsusie, followed by pairwise colocalization testing across all credible sets identified in the two analyses. Signals with PP.H4 ≥ 0.8 were classified as colocalized, and the credible set containing the variant with the highest SNP-level posterior inclusion probability was used to nominate the candidate shared causal variant.

We further investigated candidate causal variants within the *AHI1, FADS1,* and *GPX1* loci that showed evidence of colocalization between GWAS and eQTL signals. Variants in the locus demonstrating significant associations in both GWAS and eQTL were annotated using multiple functional genomic features, including chromatin accessibility, chromatin interaction evidence from High-Throughput Chromosome Conformation Capture (Hi-C) datasets, and predicted variant consequences obtained from Ensembl Variant Effect Predictor (VEP).^150^ Additional annotations were obtained through the Functional Annotation of Variants Online Resource (FAVOR) database^161^ (see **Key Resources Table**). Candidate variants were subsequently prioritized through manual evaluation of the combined genetic and functional evidence across annotations.

Regional GWAS and cis-eQTL association plots were generated in R using the locuszoomr package.^140^ GWAS summary statistics and cis-eQTL data were used to construct locus-specific association plots centered on predefined index variants (potential casual variant) with a ±1-Mb genomic window. eQTL analyses were restricted to variants associated with the target gene of interest. The results can be explored in the dedicated BrainVar website: www.brainvar.org.

### Cell–cell communication inference

Cell–cell communication was inferred from snRNA data using CellChat v2 2.2.0.9001.^43,185^ ***Cell-Chat*** integrates gene expression profiles with a database (CellChatDB.human) of ligand–receptor interactions to predict intercellular signaling networks. CellChat analysis was performed independently for each sample, rather than pooling cells across individuals, to preserve biological replication at the donor level for subsequent genotype-associated statistical analyses. Expression data were obtained from the lognorm layer of the merged snRNA H5AD object. Only cells belonging to donors assigned to either the *AHI1* rs13197384 risk or healthy comparison groups were retained. Duplicate mapped gene symbols were made unique before construction of CellChat expression matrices. To reduce computational overhead, donor-level expression matrices were subsequently restricted to genes represented as ligands or receptors in CellChatDB.human. Cell-Chat objects were generated for excitatory and inhibitory neurons, microglia, and oligodendrocyte precursor cells. Cell populations represented by fewer than 10 cells within an individual donor were excluded. Donors containing fewer than two eligible cell populations after this filtering step were not analyzed. CellChat objects were generated using createCellChat() with sparse-matrix representation and cell identity specified by the corresponding communication group. CellChat’s human interaction database was assigned to each object, after which signaling genes and interactions were identified using subsetData(), identifyOverExpressedGenes(), and identifyOverExpressedInteractions(). Communication probabilities were estimated with computeCommunProb(). The primary analysis used a 10% truncated mean (type = “truncatedMean”; trim = 0.10) to estimate average expression of signaling genes within each cell population. Cell-population-size weighting was disabled (population.size = FALSE), such that inferred communication strengths were not directly scaled according to the relative abundance of each cell population.

Communication probabilities and associated CellChat permutation *p* values were estimated using 100 resampling iterations (nboot = 100). After probability estimation, interactions involving populations represented by fewer than 10 cells were additionally removed using filterCommunication(min.cells = 10). Signaling-pathway-level probabilities were then calculated using computeCommunProbPathway(), and overall communication networks were summarized using aggregateNet(). To assess the robustness of the inferred communication networks to the method used to estimate cell-group expression, the complete donor-level CellChat analysis was repeated using CellChat’s triMean estimator (type = “triMean”). All other parameters were kept unchanged. Results obtained using the 10% truncated mean were considered the primary communication analysis, whereas triMean results were treated as a sensitivity analysis.

Differential communication analysis focused on prenatal samples. Only donors that both belonged to the predefined *AHI1* genotype groups and had a successfully generated CellChat object were included in downstream analysis. *AHI1* rs13197384 genotype status was modeled as a two-level variable, with healthy donors used as the reference category and risk samples (homozygous) as the comparison group. For each biological donor, CellChat ligand–receptor communication probabilities were extracted from the net$prob. Each observation was defined by the identity of the sender cell population, receiver cell population, ligand–receptor pair, and donor. Directed communication features were represented as: sender cell population | receiver cell population | ligand–receptor pair. Before statistical testing, detection frequencies were calculated from the unfiltered CellChat outputs. Ligand–receptor pairs with fewer than four nonzero sender–receiver– sample observations were excluded from further analysis. Individual sender–receiver–ligand–receptor settings were subsequently required to have a communication probability greater than 0.001 in at least four donors. Filtered communication values were reshaped into feature-by-sample matrices. Interactions unavailable for a particular donor were retained as missing values rather than automatically being assigned a communication probability of zero. For each candidate interaction, a minimum of three non-missing donors was required in both the healthy and risk groups before differential analysis. Differential communication between *AHI1* risk and healthy groups was assessed using linear models implemented in limma v3.62.2.^140^ Because CellChat communication probabilities are bounded, values were transformed using the arcsine square-root transformation before model fitting, with input probabilities constrained to the interval [0,1]. Linear models were fitted using lmFit(), followed by empirical Bayes moderation using eBayes(trend = TRUE, robust = TRUE). Positive coefficients indicated higher inferred communication in risk samples, whereas negative coefficients indicated lower inferred communication. Nominal *p*-values were obtained from the moderated models, and correction for multiple hypothesis testing was performed using the Benjamini–Hochberg method.

### Gene regulatory network

To identify transcription factors (TFs) associated with target-gene regulation during human brain development, we analyzed snMultiome data using a motif-informed gene regulatory network (GRN) framework. Analyses were performed separately within developmental stage and broad cell type, including glial cells, excitatory and inhibitory neurons. Developmental periods were grouped into prenatal (periods 3–6) and postnatal (periods 8–15) stages. Single-nuclei profiles were aggregated into metacells generated upstream from the RNA modality within broad cell types. These metacell assignments were used as the shared unit for RNA and ATAC aggregation. Only batches containing both valid metacell mappings and corresponding ArchR Arrow files were retained for downstream analysis. For each target gene, candidate CREs were defined as ATAC peaks within ±100 kb of the TSS. Peak accessibility was quantified at the metacell level across all valid period-by-batch groups within the selected stage and cell type. Peaks were prioritized using stage-level accessibility metrics derived from the fraction of open metacells and mean accessibility across batches. Given the relatively broad gene-centered regulatory window (±100 kb around the TSS), stringent accessibility filtering was applied to reduce the inclusion of weakly supported candidate CREs. Specifically, peaks were retained if they exceeded a minimum median open-metacell fraction of 0.05 and had a stage-level accessibility score above the median. A maximum of 200 peaks per target gene was retained, when this limit was enabled.

Motif enrichment analysis was performed on retained peak sequences using Find Individual Motif Occurrences (FIMO) with JASPAR 2024,^162^ transcription factor motifs. Peaks containing significant motif matches (motif-hit *P* < 1×10^-4; scan-level threshold 0.05) were used to construct a binary TF-by-peak motif matrix, defining biologically plausible TF–CRE relationships. Because TFs may map to multiple peaks and peaks may contain motifs for multiple TFs, the resulting regulatory groups were overlapping. For each target gene, metacell-level TF expression (X), peak accessibility (Z), and target-gene expression (y) matrices were assembled across all valid stage-predominant batches. RNA and ATAC matrices were batch-corrected using reciprocal PCA (RPCA) and LSI-based integration, respectively. Period indicators were included as unpenalized covariates in downstream models, and the target gene was excluded from the TF predictor matrix prior to regression.

GRN inference was performed using a repeated screen-and-clean framework with overlapping group lasso regularization. Specifically, TF and CRE predictors were concatenated into a combined design matrix and overlapping TF–CRE groups were defined according to motif-supported TF–peak relationships. In each iteration, metacells were randomly partitioned into selection and inference subsets. Overlapping group lasso was fit on the selection subset to identify sparse TF and CRE predictors while respecting the motif-derived group structure. Ordinary least squares regression was then applied to the inference subset using only selected predictors, yielding post-selection coefficient estimates and *p*-values. Variables not selected in a given iteration were assigned a default *p*-value of 0.95. This procedure was repeated multiple times (typically 20–100 iterations), and *p*-values were aggregated across repetitions using the Cauchy combination test, followed by Benjamini–Hochberg multiple-testing correction. For each TF and CRE, we additionally summarized the mean coefficient, median coefficient, and selection frequency across repeated runs. Final GRN results reported in the manuscript correspond to TFs with significant adjusted *p*-values together with reproducible coefficient estimates across repeated split-sample regressions.

A locus-centered analysis was performed using the same statistical framework but with modified CRE selection criteria to accommodate the more restricted genomic context. Candidate CREs were limited to ATAC peaks within ±50 kb of externally provided variants of interest. Unlike the gene-centered GRN analysis, the locus-centered pipeline applied a more permissive accessibility threshold to retain potentially causal variant-proximal regulatory elements that may be accessible in only a small subset of metacells or batches. To constrain the number of candidate predictors, peaks were prioritized by genomic proximity to the locus of interest, and a maximum of 50 peaks was retained per locus. All subsequent preprocessing, motif matching, overlapping-group regression, and repeated split-sample inference procedures followed the same framework as the primary GRN pipeline. These differences in peak selection were intentional and reflected the distinct objectives of broad gene-centered regulatory inference and focused locus-centered analysis.

### CICERO co-accessibility analysis

Cell type– and developmental stage–specific co-accessibility networks were generated with CIC-ERO v1.29.2. Analysis groups were defined based on considering both cell type and development stage. Groups with fewer than 500 cells were excluded. For groups with more than 5,000 cells, 5,000 cells were randomly sampled without replacement using the same random seed.

For each group, a binarized peak-by-cell accessibility matrix generated from the ArchR Peak-Matrix was supplied to Monocle 2 as a CellDataSet. Cell coordinates for CICERO neighborhood construction were taken from the exported IterativeLSI embeddings. A two-dimensional UMAP was computed from these coordinates with uwot, using cosine distance, min_dist = 0.3, and n_neighbors set to the smaller of 30 or the number of cells minus one.

CICERO cell neighborhoods were constructed with k = 50 for the initial runs. The distance parameter was estimated separately for each group with estimate_distance_parameter, using 100 sampled windows, a 1-Mb window, and a 500-kb distance constraint. The mean of the finite estimated distance parameters was supplied to generate_cicero_models, and peak-pair co-accessibility scores were assembled with assemble_connections. Connections with absolute co-accessibility scores greater than 0.2 were retained. CCANs were generated from these retained connections using generate_ccans with a co-accessibility cutoff of 0.2.

To account for differences in cell number, four groups were rerun using all available cells and a neighborhood parameter scaled to approximately 1% of the group size, with a minimum k of 4: Micro_prenatal (n = 571, k = 10), SMC_postnatal (n = 1,089, k = 25), Immune_postnatal (n = 3,430, k = 34), and VLMC_postnatal (n = 3,917, k = 39). These reruns replaced the initial k = 50 results for those groups; the remaining ten groups retained k = 50. Due to insufficient data quality in these groups, they were excluded from further analysis.

### Prioritization of candidate cis-regulatory elements for CRISPR activation

We focused on haploinsufficient NDD genes identified by McDiarmid et al.^53^ to systematically prioritize candidate cis regulatory elements for CRISPRa. For 164 NDD genes, we generated a comprehensive catalog of putative cREs within 500 kb upstream and downstream of each gene’s TSS, yielding approximately 75 candidate elements per gene. Each candidate cRE was annotated with multiple features associated with regulatory activity, including chromatin accessibility across developmental stages and cell types, overlap with differentially accessible regions, previously characterized brain enhancers,^37,54–56^ predicted regulatory elements,^57^ H3K27ac enrichment (EN-CODE: ENCSR220TRW),^36^ ABC scores,^53,58^ sequence conservation across species,^59^ Cicero coaccessibility, and chromatin accessibility in SH-SY5Y cells (GEO: GSM2700775).^60^ See also Key Resources Table and Table S24.

### Statistical analysis of CRISPRa-mediated SOX5 activation

The effects of CRISPRa targeting on *SOX5* mRNA and protein expression were evaluated across three independent experiments, each comprising three biological replicates per condition (nine biological replicates in total per condition). Expression levels were normalized to the corresponding non-targeting control within each experiment. For protein quantification, *SOX5* immunoblot signals were normalized to β-actin. Statistical significance was assessed using a linear model with experimental run included as a blocking factor to account for between-experiment variability.

The effects of CRISPRa targeting were evaluated relative to the non-targeting control. Fold changes were calculated for each experiment, and mean fold changes were summarized across the three independent experiments. Statistical results are provided in **Tables S25** and **S26**.

## ADDITIONAL RESOURCES

Description: BrainVar website is an interactive resource for exploring the atlas generated in this study and colocalization analysis. www.brainvar.org

Description: UCSC Cell Browser provides interactive visualization and exploration of our snMultiome datasets. brainvar.cells.ucsc.edu

Description: UCSC Genome Browser ATAC-seq tracks allow visualization of chromatin accessibility in different cell types and developmental stages. https://genome-test.gi.ucsc.edu/cgi-bin/hgTrackUi?db=hg38&g=singleCellSignalsPeaks

## Supplementary Documents

### Document S1. Structural variant discovery

**Table x.** Count of genes affected by SVs with different predicted consequences.

| Predicted Consequence of SVs | Cohort-wide |  | Averaged count per genome |  |
|---|---|---|---|---|
|  | #SV sites | #Genes | #SV sites | #Genes |
| pLoF | 604 | 859 | 37.3 | 52.0 |
| whole gene copy gain | 312 | 615 | 10.9 | 23.6 |
| intragenic exon duplication | 180 | 172 | 5.6 | 5.6 |
| partial exon duplication | 53 | 55 | 4.1 | 4.2 |
| 5' gene duplication | 254 | 274 | 8.9 | 9.3 |
| 3' gene duplication | 288 | 294 | 8.3 | 9.8 |
| INV span | 5 | 64 | 0.0 | 0.3 |
| intronic | 19827 | 7367 | 2460.9 | 1873.7 |
| UTR-overlap | 505 | 491 | 46.3 | 46.3 |
| promoter-overlap | 367 | 373 | 38.2 | 39.3 |

**Fig. 1.**
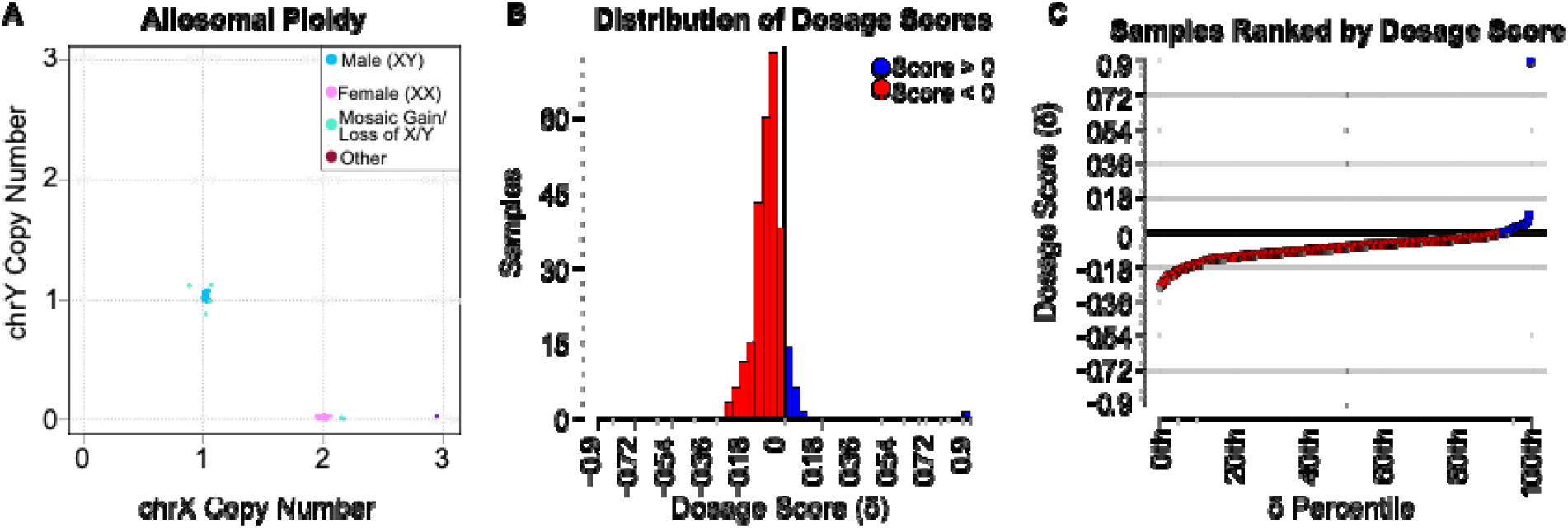
Ploidy and whole genome dosage bias (WGD) in BrainVar. a) Ploidy of sex chromosomes X and Y. b) Histogram of WGD. c) Distribution of WGS scores in ascending order among BrainVar samples.

**Fig. 2.**
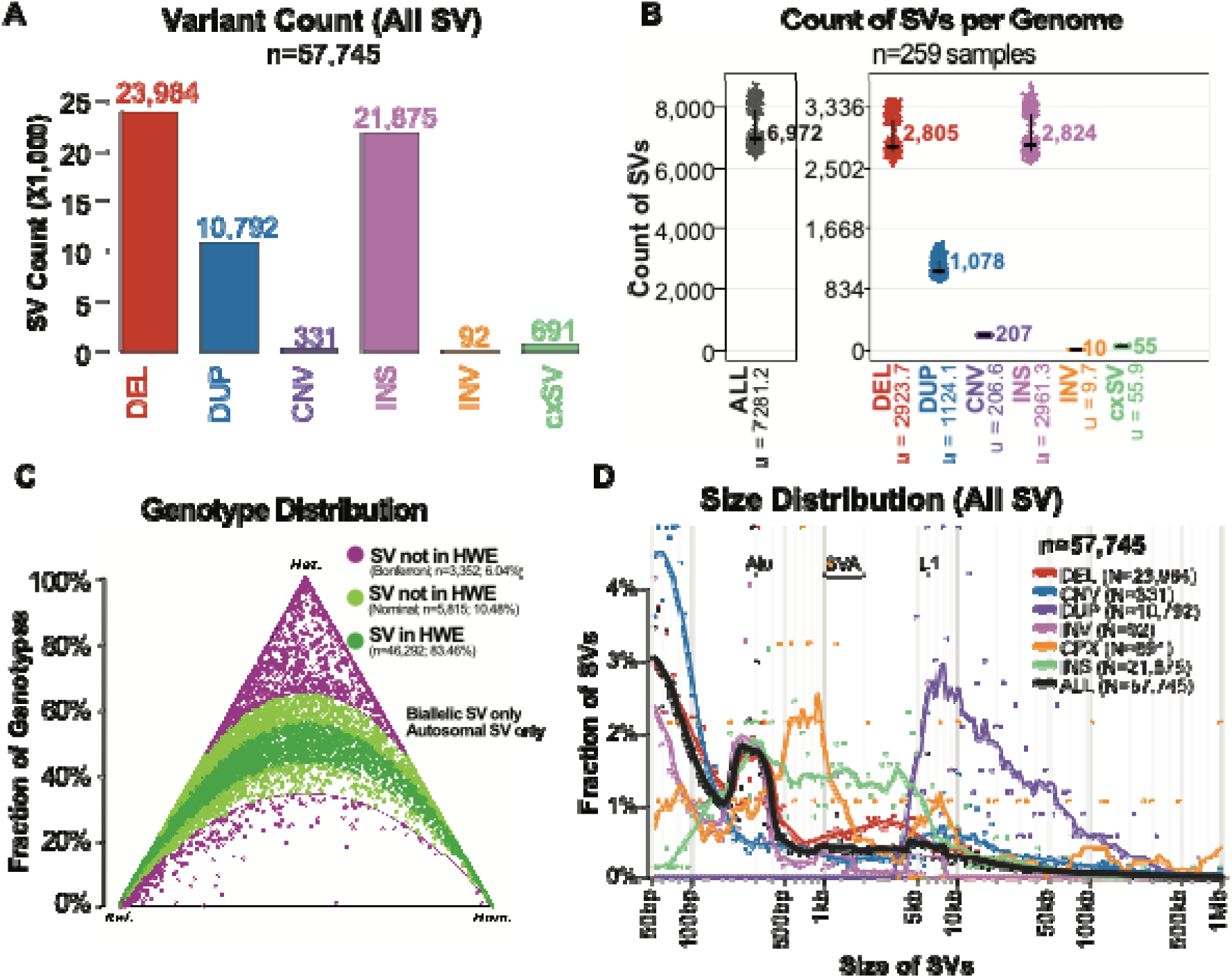
Distribution of SVs across the BrainVar dataset. A) Total counts of SVs stratified by class. B) Distribution of per-genome SV counts across BrainVar individuals. C) Hardy–Weinberg equilibrium assessment of autosomal biallelic SVs. D) Size distribution of SVs by class.

**Fig. 3.**
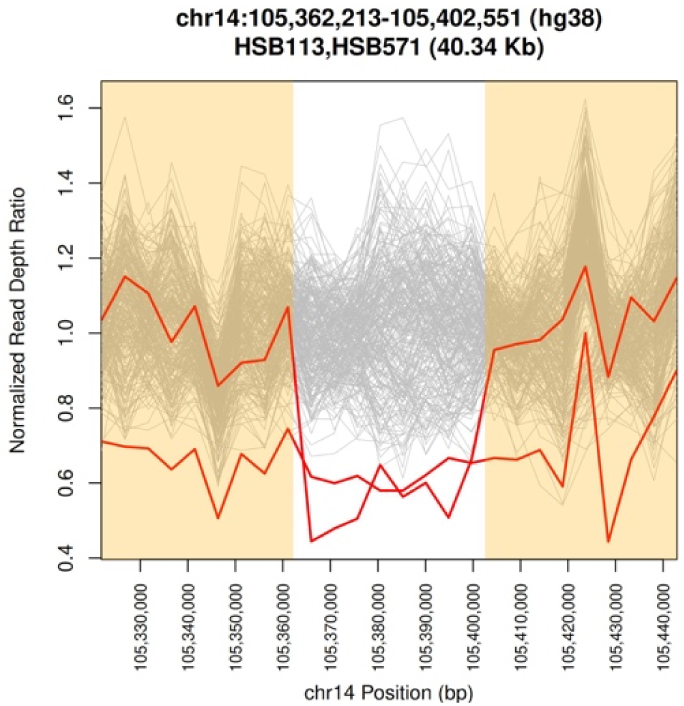
**Sequencing depth distribution across a 40 kb deletion encompassing the 3**′ end of *PACS2*

